# The proteasome maturation factor POMP moonlights as a stress-induced transcriptional regulator

**DOI:** 10.1101/2025.04.25.650603

**Authors:** Stefano L. Giandomenico, Mara Mueller, Marc van Oostrum, Kristina Desch, Bastian Krause, Georgi Tushev, Elena Ciirdaeva, Julian D. Langer, Beatriz Alvarez-Castelao, Erin M. Schuman

## Abstract

Proteostasis - the maintenance of proteins at proper concentrations, conformations, and subcellular locations - is essential for cellular function and is governed by tightly regulated protein synthesis and degradation pathways. The Proteasome Maturation Protein (POMP) is a key chaperone involved in assembling the proteasome, the primary complex responsible for protein degradation. Despite the conserved role of POMP, its loss produces contrasting proteostatic effects in yeast and mammalian cells, pointing to additional, unexplored functions. In this study, we investigated the possibility that POMP plays a moonlighting role in proteostasis regulation. We discovered that upon proteasome disruption, POMP rapidly accumulates in the nucleolus in a manner dependent on HSF1 and reactive oxygen species (ROS). Proteomic analysis of POMP interactors revealed RNA processing factors and transcriptomic profiling showed that nucleolar POMP orchestrates a protective transcriptional program. Our findings indicate POMP is a built-in sensor and effector within the proteasome assembly pathway, capable of buffering disturbances in proteasome function through a novel, non-canonical role. Notably, this mechanism is developmentally controlled and active in neurodegenerative disease contexts.

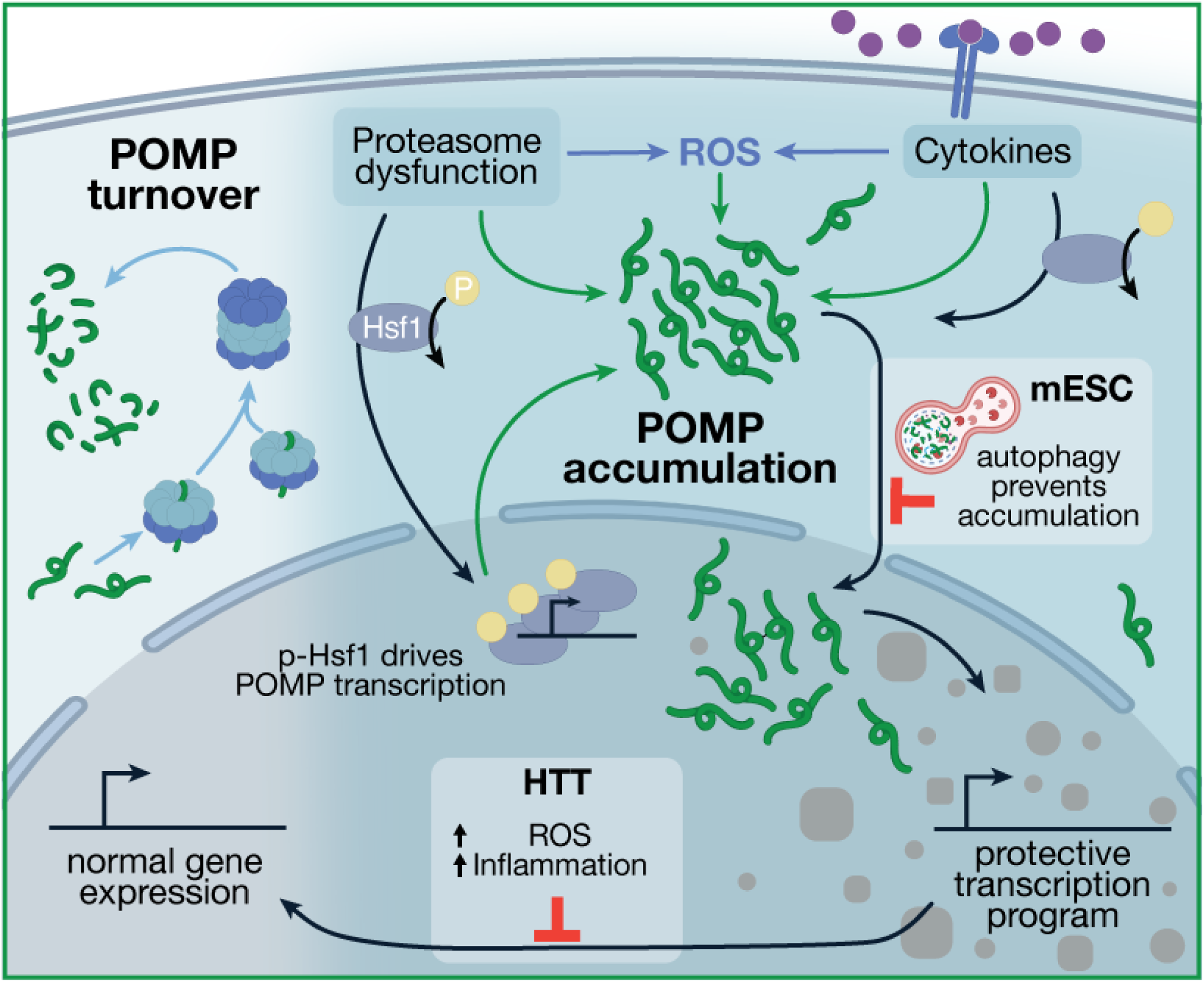

## Introduction

Proteostasis is vital for organismal life. The cytoplasm and ER serve as the primary sites for protein synthesis, quality control, and maturation where ribosomes, molecular chaperones, the ubiquitin-proteasome system, and the lysosome work together to meet cellular demands^1–3^. These complex molecular machines are central to healthy cell physiology and homeostatic mechanisms have evolved around them to ensure rapid and resilient responses to stressors^4–6^. Proteostatic perturbations, sensed in the cytoplasm and ER, initiate signaling cascades that reprogram the cell transcriptome in the nucleus, enabling adaptive responses for cellular homeostasis and survival ^6,7^. For example, when germline stem cells differentiate there is an increase in ribosome biogenesis and protein synthesis^8,9^. Similarly, to overcome proteasome dysfunction cells are able to mount compensatory feedback mechanisms, mediated by the transcription factors Rpn4^10^ (in yeast) and TCF11/NRF1 (in mammals), that initiate in the cytoplasm and at the ER, respectively, and converge in the nucleus ^11,12^. Within the nucleus, nuclear condensates, and in particular the nucleolus, are central homeostatic hubs that sense cellular insults to initiate and coordinate cellular stress responses^13–15^. These nuclear compartments are membrane-less and are formed by liquid-liquid phase separation, which makes them particularly susceptible to changes in the chemical environment^16,17^. The dynamic relocalisation of oncogenes (e.g. c-Jun and c-Myc) and tumor-suppressors (e.g. p53) between the nucleoplasm and nuclear condensates is a deciding factor in cellular life and death decisions^18–21^. In fact, if a stressor is significant and prolonged, cells must decide whether to mount a compensatory response or initiate apoptosis^6,22^. How and where these decisions are made and which factors integrate and buffer pro-apoptotic and pro-survival signals remain poorly understood.

To investigate mechanisms of proteostasis, we focused on the proteasome, a 2.5 MDa complex comprising a 20S core particle (CP) and 19S regulatory particles (RPs) that work together to recognize, unfold, and proteolytically degrade ubiquitin-tagged proteins^23,24^. The CP, composed of four heptameric rings (two outer α-rings and two inner β-rings), has a cylindrical structure with a central proteolytic chamber^23,24^. In addition to the standard proteasome, cells can assemble specialized proteasomes such as the immunoproteasome, which incorporates alternative catalytic subunits (PSMB8, PSMB9, and PSMB10) and plays a key role in antigen processing for immune surveillance ^25^. Assembly of the standard CP involves 14 subunits (PSMA1-7 and PSMB1-7) and 5 chaperones, PSMG1-4 and the proteasome maturation protein, POMP^23,26^. POMP, the mammalian homolog of yeast Ump1, coordinates the assembly of the β-ring and is degraded upon CP maturation ^27–29^(Figure 1a). POMP protein levels, therefore, could serve as a sensor of proteasome assembly and activity. Despite their apparent functional homology, Ump1 and POMP show limited sequence homology and their loss leads to different outcomes in yeast and mammals^28–30^. Yeast can overcome Ump1 loss through Rpn4-driven upregulation of proteasome subunits^10,26,30–32^, while mammalian cells cannot^33–35^, despite their ability to upregulate proteasome subunits via TCF11/NRF1 in response to pharmacological inhibition of the proteasome^11,12,36^. POMP overexpression can induce *de novo* proteasome synthesis, suggesting additional roles in regulating proteasome levels^37^. However, at present the mechanisms underlying these differences and POMP’s potential additional functions remain unknown.

**Figure 1.**
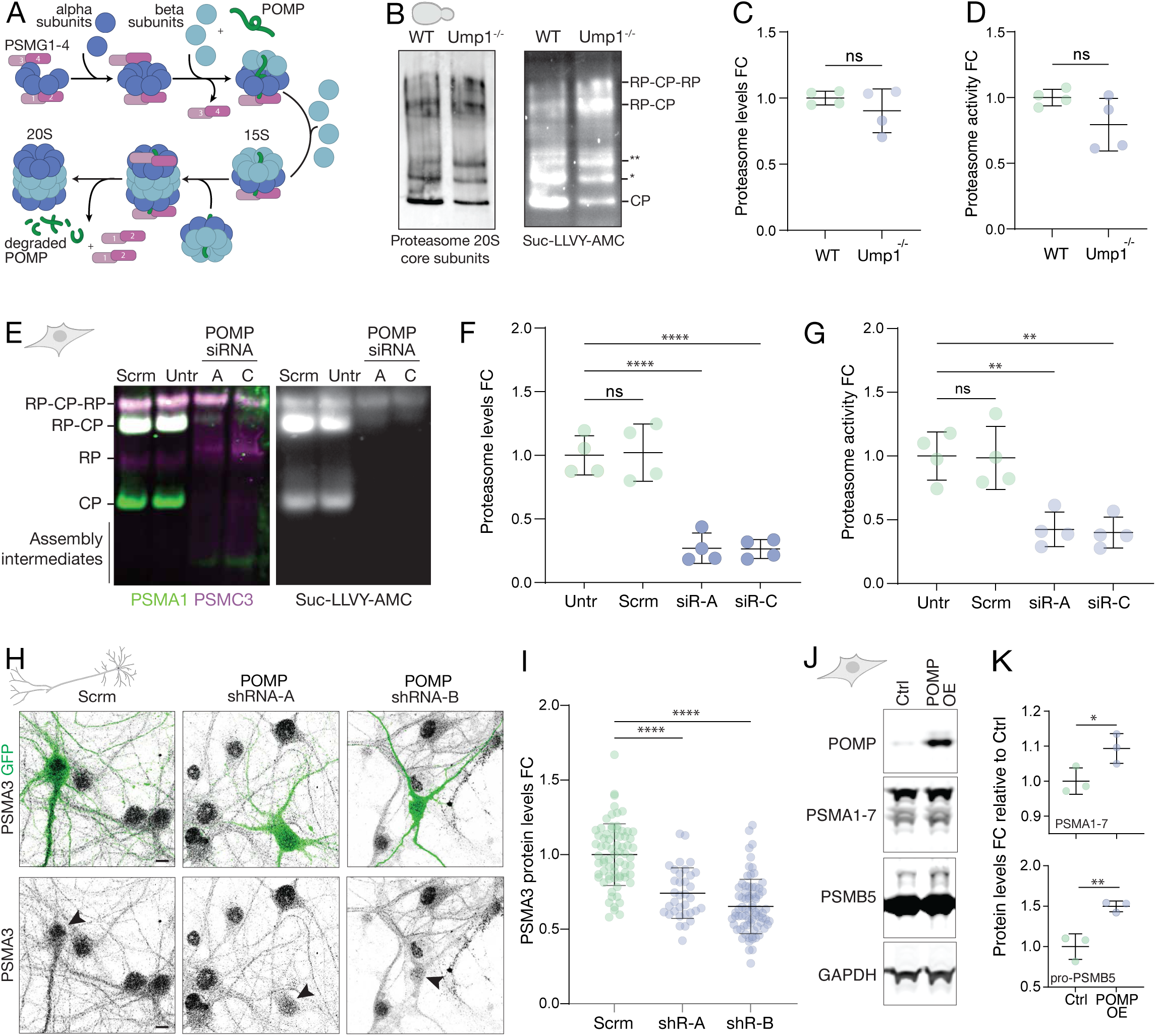
Discrepant responses to Ump1/POMP deletion in yeast and mammalian cells. (A) Scheme of proteasome biogenesis depicting the assembly of the catalytic core rings comprising alpha and beta subunits as well as the chaperones PSMG1-4 and POMP (Ump1). POMP gets degraded by the proteasome when the two hemi-proteasomes come together to form the nascent 20S proteasome. (B) Representative native gel electrophoresis labelled with an anti-PMSA1-7 antibody (left) and a proteasome fluorogenic substrate (Suc-LLVY-AMC; right) in wild-type (WT) and Ump1^-/-^ yeast strains. Native gel showing the different assembly states of the proteasome, including the 20S (CP, core particle), singly capped (RP-CP) and doubly capped (RP-CP-RP) as well as * and ** representing the CP-Blm10 and CP-Blm10_2_ alternative proteasome configurations, respectively. (C) Analysis of proteasome levels from yeast native gel experiments. Deletion of UMP1 did not lead to a significant change in the levels of the proteasome (ROI measured from gels spanned from CP to RP-CP-RP). ns=p>0.05, unpaired two-tailed t-test, n=4, mean±SD, FC=fold change. (D) Analysis of proteasome activity levels from yeast native gel experiments. Deletion of UMP1 did not lead to a significant change in the activity level of the proteasome (ROI measured from gels spanned from CP to RP-CP-RP). ns=p>0.05, unpaired two-tailed t-test, n=4, mean±SD, FC=fold change. (E) Representative native gel electrophoresis labelled with an anti-PMSA1 (green) and anti-PMSC3 (magenta) antibody (left) and a fluorescent proteasome activity reporter (Suc-LLVY-AMC; right) in control HEK293 cells (Untr) and cells transfected with a scrambled (Scr) or 1 of 2 different POMP siRNAs (A or C). In the native gel the different assembly states of the proteasome, including the 20S (CP, core particle), singly capped (RP-CP) and doubly capped (RP-CP-RP) as well as assembly intermediates are shown. (F) Analysis of proteasome levels from HEK293 cell native gel experiments. Knockdown of POMP by either siRNA led to a significant reduction in the levels of the proteasome while the scrambled construct had no effect (ROI measured from gels spanned from CP to RP-CP-RP). FC = fold change. ns=p>0.05, ****p≤0.0001, one-way ANOVA and post-hoc Dunnett’s multiple comparisons test, n=4, mean±SD, FC=fold change. (G) Analysis of proteasome activity levels from HEK293 cell native gel experiments. Knockdown of POMP by either siRNA led to a significant reduction in the activity level of the proteasome while the scrambled construct had no effect (ROI measured from gels spanned from CP to RP-CP-RP). FC = fold change. ns=p>0.05, **p≤0.01, one-way ANOVA and post-hoc Dunnett’s multiple comparisons test, n =4, mean±SD. (H) Representative images of primary rat hippocampal neurons (DIV 21) transfected with either a scrambled (Scrm) or one of two different POMP shRNAs (shRNA-A or shRNA-B) and then immunostained with an anti-PSMA3 antibody (black signal) to estimate the 20S proteasome abundance. Transfected neurons expressed GFP and are indicated by the arrowheads in the lower row of images. Scale bar=10 µm. (I) Analysis of PSMA3 proteasome levels from the neuron imaging experiment shown in H. Knockdown of POMP by either shRNA led to a significant reduction in the levels of PSMA3 relative to the scrambled control. ****p≤0.0001, one-way ANOVA and post-hoc Dunnett’s multiple comparisons test, n=71 (Scrm), 34 (A) and 74 (B) neurons from 3 biological replicates, mean±SD, FC=fold change. (J) Representative Western Blot analysis from HEK293 cells transfected with an empty vector (Ctrl) or a POMP-overexpression construct (POMP-OE) assessing the expression of POMP and the proteasome 20S alpha (anti-PSMA1-7) or beta (anti-PSMB5) subunits, as well as a loading control GAPDH. (K) Analysis of the Western blot experiments as shown in J. Shown are the PSMA1-7 protein levels or the pro-PSMB5 levels (upper band in blot in J), relative to control. GAPDH was used as a loading control. *p<0.05, **p<0.01, unpaired two-tailed t-test, n = 3, mean±SD, FC=fold change.

Here we discovered that POMP has a moonlighting function that is invoked during cellular stress, including proteasome inhibition. Under normal conditions, POMP localizes primarily to secretory vesicles, rather than the ER. When proteasome function is compromised, POMP relocalizes from cytosolic vesicles to the nucleolus and nuclear bodies. This process requires HSF1-dependent transcription and increased cellular reactive oxygen species (ROS), resulting in a shift in POMP’s interactome towards RNA binding proteins. In the nucleus, POMP induces a protective transcriptional signature that preserves ribosome and proteasome biogenesis while inhibiting apoptosis initiation. This pathway can be activated by cytokine signaling and shows differential regulation across cell types, developmental stages, and disease states.

## Results

To investigate Ump1/POMP’s discrepant responses in yeast and mammalian cells, we first used WT and Ump1^-/-^ yeast strains (Figure S1a-c) and performed native gel analysis to assess both the expression and activity levels of the assembled proteasome (Figure 1b-d and Figure S1d). As observed by others^27,30–32^ we found that the abundance of the different proteasome assembly states, including the singly (26S) and doubly (30S)-capped proteasome, was not significantly altered by the absence of Ump1 (Figure 1b,c and Figure S1d). Moreover, proteasome activity, monitored using the fluorogenic substrate Suc-LLVY-AMC^38^, was also not significantly altered by Ump1 deletion (Figure 1b,d). Since the complete loss of POMP is lethal in mammalian cells^33,39^, we used siRNAs to knock-down (KD) POMP in HEK293 cells (Figure S1e,f). We again performed native gel analysis and found (as previously described^33,34^) that POMP KD led to a significant decrease in both the assembled proteasome and its activity levels, as well as the accumulation of assembly intermediates (Figure 1e-g and Figure S1g,h). To verify that the proteasome sensitivity to POMP KD is a general feature of mammalian cells, we conducted similar experiments in neurons. Primary rat hippocampal neurons transfected with POMP shRNAs (Figure S1i,j) exhibited a significant reduction in PSMA3 protein levels (Figure 1h,i), indicating that the failure to upregulate proteasome subunit levels in response to a loss of POMP is a common feature of mammalian cells. To examine whether the mammalian POMP-sensitive regulation of proteasome levels is bi-directional, we overexpressed POMP in HEK293 and conducted Western blot analysis to measure proteasome subunit levels. We observed a significant increase in the levels of PSMA1-7 and immature (pro-PSMB5) subunits (Figure 1j,k), confirming that POMP overexpression is sufficient to increase proteasome subunit levels^37^. Taken together these data indicate that yeast and mammalian cells exhibit contrasting proteasome responses when the proteasome biogenesis factor Ump1/POMP is reduced.

To gain more insight into the individual protein changes that occur in response to loss of Ump1/POMP, we performed mass spectrometry (MS)-based proteomic analysis. Strikingly, while the Ump1^-/-^ yeast strain displayed a significant increase in proteasome core particle subunits (Figure 2a), POMP KD in mammalian (HEK293) cells, by contrast, led to a significant decrease in proteasome core particle subunits (Figure 2b). With this in mind, it was surprising to observe that, despite the contrasting proteostatic responses, both yeast and HEK293 cells showed a significant upregulation in the transcription factors Rpn4 and TCF11/NRF1 (hereafter NRF1), respectively, in response to loss of Ump1/POMP (Figure 2c,d and Figure S2a,b). As mentioned above, NRF1 drives the transcription of proteasome genes in response to prolonged proteasome inhibition^11,12^. To examine POMP’s role in this pathway, we measured POMP levels after treatment with three different proteasome inhibitors (Carfilzomib, Epoxomicin, MG132; 5 hrs) and found that both POMP mRNA and protein were increased (Figure 2e-g). To gain temporal resolution and determine which molecular player may be the driver, we monitored the expression levels of both POMP and NRF1 following 1, 2.5 or 5 hrs of proteasome inhibition. While POMP levels quickly rose to their maximum at 1h and then plateaued, NRF1 exhibited a robust and steady increase over the 5h period (Figure 2h,i). Over the same time course, surprisingly, there was no change in proteasome subunits (PSMA1-7 and PSMB5) (Figure 2h,i and Figure S2c,d). The difference in the accumulation kinetics of POMP and NRF1 does not support the simple idea that NRF1 is responsible for the observed rapid proteasome inhibitor-induced increase in POMP. In addition, the absence of a concomitant change in proteasome subunits suggests a potential proteasome-independent function of POMP. To test directly the role of NRF1 in the upregulation of POMP, we knocked it down using two different siRNAs (Figure 2j) and measured changes in POMP protein levels following MG132 treatment. Although the typical accumulation of NRF1 in response to MG132 treatment was strongly reduced by knockdown, POMP induction was unaffected (Figure 2j,k), indicating that NRF1 does not promote the early accumulation of POMP.

**Figure 2.**
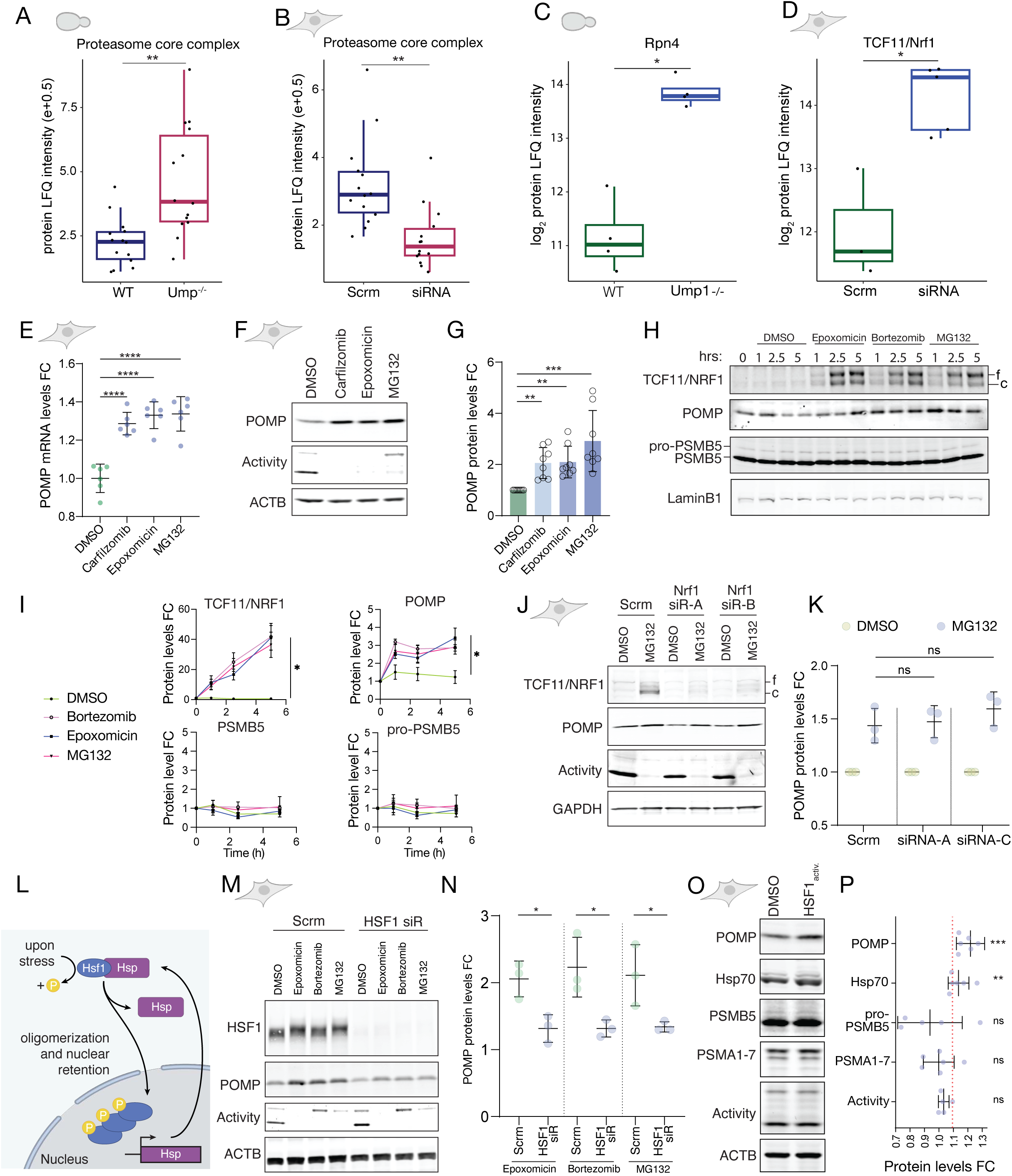
Proteasome inhibition leads to an HSF1-dependent early increase in POMP mRNA and protein levels. (A) Analysis of proteasome core complex (α1-7 and β1-7) protein label-free quantification (LFQ) intensity values from mass spectrometry experiments in WT and Ump1^-/-^ yeast strains. The proteasome core complex proteins were significantly upregulated in the absence of Ump1^-/-^. **p≤0.01, Welch two-tailed t-test. Boxplots show the median (line), interquartile range (box), and 1.5×IQR whiskers. Individual data points shown represent the mean intensity of the individual CP components. (B) Analysis of proteasome core complex (PSMA1-7 and PSMB1-7) protein label-free quantification (LFQ) intensity values from mass spectrometry experiments in HEK293 cells treated with a POMP siRNA or a scrambled control siRNA (Scrm). The proteasome core complex proteins were significantly downregulated following POMP knockdown. Individual data points shown represent the mean intensity of the individual CP components. **p≤0.01, Welch two-tailed t-test. Boxplots show the median (line), interquartile range (box), and 1.5×IQR whiskers. Individual data points shown represent the mean intensity of the individual CP components. (C) Analysis of the transcription factor Rpn4 protein label-free quantification (LFQ) log_2_-scaled intensity values from mass spectrometry experiments in WT and Ump1^-/-^ yeast strains. Rpn4 was significantly upregulated in the absence of Ump1^-/-^. *p≤0.05, two-tailed Wilcoxon rank-sum test, n=4. Boxplots show the median (line), interquartile range (box), and 1.5×IQR whiskers. Individual data points shown represent biological replicates. (D) Analysis of Nrf1/TCF11 protein label-free quantification (LFQ) log_2_-scaled intensity values from mass spectrometry experiments in HEK293 cells treated with a POMP siRNA or a scrambled control siRNA (Scrm). Nrf1/TCF11 protein was significantly upregulated following POMP knockdown. *p≤0.05, two-tailed Wilcoxon rank-sum test, n=3 (Scrm) and 5 (POMP siRNA). Boxplots show the median (line), interquartile range (box), and 1.5×IQR whiskers. Individual data points shown represent biological replicates. (E) POMP mRNA levels in HEK293 cells following 5 hr treatment with each of three different proteasome inhibitors (Carfilzomib, Epoxomicin and MG132) or DMSO. ****p≤ 0.0001, one-way ANOVA and post-hoc Dunnett’s multiple comparisons test, n=6, mean±SD, FC=fold change. (F) Representative Western Blot analysis from HEK293 cells treated for 5 hr with each of three different proteasome inhibitors (Carfilzomib, Epoxomycin or MG132) or DMSO, assessing proteasome activity, POMP and ACTB loading control levels. (G) Analysis of the Western blot experiments as shown in F. Shown are POMP protein levels relative to the DMSO control. ACTB was used as a loading control. ** = p<0.01, ***p≤0.001, RM one-way ANOVA and post-hoc Dunnett’s multiple comparisons test on non-normalised data, n=8, mean±SD, FC=fold change. (H) Representative Western blot time course from HEK293 cells treated with each of three different proteasome inhibitors (Epoxomicin, Bortezomib, or MG132) or DMSO for 1, 2.5 and 5 hr assessing the levels of TCF11/NRF1, POMP, PSMB5 and, as a loading control, LaminB1. (I) Analysis of the Western blot time course as shown in H. Reported are LaminB1-normalised protein level fold changes relative to T = 0 hr in response to treatment with DMSO, Epoxomycin, Bortexomib, or MG132 for POMP, TCF11/NRF1, PSMB5 and its precursor form pro-PSMB5 (upper band). *p ≤0.05, one-way ANOVA and post-hoc Dunnett’s multiple comparisons test on the 5 h time point, n=4, mean±SEM, FC=fold change. (J) Representative Western Blot analysis of HEK293 cells transfected with two different siRNAs against TCF11/NRF1 or Scrm control, and at 72 hr post-transfection treated with the proteasome inhibitor MG132 for 5 hr. Shown are proteasome activity levels and protein levels for TCF11/NRF1, POMP and the loading control GAPDH. (K) Analysis of the Western blot experiment as shown in J. Reported are POMP protein level changes in response to 5 hr MG132 treatment in TCF11/NRF1 knock-down and Scrm control cells. The early induction of POMP protein is not affected by TCF11/NRF1 knock-down. ns=p > 0.05, one-way ANOVA and post-hoc Dunnett’s multiple comparisons test, n=3, mean±SD, FC=fold change. (L) Schematic illustration of the HSF1 transcription factor activation mechanism. Under basal conditions, HSF1 is kept inactive in the cytoplasm by interaction with heat-shock proteins (Hsps). In response to stress HSF1 gets phosphorylated, released by the Hsps and translocates into the nucleus to form transcriptionally-competent trimers that activate the stress response. (M) Representative Western Blot analysis of HEK293 cells transfected with one siRNA against HSF1 or Scrm control, and at 72 hr post-transfection treated with three different proteasome inhibitors (Epoxomicin, Bortezomib, MG132) for 5 hr. Shown are proteasome activity and protein levels for HSF1, POMP and ACTB as a loading control. (N) Analysis of the Western blot experiment as shown in M. Shown are ACTB-normalised POMP protein level fold changes in response to a 5 hr treatment with Epoxomicin, Bortezomib or MG132 in HSF1 knock-down and Scrm control cells. HSF1 knock-down significantly reduced POMP induction in response to proteasome inhibition. *p≤0.05, unpaired two-tailed t-tests, n=3, mean±SD, FC=fold change. (O) Representative Western blot analysis of HEK293 cells treated with DMSO or the HSF1 activator HSF1B for 7 hr. Shown are proteasome activity levels (Activity) and protein levels for POMP, Hsp70, PSMB5, PSMA1-7, and ACTB. (P) Analysis of the Western blot experiment as shown in O. Shown are ACTB-normalised protein level fold changes in response to treatment with the HSF1 activator. HSF1 activation significantly elevated both POMP and Hsp70 levels, but did not significantly affect proteasome levels.**p≤0.01, ***p≤0.001, paired two-tailed t-tests, n=7 (POMP), 6 (Hsp70, PSMA1-7, Activity), 5 (pro-PSMB5), mean±SD, FC=fold change.

To identify the transcription factor that drives early POMP transcription in response to proteasome inhibition, we surveyed the POMP promoter region for *cis*-regulatory elements recognized by stress-response genes. Near the POMP promoter start site we found three HSF1 binding sites (Figure S2e). In response to proteotoxic stress HSF1 is phosphorylated resulting in activated trimers that mediate a primary transcriptional response to proteotoxic stress^40,41^ (Figure 2l). We examined whether proteasome inhibition leads to HSF1 activation and induction of its downstream target Hsp70^42,43^. Treatment of HEK293 cells with three different proteasome inhibitors led to a strong induction of HSF1 phosphorylation and a significant increase in Hsp70 (Figure S2f,g). To probe HSF1’s role in regulating POMP levels, we knocked-down HSF1 and measured both the basal and proteasome-sensitive levels of POMP. Depletion of HSF1 led to a decrease in basal POMP levels (Figure S2h,i) as well a significant reduction in the level of POMP induction by proteasome inhibitors (Figure 2m,n). A similar pattern of results was obtained with an HSF1 inhibitor^44^ (Figure Sup 2j-l). Lastly, we tested whether HSF1 activation alone is sufficient to upregulate POMP levels. Treatment of HEK293 cells with the activating compound HSF1B (HSF1_activ._; analogue of HSF1A^45^) elicited a significant increase in Hsp70 and POMP protein levels without affecting proteasome levels and activity (Figure 2o,p). Altogether, these data indicate that proteasome inhibition leads to a rapid, HSF1-dependent increase in POMP levels without concomitant changes in its partner proteasome subunits.

The above experiments prompted us to consider whether POMP serves an alternative role in the early response to cellular stress. To better understand the cellular sites where POMP localizes under normal and proteostatically-challenging conditions, we visualized it in cells using immunofluorescence. Previous reports on POMP localization in mammalian cells are somewhat contradictory, including one description of its localization to cytoplasm and nuclear bodies^46^, and another describing POMP as an ER-associated peripheral membrane protein^47^. To explore this further, we validated an antibody for immunofluorescence (Figure Sup 3A a,b) and stained both HEK293 cells and primary neurons for POMP and the ER marker Calnexin. POMP appeared as discrete puncta with cells showing only a modest colocalization with the ER (Figure 3a, b). To verify that we could indeed detect *bona fide* colocalization at the ER, we immunostained HEK293 and primary neurons for two ER resident proteins, Calnexin and Calreticulin; this time we observed an almost perfect colocalization in both cell types (Figure S3A c,d). To identify with which cellular structures POMP interacts we performed subcellular fractionation by differential centrifugation using neuronal whole cell extracts (Figure 3c). We recovered four fractions and performed MS-based proteomic profiling to confirm their quality and identify marker proteins for Western blot validation (Figure 3d,e; Figure S3A e,f). The four fractions corresponded to the following: fraction I-nuclear, plasma membrane, synapse, mitochondria and membrane protein complex components; fraction II-mitochondria and organellar membranes; fraction III-vesicle, secretory granule and proteasome components; and fraction IV-cytoplasm (Figure S3A e). Like the (immature form of the) proteasome subunit PSMB5, POMP was enriched in fraction III, a vesicular fraction (Figure 3d,e). To validate this localization, we performed immunolabeling and assessed the colocalization of POMP and the vesicle-associated vacuolar-type ATPase subunit ATP6V1H and the autophagosomal protein ATG5 (Figure 3f-i). We noted that the co-localization of POMP with these markers was much higher (Figure 3g, i) than that observed for the ER markers (Figure 3b).

**Figure 3.**
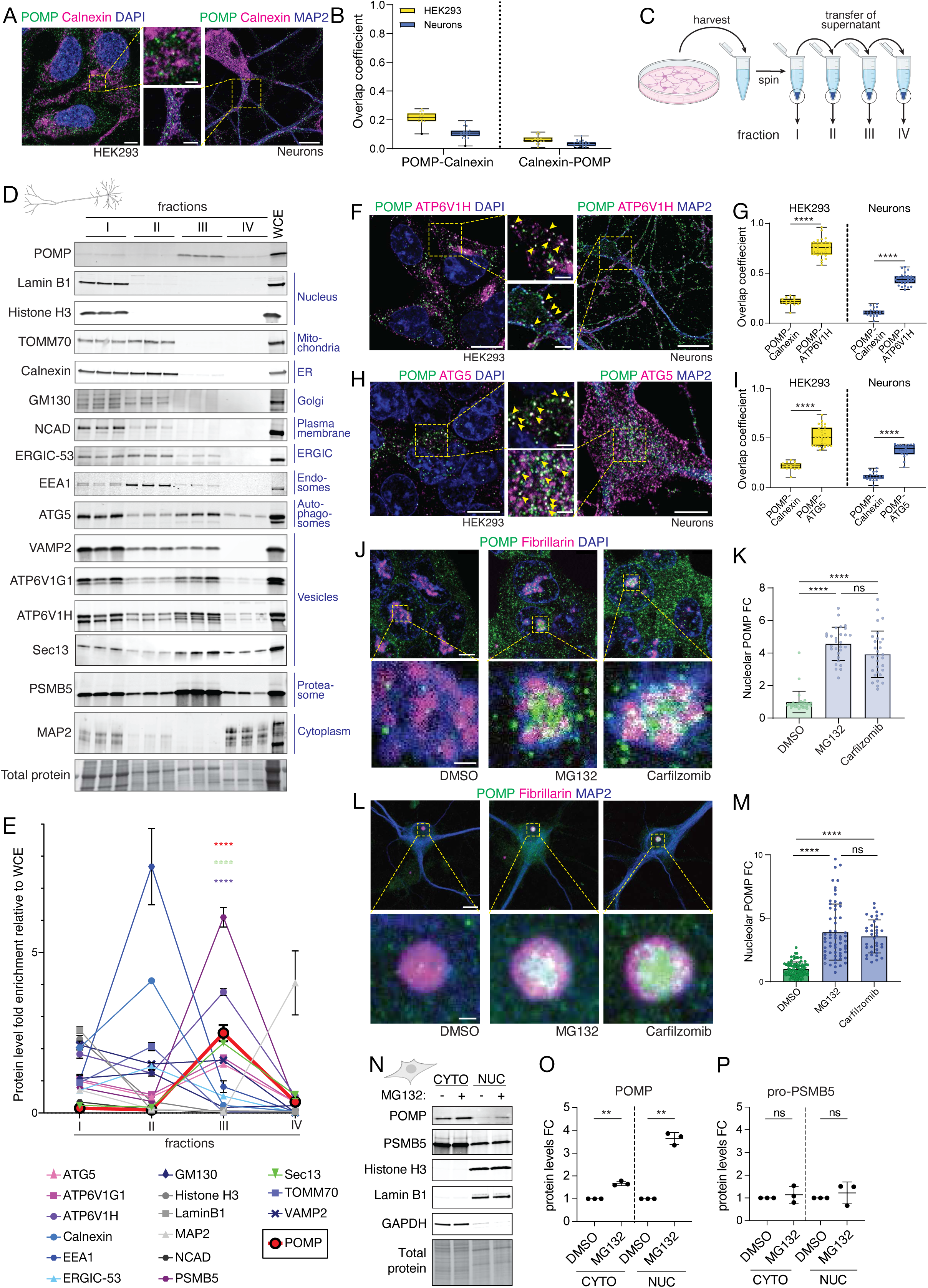
Under basal conditions, POMP localizes with the proteasome to vesicles but re-localizes to the nucleolus-without the proteasome-following proteasome inhibition. (A) Representative immunolabeling images from either HEK293 cells (left) or hippocampal neurons (right) showing labelling for POMP (green), the ER marker Calnexin (magenta) and either DAPI or the microtubule marker MAP2 (blue). Respective zoom-ins show the relatively low colocalization of POMP and Calnexin in both cell types. Scale bars, left and right = 5 and 10 µm, respectively, 1 µm in the insets. (B) Mander’s overlap coefficient analysis of immunolabeling experiments like that shown in A. On the left, the number of POMP particles that overlap with Calnexin, on the right, the number of Calnexin particles that overlap with POMP in HEK293 cells (yellow) or neurons (blue). (C) Scheme of neuronal subcellular fractionation experiments (see methods) conducted to determine the subcellular compartment(s) where POMP resides. (D) Western blot analysis of subcellular fractions derived from cortical neurons. Specific antibodies were used to detect the proteins indicated on the left, used as markers for the compartments indicated in blue on the right. WCE = whole cell extract. POMP was most abundant in Fraction III, a fraction characterized by the enrichment of multiple vesicular markers. (E) Analysis of Western blot fractionation experiments like that shown in D. The abundance of each marker protein, normalized to the total protein in its respective fraction, was quantified relative to its abundance in the whole-cell extract (WCE). POMP was significantly elevated in Fraction 3, along with several vesicular marker proteins like ATP6V1H and Sec13. ****p≤ 0.0001, p, one-way ANOVA and post-hoc Tukey’s multiple comparisons test, n=3, mean±SEM. (F) Representative immunolabeling images from either HEK293 cells (left) or hippocampal neurons (right) showing labelling for POMP (green), the vesicular marker ATP6V1H (magenta) and either DAPI or the microtubule marker MAP2 (blue). The yellow arrowheads in the zoom-ins show the clear colocalization of POMP and ATP6V1H in both cell types. Scale bars, left and right = 10 µm, 1 µm in the insets. (G) Mander’s overlap coefficient analysis of immunolabeling experiments like that shown in F. On the left, the number of POMP particles that overlap with ATP6V1H (plotted with the Calnexin data for comparison) in HEK293 cells (yellow) or neurons (blue). In both cell types, the POMP overlap with ATP6V1H was significantly greater than with Calnexin. ****p≤ 0.0001, one-way ANOVA and post-hoc Dunnett’s multiple comparisons test, n=14 (HEK293, POMP-Calnexin), 27 (HEK293, POMP-ATP6V1H), 20 (Neurons, POMP-Calnexin) and 27 (Neurons, POMP-ATP6V1H). Boxplots show the median (line), interquartile range (box), and Min-Max whiskers. (H) Representative immunolabeling images from either HEK293 cells (left) or hippocampal neurons (right) showing labelling for POMP (green), the vesicular marker ATG5 (magenta) and either DAPI or the microtubule marker MAP2 (blue). The yellow arrowheads in the zoom-ins show the clear colocalization of POMP and ATG5 in both cell types. Scale bars, left and right = 10 and 5 µm, respectively, 1 µm in the insets. (I) Mander’s overlap coefficient analysis of immunolabeling experiments like that shown in H. On the left, the number of POMP particles that overlap with ATG5 (plotted with the Calnexin data for comparison) in HEK293 cells (yellow) or neurons (blue). In both cell types, the POMP overlap with ATG5 was significantly greater than with Calnexin. ****p≤0.0001, one-way ANOVA and post-hoc Dunnett’s multiple comparisons test, n=14 (HEK293, POMP-Calnexin), 29 (HEK293, POMP-ATG5), 20 (Neurons, POMP-Calnexin) and 19 (Neurons, POMP-ATG5). Boxplots show the median (line), interquartile range (box), and Min-Max whiskers. (J) Representative immunolabeling images from HEK293 cells showing labelling for POMP (green), the nucleolar marker Fibrillarin (magenta) and DAPI (blue) under control conditions (DMSO) or after 5 hr treatment with two different proteasome inhibitors, MG132 or Carfilzomib. The insets show the clear increase in colocalization of POMP and Fibrillarin following proteasome inhibition. Scale bars, overviews and insets = 5 and 1 µm, respectively. (K) Analysis of immunolabeling experiments like that shown in J. Shown is the nucleolar POMP level fold change relative to control. Both proteasome inhibitors induced a significant increase in nucleolar POMP levels. ****p≤0.0001, one-way ANOVA and post-hoc Tukey’s multiple comparisons test, n= 30 (DMSO, Carfilzomib) and 29 (MG132), mean±SD, FC=fold change. (L) Representative immunolabeling images from hippocampal neurons showing labelling for POMP (green), the nucleolar marker Fibrillarin (magenta) and MAP2 (blue) under control conditions (DMSO) or after 5 hr treatment with two different proteasome inhibitors, MG132 or Carfilzomib. The insets show the clear increase in colocalization of POMP and Fibrillarin following proteasome inhibition. Scale bars, overviews and insets = 10 and 1 µm, respectively. (M) Analysis of immunolabeling experiments like that shown in L. Shown is the nucleolar POMP level fold change relative to control. Both proteasome inhibitors induced a significant increase in nucleolar POMP levels. ****p≤0.0001, one-way ANOVA and post-hoc Tukey’s multiple comparisons test, n= 90 (DMSO), 62 (MG132) and 37 (Carfilzomib), mean±SD, FC=fold change. (N) Representative Western blot analysis from a cytoplasmic-nuclear fractionation experiment conducted in HEK293 cells incubated in the presence or absence of the proteasome inhibitor MG132 for 5 hrs. Proteasome inhibition led to a modest relative increase in cytoplasmic POMP and a much larger relative increase in nuclear POMP. (O) Analysis of POMP levels in Western blots of fractionation experiments like that shown in N. MG132 led to a modest but significant relative increase in cytoplasmic POMP and large and significant relative increase in nucleolar POMP. **p≤0.01, paired two-tailed t-tests on non-normalised data, n=3, mean±SD, FC=fold change. (P) Analysis of pro-PSMB5 levels in Western blots of fractionation experiments like that shown in N. MG132 did not elicit a significant change in pro-PSMB5 in either compartment. ns=p>0.05, paired two-tailed t-tests on non-normalised data, n=3, mean±SD, FC=fold change.

The above data indicate that, under normal conditions, POMP is localized to a diverse set of vesicles of the secretory pathway. Next, we asked whether POMP’s intracellular distribution changes in response to proteasome inhibition. Strikingly, HEK293 and primary neurons treated with proteasome inhibitors exhibited a relocalisation of POMP to the nucleolus and nuclear bodies, revealed by immunofluorescence (Figure 3j-m). To examine whether the nuclear localization of POMP was uncoupled from its proteasomal partners, we performed cytoplasmic-nuclear fractionation on HEK293 treated with MG132 and tested the fractions by Western blot analysis. The fractions collected exhibited a clear separation of cytoplasm (GAPDH) and nuclei (Histone H3) (Figure 3n). We measured the change in POMP and its interaction partner pro-PSMB5 in the cytoplasmic and nuclear compartments following proteasome inhibition (MG132). While neither the levels nor the intracellular distribution of pro-PSMB5 changed, POMP levels increased in both the cytoplasmic and nuclear compartments in response to proteasome inhibition (Figure 3n-p). Importantly, proteasome inhibition resulted in a much greater relative increase in nuclear POMP (∼4-fold) than cytoplasmic POMP (∼1.5-fold) (Figure 3o,p). This experiment suggested that the increased POMP levels produced by preventing its turnover as well as (HSF1-dependent) transcription resulted in its translocation into the nucleolus and nuclear bodies.

The proteasome-independent relocalisation of POMP prompted us to test whether the increased POMP that results from proteasome inhibition forms novel protein complexes. It is known, for example, that in response to prolonged proteotoxic stress (including proteasome inhibition for 12 hrs^48^) protein-RNA aggregates (nucleolar aggresomes) form within the nucleolus. We first confirmed that our relatively brief (5 hr) period of proteasome inhibition did not result in appreciable nuclear aggresome formation (Figure S3B). To identify and characterize other auxiliary complexes that POMP might associate with, we treated primary neurons with MG132, performed sucrose density gradient ultracentrifugation and ran non-reducing SDS-PAGE and Western blots of the different fractions staining for proteasome components, POMP and ribosomal proteins to estimate the size of novel complexes (Figure 4a). We reasoned that if POMP binds to complexes or factors distinct from the proteasome, then there should be a shift in POMP distribution across the fractions, independent of proteasome components (Figure 4a). The distribution and total levels of proteasomes, exemplified by the core particle component PSMA3 and the regulatory particle subunit PSMC3, were minimally affected and did not shift across the fractions (Figure 4b,c). By contrast, overall POMP levels increased and its distribution shifted to heavier fractions (Figure 4b,c). Additionally, in heavier fractions we observed the appearance of higher molecular weight forms of POMP following treatment with MG132 (Figure 4b).

**Figure 4.**
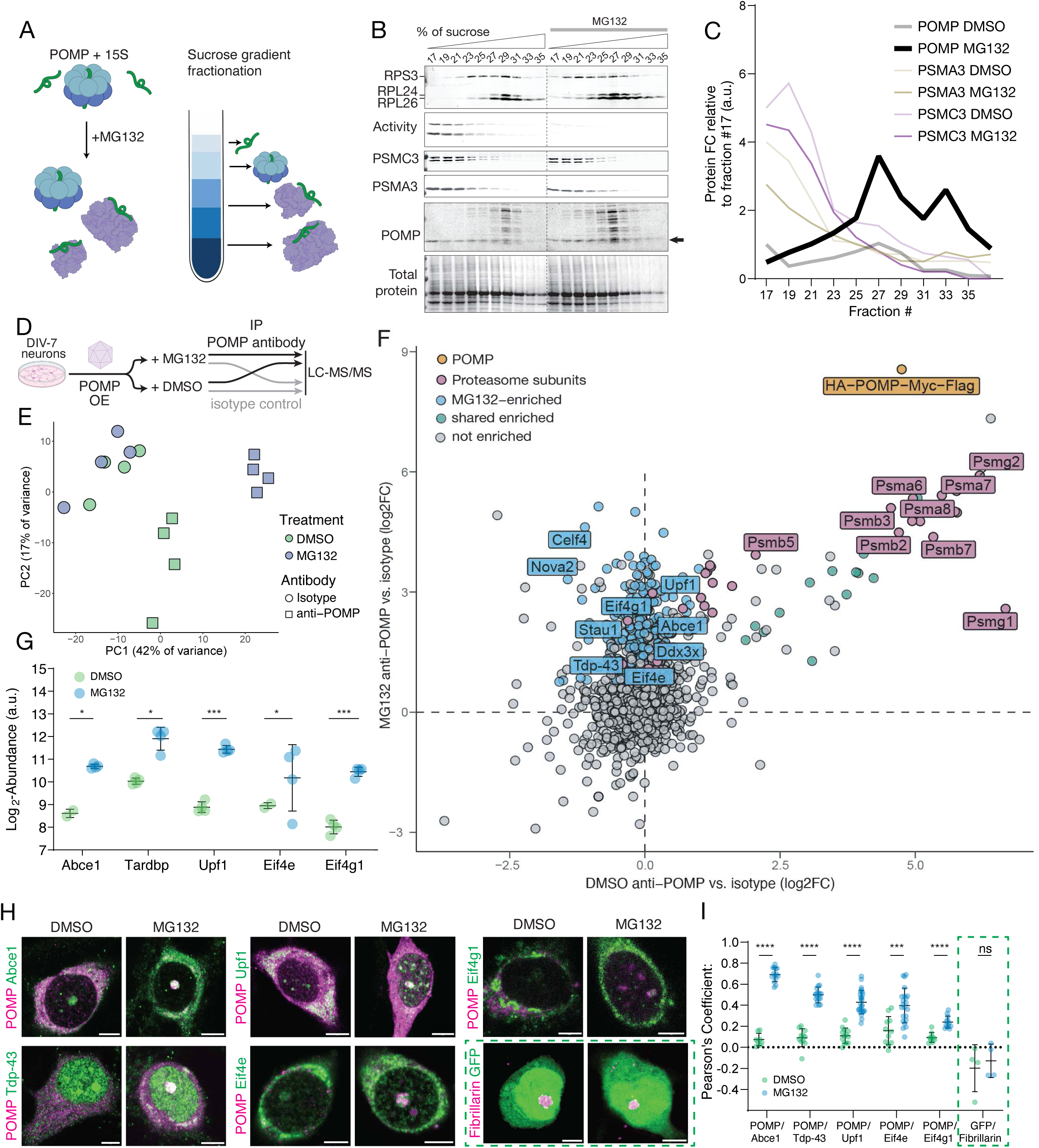
Proteasome inhibition-dependent nucleolar relocalisation rewires the POMP interactome. (A) Scheme of sucrose density fractionation experiment performed in neurons to identify changes in POMP’s localization and interaction partners associated with proteasome inhibition. (B) Non-reducing SDS PAGE Western blot analysis of gradient fractions 17-35 obtained from neurons treated (right) or not (left) with MG132 for 5 hrs. Antibodies to ribosomal proteins RPS3 and RPL24 and 26 were used to estimate the size of the complexes. Proteasome inhibition led to the expected loss in activity (Activity) and did not appreciably alter the position of the proteasome subunits (PSMA3 and PSMC3) in the gradient. In contrast, POMP displayed an activity-dependent increase and right-ward shift towards heavier fractions in the gradient, as well as an increase in its occupancy in higher molecular weight complexes. Arrow points to the POMP monomer band used for the analysis in C. (C) Analysis of the Western blot experiments like that shown in B. Unlike all other proteins examined, POMP exhibited an increase and shift towards heavier fractions in the gradient following proteasome inhibition. (D) Scheme of the immunoprecipitation-mass spectrometry experiment. Cortical neurons were transduced with AAVs expressing either POMP or GFP, and after two weeks in culture they treated with DMSO or MG132 for 5 hr and used to make cell extracts for IP with an antibody against POMP or an isotype control and eluates were analysed by LC-MS/MS. (E) Principal Component Analysis (PCA) plot of the IP-MS experiment depicted in D. Each point represents an individual biological replicate, colored according to treatment and shaped according to antibodies used for IP. While POMP-IP samples cluster apart in response to treatment, isotype control samples do not. n=4 biological replicates. (F) Scatter plot of protein enrichment in POMP antibody IP versus isotype control under control and treatment conditions. Each point represents a protein. X-axis shows the log₂FC of the POMP antibody over isotype control in DMSO control condition, and the y-axis shows the log₂FC in the MG132 condition. Proteins enriched in both conditions, such as proteasome components, appear along the diagonal, while those preferentially enriched in response to proteasome inhibition deviate from the diagonal. (G) Analysis of the protein abundance measured in the IP-MS experiment of cortical neurons treated with DMSO or MG132 for 5h. Shown is a subset of the detected interactors with known roles in RNA biogenesis. *p≤0.05, ***p≤0.001, significance was assessed using model-based statistics in MSstats (v4) based on linear-mixed effects models with Benjamini-Hochberg correction for multiple comparisons, n=4 biological repeats, mean±SD. (H) Representative immunofluorescence images of hippocampal neurons expressing the HA-POMP-Myc-Flag construct for 24 hr prior to treatment with DMSO or MG132 for 5h. The cells were stained for the newly identified interactors shown in G and myc-tag (indicated as POMP). Scale bars, 5 µm. (I) Pearson’s correlation analysis of the POMP immunofluorescence signal with that of its newly identified interactors in the nucleus of neurons, as shown in panel H. In all cases proteasome inhibition leads to relocalisation of POMP to the nucleolus with a significant increase in the correlation between the two immunofluorescence signals in the nucleus. ****p≤0.0001, unpaired two-tailed t-tests, n= 8 (Abce1, DMSO), 14 (Abce1, MG132), 11 (Tdp-43, DMSO), 17 (Tdp-43, MG132), 13 (Upf1, DMSO), 26 (Upf1, MG132), 11 (Eif4e, DMSO), 22 (Eif4e, MG132), 9 (Eif4g1, DMSO), 12 (Eif4g1, MG132) and 4 (GFP DMSO and MG132) neurons, mean±SD.

Thus far, the data suggest that in response to a proteostatic challenge POMP forms new protein complexes associated with the nucleolus. To identify POMP’s new binding partners, we used an immunoprecipitation mass spectrometry (IP-MS) strategy with a tagged version of POMP, optimizing our construct for neuronal expression and its ability to successfully integrate into hemi-proteasomes, expressed from a plasmid (Figure S4a-c) and once packaged into an adeno-associated virus (Figure S4d-g). AAV-transduced neurons were treated with DMSO or MG132, then POMP was immunoprecipitated and the resulting eluates were analyzed by LC-MS/MS. Principal component analysis revealed a reconfiguration of the POMP interactome following proteasome inhibition: the POMP IP samples from MG132-treated neurons clearly clustered away from the control neurons; this pattern was not observed in isotype control IP samples (Figure 4e). We first quantified the proteins enriched in the POMP IP compared to the control. As expected, POMP itself, along with multiple proteasome subunits, was significantly enriched in both control and MG132 conditions (Figure 4f). To identify the proteins that were differentially enriched upon MG132 treatment, we applied a linear model to normalize each condition against its isotype control and assess the differential enrichment. This analysis revealed a specific group of POMP-interacting proteins uniquely enriched in response to proteasome inhibition (Figure 4f). Among the new factors identified were many RNA-binding proteins, including Stau1, Celf4, Ddx3x, Upf1, Tdp-43, Abce1, Eif4e and Eif4g1 (Figure 4f) and analysis of the inputs confirmed that the interaction was not due to simple accumulation of these proteins in response to MG132 (Figure S4h). GO term analysis confirmed that the terms ‘RNA binding’ and ‘mRNA binding’ were significantly overrepresented (Figure S4i). These findings indicate that proteasome inhibition leads to a rewiring of the POMP interactome, resulting in the recruitment of RNA-binding proteins. To examine the cellular localization of these MG132-dependent interactions and validate the IP-MS results, we evaluated a panel of interacting RNA-binding proteins, including the ribosome biogenesis factor Abce1, the RNA-processing regulator Tdp-43, the RNA helicase Upf1 and the translation initiation factors Eif4e and Eif4g1, which were recently found to have roles in pre-mRNA splicing and transcriptional regulation^49–51^ (Figure 4g). We tested whether POMP and the above interactors colocalized in the nucleolus in response to proteasome inhibition using immunofluorescence (Figure 4h,i). For each of the 5 interactors, we observed a significant increase in their spatial overlap with POMP in the nucleus (Figure 4i). Altogether, these data revealed that early in the response to proteasome inhibition, POMP accumulates in the nucleolus and other nuclear bodies, interacting with RNA-binding proteins.

One surprising aspect of the IP-MS experiment was that the rewiring of the POMP interactome was *only* observed following proteasome inhibition and was not observed under control conditions, despite the “overexpression” of the tagged POMP. We thus queried directly whether POMP overexpression alone is sufficient to trigger nuclear puncta formation. To do so, we expressed the tagged POMP in HEK293 cells and treated them with either DMSO or MG132. We directly quantified the POMP levels and observed that, despite an ∼8-fold increase in cellular POMP levels following expression of the tagged POMP compared to MG132 treatment alone (Figure 5a,b), overexpression alone was not sufficient to induce nuclear accumulation (Figure S5a) or puncta formation (Figure 5a,c). To determine whether the explicit localization of POMP to the nucleus by intrinsic protein features can drive puncta formation, we generated a POMP fusion construct with a nuclear localization signal (POMP-NLS). While the expression of the POMP-NLS was sufficient to elevate nuclear POMP to the same levels achieved by treatment with MG132 (Figure 5d,e), it also did not lead to formation of nuclear puncta (Figure 5d,f and Figure S5b-d).

**Figure 5.**
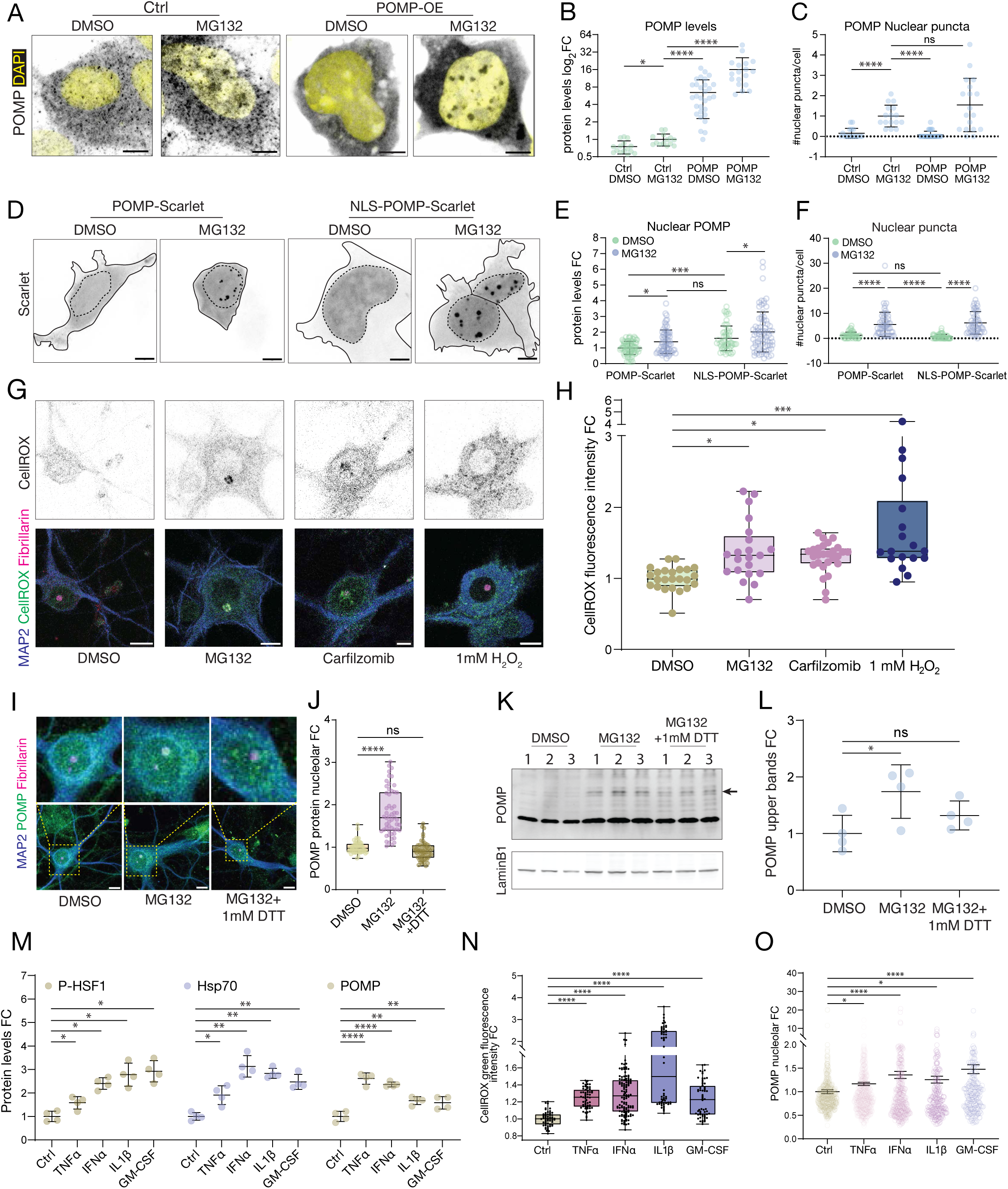
Formation of POMP nuclear puncta is context-dependent and requires an elevation in cellular ROS. (A) Representative immunofluorescence images of HEK293 cells expressing either an empty vector control or a POMP plasmid for 72h treated with either DMSO or MG132 for 5h. The formation of POMP nuclear puncta is observed only in response to proteasome inhibition and is independent of POMP overexpression. Scale bars = 5 µm. (B) Analysis of the POMP expression levels in the HEK293 cells of the experiment shown in A. While proteasome inhibition leads to a modest but significant increase in POMP levels, POMP overexpression in combination with proteasome inhibition leads to significant and progressively large increase in cellular POMP levels. *p≤0.05, ****p≤0.0001, Welch’s ANOVA test and post-hoc Dunnett’s T3 multiple comparisons test, n=15 (Cntrl DMSO), 14 (Cntrl MG132), 30 (POMP DMSO) and 18 (POMP MG132), mean±SD, FC=fold change. (C) Analysis of the number of nuclear POMP puncta in the HEK293 cells of the experiment shown in A. The formation of POMP nuclear puncta is independent of POMP expression levels in cells and only proteasome inhibition leads to a significant increase in their number. ns=p>0.05, ****p≤0.0001, Welch’s ANOVA test and post-hoc Dunnett’s T3 multiple comparisons test, n=17 (Cntrl DMSO), 16 (Cntrl MG132), 30 (POMP DMSO) and 18 (POMP MG132), mean±SD. (D) Representative immunofluorescence images of HEK293 cells expressing either POMP-Scarlet or NLS-POMP-Scarlet (nuclear localised) overexpression plasmids for 72h treated with either DMSO or MG132 for 5h. Enforcing nuclear localisation of POMP was not sufficient to drive puncta formation, which requires proteasome inhibition.Scale bars = 5 µm. (E) Analysis of the nuclear POMP expression levels in the HEK293 cells of the experiment shown in D. Overexpression of POMP-Scarlet +/- MG132 and NLS-POMP-Scarlet +/- MG132 led to a significant and progressive rise in nuclear POMP levels. ns=p>0.05, *p≤0.05, ***p≤0.001, one-way ANOVA and post-hoc Tukey’s multiple comparisons test, n= 76 (POMP-Scarlet DMSO), 70 (POMP-Scarlet MG132), 56 (NLS-POMP-Scarlet DMSO), 76 (NLS-POMP-Scarlet MG132), mean±SD, FC=fold change. (F) Analysis of the number of nuclear POMP puncta in the HEK293 cells of the experiment shown in D. The formation of POMP nuclear puncta was independent of nuclear POMP levels in cells and only proteasome inhibition led to a significant increase in their number. ns=p>0.05, ****p≤0.0001, one-way ANOVA and post-hoc Tukey’s multiple comparisons test, n=50 (POMP-Scarlet DMSO), 55 (POMP-Scarlet MG132), 54 (NLS-POMP-Scarlet DMSO), 54 (NLS-POMP-Scarlet MG132), mean±SD. (G) Representative immunofluorescence images of hippocampal neurons treated with either DMSO, the proteasome inhibitors MG132, Carfilzomib or the ROS precursor H₂O₂, and stained for MAP2, the nucleolar marker Fibrillarin and the oxidative stress probe CellROX Green. Treatment with H₂O₂ and proteasome inhibition lead to an increase in cellular ROS levels, measured by the increase in CellROX Green fluorescence. Scale bars = 5 µm. (H) Analysis of the CellROX Green fluorescence intensity in the hippocampal neurons of panel G. Treatment with H₂O₂ and proteasome inhibitors led to a significant increase in CellROX Green fluorescence. *p≤0.05, ****p≤0.0001, one-way ANOVA and post-hoc Dunnett’s multiple comparisons test, n=24 (DMSO), 21 (MG132), 27 (Carfilzomib) and 18 (H₂O₂), Boxplots show the median (line), interquartile range (box), and Min-Max whiskers. FC=fold change. (I) Representative immunofluorescence images of hippocampal neurons treated with either DMSO, MG132 alone or MG132 in combination with the reducing agent DTT (1 mM), and immunostained for POMP, MAP2 and the nucleolar marker Fibrillarin. Preventing ROS production with DTT blocked POMP relocalisation to the nucleolus. Scale bar in the low and high-magnification images 10 and 5 µm, respectively. (J) Analysis of neuronal nucleolar POMP levels in the the experiment in I. While proteasome inhibition led to a significant increase in nucleolar POMP levels, blocking ROS production by DTT treatment prevented the nucleolar relocalisation of POMP. ns=p>0.05, ****p≤0.0001, one-way ANOVA and post-hoc Dunnett’s multiple comparisons test, n=54, Boxplots show the median (line), interquartile range (box), and Min-Max whiskers, FC=fold change. (K) Representative non-reducing SDS-PAGE Western blot analysis of cortical neurons treated with either DMSO, MG132 or MG132+1 mM DTT for 5h. Treatment with MG132 led to elevated POMP levels and the appearance of higher molecular species that were redox sensitive and reduced by co-incubation of MG132 with 1 mM DTT on cells. LaminB1 was used as loading control. (L) Analysis of the Western blot shown in K. DTT blocked the relocalization of POMP. The discrepancy in the size of the effects seen for POMP relocalisation I,J and POMP higher molecular weight species levels in K and L can be explained by the re-oxidation POMP during non-reducing SDS-PAGE in atmospheric oxygen. ns=p>0.05, *p≤0.05, RM one-way ANOVA and post-hoc Dunnett’s multiple comparison test, n=4, mean±SD. (M) Analysis of the Western blots shown in Figure S5Q of HEK293 treated with pro-inflammatory cytokines (TNFα, INFα, IL1β, GM-CSF) or water control for four days. Treatment with all four cytokines leads to a significant upregulation in P-HSF1 and its downstream targets Hsp70 and POMP. *p≤0.05, **p<0.01, ****p≤0.0001, RM one-way ANOVA and post-hoc Dunnett’s multiple comparison test, n=4, mean±SD. FC=fold change. (N) Analysis of the CellROX Green fluorescence intensity in HEK293 treated with pro-inflammatory cytokines for four days. In all cases treatment leads to a significant increase in CellROX Green fluorescence. ****p≤0.0001, one-way ANOVA and post-hoc Kruskal-Wallis multiple comparisons test, n=57 (Cntrl), 60 (TNFα, IL1β, GM-CSF), 120 (INFα), Boxplots show the median (line), interquartile range (box), and Min-Max whiskers. FC=fold change. (O) Analysis of the nucleolar POMP levels in HEK293 treated with pro-inflammatory cytokines for four days. Treatment with all four cytokines leads to a significant increase in nucleolar POMP levels. *p≤0.05, ****p≤0.0001, one-way ANOVA and post-hoc Dunnett’s multiple comparison test, n=712 (Cntrl), 795 (TNFα), 472 (IFNα), 291 (IL1β), 272 (GM-CSF), mean±SD. FC=fold change.

The insufficiency of a variety of manipulations (which either increase cellular POMP levels or directly promote its nuclear localization) to drive nuclear POMP puncta formation suggested that proteasome inhibition may also induce a unique cellular context. It is known, for example, that nucleolar oxidation occurs in response to a variety of cellular stressors^52^. We thus examined whether proteasome inhibition changes the cellular oxidative environment using a fluorescent probe sensitive to reactive oxygen species (ROS). We observed that the ROS probe’s intensity increased following treatment with proteasome inhibitors in both primary neurons (Figure 5g,h) and HEK293 cells (Figure S5e,f). In neurons, in particular, proteasome inhibition led to a distinct accumulation of the probe in the nucleolus (Figure 5g). Redox-sensitive amino acids include Cys, Met and His and POMP is known to harbor a highly conserved Cys residue^46,53^ (Figure S5g). In addition, the Cys in POMP/Ump1 is highly reactive and readily forms disulfide bonds and oligomeric species upon exposure to atmospheric oxygen^46,53^, reminiscent of the high molecular weight POMP species observed in MG132-treated samples (Figure 4b). To test whether the MG132-induced high molecular weight POMP species were sensitive to ROS we used the reducing agent dithiothreitol (DTT). Extracts from neurons treated with proteasome inhibitors were analysed by (non-reducing and reducing) SDS-PAGE and Western blot (Figure S5h). We observed that the higher molecular weight POMP species, induced by proteasome inhibition, were modestly, but significantly, sensitive to DTT treatment (Figure S5i-k). In independent experiments, we determined that a mutant version of POMP unable to form disulfide bonds (C36A) was compromised in its ability to form POMP oligomers (Figure S5l-p). The above data indicate that proteasome inhibition leads to an increase in cellular reactive oxygen species, particularly in the nucleolus, and that POMP is redox-sensitive and capable of forming oligomers in response to proteasome inhibition. We next tested directly whether preventing oxidative stress blocks the nucleolar accumulation of POMP. We incubated neurons with either DMSO, MG132 alone or MG132 and DTT and performed immunofluorescence imaging and Western blot analysis (Figure 5i-l). Immunofluorescence imaging revealed that 1 mM DTT was sufficient to prevent nucleolar accumulation of POMP (Figure 5i,j). Western blot analysis revealed an MG132-dependent accumulation of POMP oligomers and a significant reduction in their levels by treatment of cells with 1 mM DTT (Figure 5k,l). Taken together, these results show that upon nuclear relocalization POMP can oligomerize via disulfide-bond formation of its cysteine residue, C36.

Are there additional stimuli that can elicit this relocalization of POMP? Previous work has shown that POMP is strongly induced by cytokines in the context of inflammation^28,33,54^ and leads to a transcriptional switch from the constitutive to immunoproteasome^33^. We treated HEK293 with four different pro-inflammatory cytokines (TNFα, IFNα, TGFβ and GM-CSF) and assessed the levels of POMP, its transcriptional regulators, the immunoproteasome and constitutive 20S proteasome subunits, and proteasome activity (Figure S5q). Cytokine treatment led to strong accumulation of POMP without a corresponding increase in constitutive proteasome subunits, and only a small induction of immunoproteasome subunits (i.e. PSMB8, PSMB9, PSMB10) (Figure 5m and Figure S5q), thus leading to no detectable change in proteasome activity pattern or levels (Figure S5q). All 4 cytokines also led to a significant increase in oxidative stress levels (Figure 5n). Consistent with our model, the accumulation of ROS and increased POMP levels led to nucleolar relocalisation of POMP (Figure 5o), demonstrating that this phenomenon can occur in the absence of proteasome inhibition.

Taken together the above data indicate that during proteasome dysfunction and redox stress, POMP re-localises to the nucleolus and nuclear bodies and engages with RNA biogenesis factors, independent of its conventional proteasome assembly partners. What are the functional consequences of this new subcellular location? Because perturbing POMP has a direct effect on proteasome levels and function, we devised a strategy to disentangle proteasome-dependent and -independent transcriptional effects of POMP knockdown. To do so, we compared the effects of POMP knockdown (via siRNA) to knockdown of the proteasome (siRNA of PSMB1 or PSMB7) (Figure 6a). In doing so, we achieved one condition with *reduced* POMP levels (siRNA-POMP; “POMP-”) and another with *elevated* POMP levels (siRNA-PSMB; “POMP+”) (Figure S6A a-f) whilst achieving the same level of proteasome impairment in both groups (Figure 6a,b and Figure S6A a-f). The knock-down of POMP led to a distinct transcriptomic response, as revealed by a principal components analysis (Figure S6A g). Differential expression analysis between the siRNA-POMP and siRNA-PSMB conditions revealed that while POMP’s persistence (during knockdown of PSMB subunits; POMP+) resulted in the upregulation of ribosomal transcripts (e.g., *RPS29*, *RPSA*, *RPL30*, *RPL22*, and *RPL34*) and components of the respiratory chain (e.g., *ATP5MC1*, *COX7C*), POMP knockdown (POMP-) induced a marked increase in the expression of extracellular matrix metalloproteinases (e.g., *MMP1* and *10*), oncogenic transcription factors (e.g., *FOS*, *MYC*, *JUNB*), and inflammatory cytokines (e.g., *CCL2*, *CCL7*) (Figure 6c). We note that in the absence of POMP the transcriptional upregulation of CP subunits was significantly reduced compared to conditions where POMP was retained (Figure S6A h). Gene ontology (GO) and KEGG pathway enrichment analysis revealed that loss of POMP was associated with the activation of cancer-, inflammation-, and immunity-related pathways (Figure 6d and Figure S6A i), while retention of POMP was linked to global changes in cellular metabolism, upregulation of ribosomal subunits, and gene expression profiles associated with neurodegenerative diseases (Figure 6e and Figure S6A j), such as upregulation of SOD1 (Figure 6c). These data indicate that, under conditions of proteasome inhibition, POMP drives a transcriptional program that prevents metabolic decline of the cell, by preserving ribosome biogenesis and cellular respiration, while suppressing the activation of an inflammation-like signaling axis (Figure 6c-e and Figure S6A k-m).

**Figure 6.**
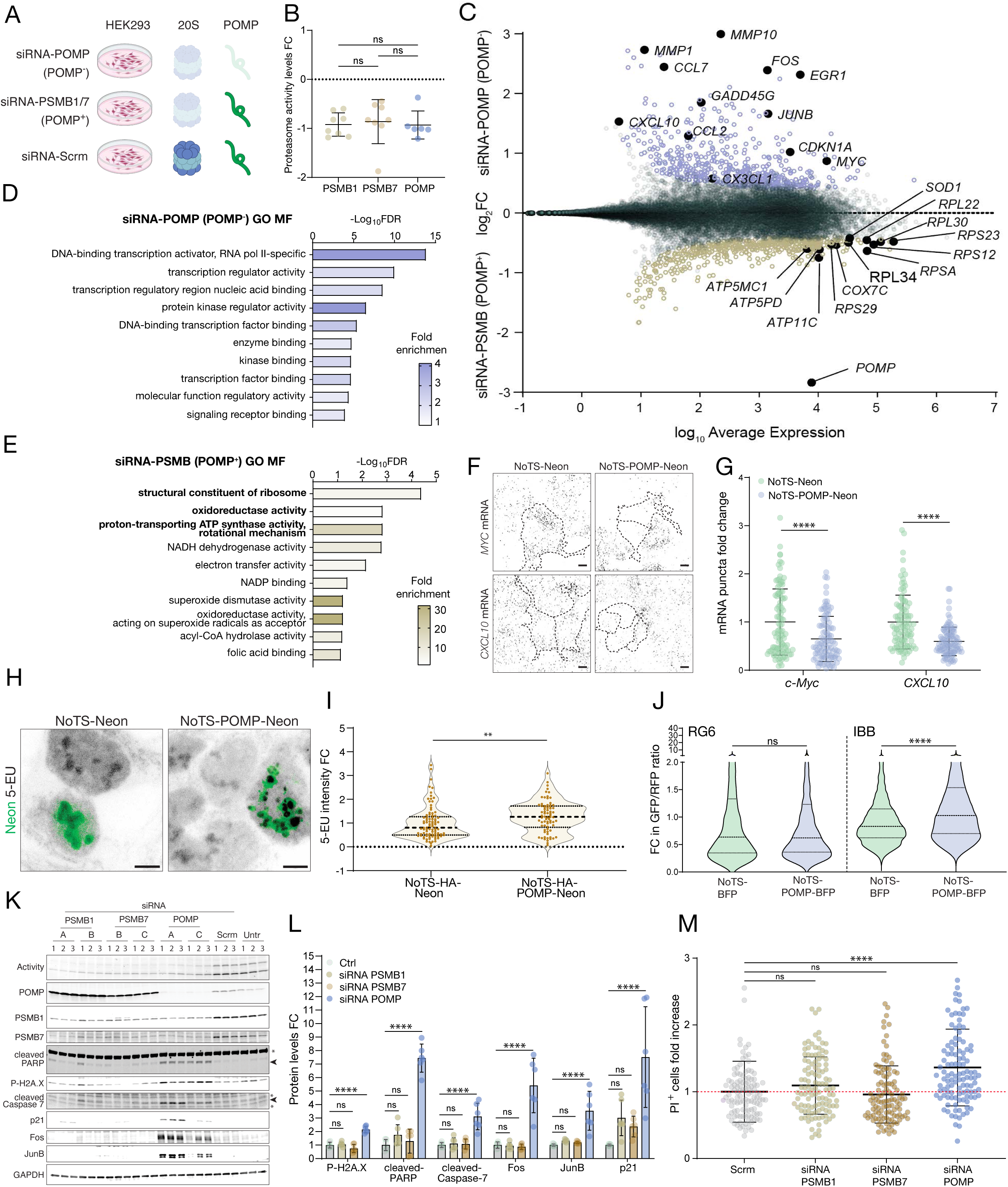
Nucleolar POMP drives a pro-survival and positive metabolic transcriptional program during proteasome inhibition. (A) Scheme of siRNA-based approach to isolate POMP’s proteasome independent actions on the transcriptome by creating two conditions where proteasome activity is reduced to the same extent: one in which POMP is absent (POMP-) and one in which POMP is present (POMP+). (B) Analysis showing that proteasome activity in the different siRNA conditions (see Figure S6A a-f) was reduced to the same extent across conditions. For every gene, results obtained with two different siRNAs were pooled together for analysis. ns=p>0.05, one-way ANOVA and post-hoc Tukey’s multiple comparison test, n=8 (PSMB1/7), 6 (POMP), mean±SD. FC=fold change. (C) Differential expression analysis of the HEK293 cell transcriptome under conditions where the proteasome was inhibited and POMP was either present (POMP+) or absent (POMP-). Y-axis depicts the differential expression of transcripts; x-axis depicts the average expression level. Transcripts that were significantly upregulated in POMP+ and POMP- are labelled in gold and blue, respectively. (D) Gene Ontology (GO) molecular function (MF) analysis of transcripts upregulated in POMP-condition. The x-axis depicts the statistical significance of the enrichment (-log_10_FDR) and the colour depicts fold enrichment. FDR<0.05, Fisher’s exact test. (E) Gene Ontology (GO) molecular function (MF) analysis of transcripts upregulated in POMP+ condition. The x-axis depicts the statistical significance of the enrichment (-log_10_FDR) and the colour depicts fold enrichment. FDR<0.05, Fisher’s exact test. (F) Representative images of HEK293 cells transfected with either a construct encoding a nucleolar-targeting signal (NoTS) fused to the fluorescent protein Neon (control) or to POMP-Neon and then probed for the *CXCL10* or *Myc* mRNA, using *in situ* hybridization (black signal). Dashed lines outline the perimeter of transfected cells for comparison to neighboring, untransfected cells. Scale bars = 5 µm. (G) Analysis of the experiments like that shown in F. The expression of NoTS-POMP-Neon significantly reduced both *CXCL10* and c-*Myc* mRNA. ****p≤0.0001, unpaired two-tailed t-tests, n=100, mean±SD, FC=fold change. (H) Representative images of HEK293 cells transfected with either a construct encoding a nucleolar-targeting signal (NoTS) fused to the fluorescent protein Neon (control) or to POMP-Neon cultured for 72 hr prior to 1 hr incubation with 5-EU to label nascent RNA (black signal). The green signal indicates Neon fluorescence. Scale bars= 5 µm. (I) Analysis of the nucleolar nascent RNA experiments like that shown in H. The expression of NoTS-POMP-Neon significantly enhances the nascent RNA signal in the nucleolus compared to NoTS-Neon. ****p<0.0001, unpaired two-tailed t-tests, n=92 (NoTS-Neon), 74 (NoTS-POMP-Neon) cells from 3 experiments. Violin plots show median, interquartile range, minimum and maximum values, FC=fold change. (J) Analysis of the splicing reporter experiments in which expression of NoTS-POMP resulted in significant altered splicing (exon retention; higher GFP:RFP ratio) of the TDP-43-sensitive (IBB) but not the general (RG6) splicing reporter. ****p≤0.0001, Mann-Whitney U tests, n = 2852 (NoTS-BFP, RG6), 3843 (NoTS-POMP-BFP, RG6), 3480 (NoTS-BFP, IBB), 4025 (NoTS-POMP-BFP, IBB) cells from 3 experiments. (K) Western blot analysis for the indicated proteins under control (untreated and scramble siRNA) or PSMB1/7 or POMP siRNA knockdown conditions. Letters refer to different siRNA constructs, numbers refer to biological replicates. Arrowheads point to the cleaved products of Caspase 7 or PARP, * points to non-specific bands in the vicinity of the target band. (L) Analysis of the experiment shown in K. For all of the proteins or protein forms indicated on the x-axis, POMP siRNA, but not PSMB1/7 siRNA, led to a significant increase in the protein. ****p≤0.0001, one-way ANOVA with post-hoc Dunnett’s test, n= 6, 2 different siRNA constructs with 3 biological replicates, mean±SD. FC=fold change. (M) Analysis of cell death experiments using propidium iodide (PI). POMP siRNA led to a significant increase in the number of PI+-cells whereas PSMB1/7 siRNA had no effect. ****p<0.0001, one-way ANOVA with post-hoc Dunnett’s test, n=113 (Scrm), 114 (PSMB1/4, POMP), mean±SD. FC=fold change.

We conducted additional experiments to address whether POMP’s nucleolar localization plays a direct role in regulating the above transcripts. First, we selected two transcripts, the pro-inflammatory cytokine *CXCL10*, and the proto-oncogene *Myc*, which were upregulated in response to loss of POMP (Figure 6c) and whose RNA biogenesis has been reported to involve the nucleolus^55,56^. We tested whether overexpression of POMP fused to a nucleolar targeting signal (NoTS) is sufficient to reduce their expression, as assessed by *in situ* hybridization. We validated construct expression (Figure S6B a-c) and found that POMP-NoTS expression reduced both *CXCL10* and *Myc* (Figure 6f,g). We next asked whether the expression of POMP-NoTS alone can increase the levels of transcripts specifically upregulated (Figure 6c) following PSMB knockdown (POMP+). To do so, we examined the ribosomal proteins RPSA, RPS3A, RPS23, RPL22 and RPL34. Immunofluorescence analysis revealed that expression of POMP-NoTS was sufficient to elevate cellular levels of these ribosomal subunits (Figure S6B d,e). Ribosomal proteins are particularly susceptible to oxidative damage^57^. The observation that under conditions of proteasome inhibition (associated with elevated ROS - Figure 5g,h and Figure S5e,f) increased POMP preserves ribosome biogenesis suggests a protective role of POMP in maintaining proteostasis. As ribosome biogenesis requires ribosomal proteins and rRNA, however, a truly protective mechanism should also induce rRNA synthesis to avoid a stoichiometric imbalance. We thus expressed POMP-NoTS and performed metabolic (5-EU) labelling to measure RNA synthesis in the nucleolus. We found that POMP-NoTS is sufficient to drive an increase in nucleolar nascent RNA (Figure 6h,i), suggesting that nucleolar POMP serves to coordinate ribosome biogenesis by promoting both rRNA and ribosomal protein synthesis under stress conditions.

Next, we surveyed the gene features of the differentially (siRNA-POMP vs siRNA-PSMB) regulated transcripts and found that the two gene sets had contrasting properties: POMP retention favoured transcripts with shorter CDS, 5’ and 3’ UTRs and a lower GC content whereas POMP loss favoured the opposite (Figure S6C a). A linear mixed model confirmed that GC content was the major contributor of the transcriptomic differences between the conditions (Figure Sup 6C b,c). Feature enrichment analysis revealed that POMP retention was associated with the expression of genes with multiple (>4) exons, a high number of annotated alternative isoforms, and evidence for regulation by splicing (Figure S6C d-f). We thus performed differential transcript usage analysis and found that POMP retention (siRNA PSMB, POMP+) was uniquely associated with higher expression of transcript isoforms associated with cellular response to stress and DNA damage (GO; Biological process; Figure S6C f,g). These findings, together with the observation that many of the newly identified POMP interactors have roles in splicing (e.g. *Upf1*, *TDP-43*, *Ddx3x*), prompted us to directly test the effect of nucleolar-targeted POMP on splicing. Expression of POMP-NoTS did not alter splicing of the general splicing reporter RG6^58^ (Figure 6j and Figure S6C h). However, the TDP-43-specific reporter, based on exon 9 of the CFTR gene, which can be skipped only by the action of TDP-43^59^ (identified in our IP-MS dataset as an MG132-induced POMP interactor; Figure 4f) displayed higher rates of exon retention following overexpression of NoTS-POMP (Figure 6j and Figure S6C h). To verify that changes in splicing/transcript isoform usage resulted in changes in protein isoforms we selected *PARP3* (Figure S6C e,i) which is also under the control of TDP-43^60^ and exhibits isoform usage regulation^61,62^. Western blot analysis confirmed that in response to PSMB knockdown, but not POMP knockdown, a lower molecular weight isoform of PARP3 was specifically increased (Figure S6C j,k). These data suggest that nucleolar POMP alters splicing of a specific subset of genes working together with its interactors, including TDP-43.

Altogether these data suggest that POMP relocalisation in response to proteasome inhibition elicits a specific transcriptional signature by favouring certain transcripts based on their GC content (Figure S6C a-c) and influencing the splicing/isoform usage of a subset of genes, which play a protective role in response to proteostatic stress (Figure S6C e-k). In fact, the concomitant upregulation of proto-oncogenes, pro-apoptotic factors and cell-cycle inhibitors is a common response profile of stressed cells which eventually undergo apoptosis^63–65^. Using Western blot analysis, we validated the increased levels of oncogenes and further examined the commitment to apoptosis of cells with impaired proteasome function that either retain (siRNA PSMB) or have lost (siRNA POMP) POMP (Figure 6k,l). Consistent with the sequencing data, we found that POMP knockdown displayed a unique and strong induction of the FOS, JUNB and the p21 proteins (Figure 6k,l). To visualize directly whether POMP’s absence in a proteostatically stressed context can lead to apoptosis we probed for the DNA-damage marker P-H2A.X and the cleaved forms of the apoptotic markers Caspase7 and PARP (Figure 6k,l). We found that POMP knockdown (but not PSMB knockdown) upregulated these markers. To directly detect cell death we used Propidium iodide (PI) staining and observed that, unlike proteasome knockdown, POMP knockdown triggered apoptosis (Figure 6m). Collectively, these data indicate that in response to a diverse set of cellular insults, POMP levels increase and promote a protective pathway that prevents the initiation of programmed cell death.

The newly identified pro-survival moonlighting function of POMP suggests that its regulation could confer advantages to cells that are exposed to constant proteostatic stress, such as stem cells in the ongoing processes of self-renewal and differentiation. We thus analyzed two different mouse embryonic stem cell (mESCs; E14Tg2A and R1) lines for differences in both basal and proteasome inhibitor-induced POMP levels and compared them to HEK293 and neurons, using both Western blot analysis and immunofluorescence (Figure 7a,b and Figure S7a,b). Surprisingly, we observed a flip in the response to proteasome inhibition-instead of the typical increase in POMP levels seen in HEK293 and neurons, the mESCs exhibited a substantial decrease in POMP levels (Figure 7a,b and Figure S7a,b). Despite this, POMP exhibited the same expected rapid turnover in mESCs and neurons (Figure S7a,b). In addition, treatment with a proteasome inhibitor (MG132 or Carfilzomib) failed to induce nucleolar localization of POMP in mESCs (Figure 7c,d), though it still promoted the generation of ROS (Figure 7e,f). To examine whether the dramatically different response of mESC POMP to proteasome inhibition is developmental stage-sensitive, we differentiated mESCs into functional active neurons (Figure S7c-h). We then compared the responses of differentiated neurons and the parent mESCs to proteasome inhibition. While proteasome inhibition led to a decrease in proteasome activity in both the mESCs and the neurons (Figure 7g,h), the differentiated neurons, like primary neurons (and all other cells we have tested), increased POMP levels (Figure 7g, i). The levels of the proteasome subunit PSMB5 and its precursor form (pro-PSMB5) remained unaffected by treatment in both cell types (Figure S7i-k). We reasoned that the observed mESC reduction in POMP could be explained by turnover by the other major cellular degradation pathway, autophagy and tested this using the inhibitor chloroquine (Figure 7j,k). Indeed, we found that blocking autophagy prevented the POMP protein decrease that occurs in response to proteasome inhibition (Figure 7j,k). The observed autophagic reduction in POMP is likely selective as markers of bulk autophagic flux (LC3B-II and p62) were similarly affected by proteasome inhibition in both mESCs and neurons (Figure S7l-n). We also noted that the colocalization of POMP and the autophagosome marker ATG5 was much higher in the mESCs than the neurons (Figure S7o,p), suggesting a pre-existing disposition for POMP degradation by autophagy in mESCs.

**Figure 7.**
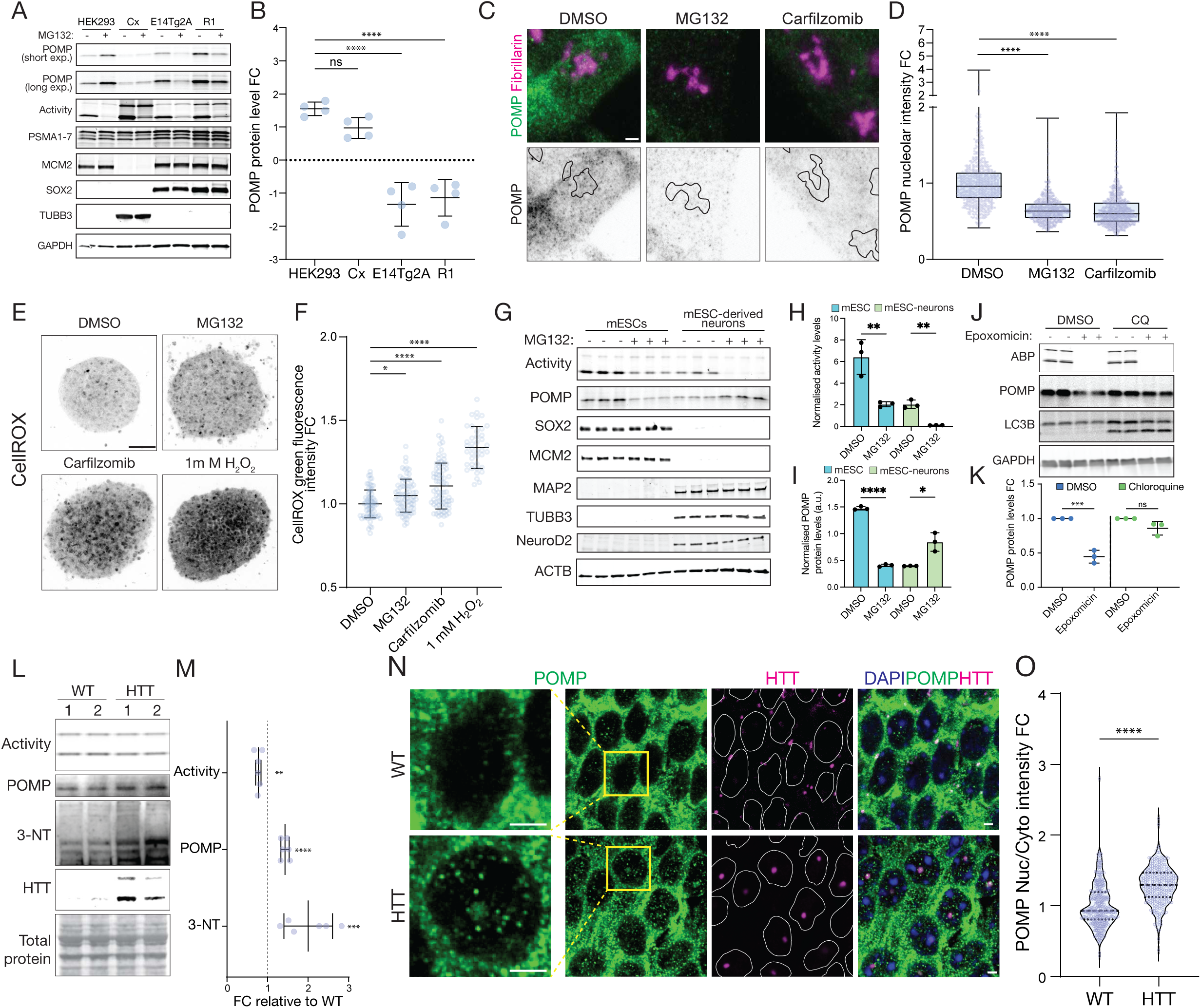
POMP’s nucleolar function is differentially regulated in development and disease. (A) Western blot analysis of HEK293, rat cortical neurons, E14Tg2A and R1 mouse embryonic stem cells treated with DMSO or MG132 for 5 hr. All cells showed a robust decrease in proteasome activity, measured by the in-gel activity-based probe (ABP) fluorescence, in response to MG132 treatment. While HEK293 and rat cortical neurons respond by increasing POMP levels, mESCs reduce POMP levels independently of the proteasome (PSMA3). Stainings for the mitotic marker MCM2, the pluripotency marker SOX2 and the neuronal marker TUBB3 demonstrate the identity of the cells assayed. (B) Analysis of experiments like the one shown in A. While HEK293 and cortical neurons respond to proteasome inhibition with a significant increase in POMP levels, mESCs display a significant reduction in POMP levels. ****p<0.0001, one-way ANOVA with post-hoc Dunnett’s test, n=4, mean±SD. FC=fold change. (C) Representative high-magnification immunofluorescence images of E14Tg2A mESCs treated with either DMSO or proteasome inhibitors (MG132, Carfilzomib) for 5 hr, and stained for POMP and the nucleolar marker fibrillarin. The images show a decrease in POMP levels across all cell compartments, including the nucleolus. Traced in black in the POMP single channel image are the nucleolar outlines. Scale bars = 2 µm. (D) Analysis of POMP nucleolar intensity in experiments like the one shown in C. In response to proteasome inhibition, cells significantly reduce global cellular POMP levels, including the nucleolus. ****p<0.0001, one-way ANOVA with post-hoc Dunnett’s test, n=715 (DMSO), 616 (MG132), 796 (Carfilzomib), mean±SD. FC=fold change. (E) Representative immunofluorescence images of E14Tg2A mESC colonies treated with DMSO or proteasome inhibitors (MG132, Carfilzomib) for 5 hr, and 1 mM H₂O₂ for 1h as a positive control. Proteasome inhibition leads to an increase in CellROX green fluorescence. Scale bars = 50 µm. (F) Analysis of experiments like the one shown in E. Proteasome inhibition leads to a significant increase in cellular oxidative stress levels. *p≤0.05, ****p<0.0001, one-way ANOVA with post-hoc Dunnett’s test, n=69 (DMSO), 74(MG132), 75(Carfilzomib) and 40 (H₂O₂), mean±SD. FC=fold change. (G) Western blot analysis of mESC and mESC-derived cortical neuron cultures treated with DMSO or MG132 for 5 hr. While the mitotic (MCM2) and pluripotency markers (SOX2) could be detected only in the mESCs, neuronal markers (MAP2, TUBB3, NeuroD2) are present only after differentiation. While mESCs respond to proteasome inhibition by downregulating POMP, after differentiation mESC-derived neurons show increased POMP levels in response to proteasome inhibition. Both cell types display a strong reduction in proteasome activity measured by in-gel ABP fluorescence. (H) Analysis of the proteasome activity levels measured by in-gel ABP fluorescence shown in G. MG132 treatment significantly reduces proteasome activity in mESCs and mESC-derived neurons. **p<0.01, unpaired two-tailed t-tests, n=3, mean±SD. (I) Analysis of the POMP protein levels’ changes shown in G. Proteasome inhibition leads to contrasting responses in POMP levels in mESCs and mESC-derived neurons. *p≤0.05, ****p<0.0001, unpaired two-tailed t-tests, n=3, mean±SD. (J) Western blot analysis of mESCs treated with either water or the autophagy inhibitor chloroquine for 2 hr prior to addition of DMSO or the proteasome inhibitor epoxomicin for 5 hr. In gel ABP fluorescence shows a near complete inhibition of proteasome activity in the presence or absence of chloroquine. Chloroquine treatment leads to accumulation of the lower molecular weight lipidated form of LC3B (LC3B-II), reflecting efficient autophagic flux blockage, and prevents the decrease in POMP levels induced by proteasome inhibition. (K) Analysis of experiments like the one shown in J. Inhibition of autophagic degradation with chloroquine prevents degradation of POMP in response to proteasome inhibition in mESCs. ns=p>0.05, ***p<0.001, unpaired two-sample t-tests, n=3, mean±SD. (L) Western blot analysis of hippocampal native extracts from WT and mice expressing mutant huntingtin (HTT). The samples were assayed for proteasome activity levels by in-gel ABP fluorescence, the levels of POMP, HTT and the oxidative-stress marker 3-NT. HTT mouse extracts display elevated levels of HTT, a subtle reduction in proteasome activity and elevated levels of POMP and 3-NT. (M) Analysis of experiments like the one shown in L. Hippocampi of the HTT mouse model displayed a modest, but significant reduction in proteasome activity with a concomitant significant increase in POMP and 3-NT levels. **p<0.01, ***p<0.001, ****p<0.0001, unpaired two-sample t-tests, n=7 (WT) and 6 (HTT) mice, mean±SD. (N) Representative high-magnification immunofluorescence images of WT and HTT mouse hippocampal neurons’ somata stained for HTT and POMP. In the Huntington’s disease mouse model accumulation of HTT as nucleolar puncta is visible, and POMP also displays nuclear enrichment and accumulation in the HTT^+^ nucleoli. Traced in white in the HTT single channel image are the nuclear outlines. Scale bars = 2 µm. (O) Analysis of images like the ones shown in N. The levels of POMP were measured in the nuclei and cytoplasm of hippocampal and cortical neurons of WT and HTT mice and the nuclear-cytoplasmic ratio of the signal was calculated. Neurons of the Huntington’s disease mouse model display a significant relative enrichment in nuclear POMP compared to the WT control. ****p<0.0001, unpaired two-sample t-test, n= 342 (WT) and 273 (HTT) from 6 different animals. Violin plots display median, interquartile range, minimum and maximum values, FC=fold change.

Our KEGG pathway analysis indicated that POMP accumulation (prompted by PSMB knockdown), is associated with transcriptional signatures of neurodegenerative conditions (Figure S6A j). We tested whether nucleolar relocalisation of POMP could be dysregulated in the context of disease. We used a Huntington’s disease mouse model^66^ (HTT) and examined proteasome activity and POMP levels. We found that proteasome activity showed a modest but significant reduction and that the levels of POMP and nitrotyrosine^67^ (a marker for oxidative damage) were significantly elevated (Figure 7l,m). We next examined the presence of POMP puncta in the nucleus of healthy vs. HTT diseased neurons (Figure 7n). Immunofluorescence analysis revealed a significant accumulation of POMP+ puncta in the nucleus of HTT neurons (Figure 7 n,o). Taken together, our data suggest that relocalisation of POMP to the nucleus and nuclear bodies of cells is part of a protective cell-stress response that gets activated in a number of biological settings and that certain cells counter by engaging autophagy to reduce POMP levels.

## Discussion

Here we used a combination of cell types to examine proteostatic mechanisms that are invoked upon cellular stress, focusing on the proteasome chaperone POMP and manipulations that compromise proteasome activity. We found that in mammalian cells when proteasome function is impaired, the normally short-lived assembly chaperone POMP accumulates and exhibits an HSF1-dependent increase in transcription. Proteasome inhibition also leads to an increase in reactive oxygen species, which is necessary for POMP translocation and retention in the nucleolus. Within the nucleolus, re-localized POMP acquires a new set of interaction partners including RNA-processing factors like Upf1 and TDP-43, suggesting a role in RNA regulation. Transcriptomic experiments that functionally dissected the proteasome-independent actions of POMP, in combination with targeted gain- and loss-of-function experiments, further revealed that POMP promotes a protective transcriptional program that prevents induction of a pro-inflammatory and pro-apoptotic axis. POMP’s nucleolar relocalization was associated with a reprogramming of cellular metabolism, promoting ribogenesis and compensatory transcription of all proteasome subunits. This ‘hard-wired’ pro-survival mechanism is developmental stage-dependent; while embryonic stem cells prevent its activation by engaging the autophagy pathway to degrade POMP, differentiated neurons no longer have this ability. Furthermore, we found that this noncanonical function of POMP is activated in the context of neurodegenerative disease, where proteasome dysfunction, inflammation and oxidative stress converge^2,68–70^.

The observation that, in response to proteasome dysfunction, POMP interacts with RNA biogenesis factors in the nucleolus to shape the transcriptional response may help explain several long-standing questions in the field. Unlike yeast cells which, in response to loss of the proteasome biogenesis factor Ump1, exhibit a robust “bounce-back effect” and upregulate proteasome subunits’ levels to maintain proteasomes^10,27,30^, we observed that mammalian cells lacking (the Ump1 homolog) POMP fail to compensate and instead exhibit a reduction in assembled cellular proteasomes. This failure is despite the ability of mammalian cells to increase expression of proteasome subunits through NRF1-driven compensatory transcription in response to proteasome dysfunction^11,12^. We have shown that, in response to the same level of proteasome impairment, POMP-deficient cells show a reduced ability to induce CP subunit transcripts. In addition, consistent with a previous report^37^, we found that overexpression of POMP is sufficient to increase the cellular levels of proteasome α (PSMA1-7) and β5 (PSMB5) subunits. Altogether, these data suggest that nuclear POMP serves as a critical facilitator of NRF1-dependent CP subunit transcription. Previous work has shown that POMP itself is induced through NRF1-driven transcription to promote proteasome biogenesis; however, this effect was observed after prolonged (10-16 hrs) periods of proteasome inhibition^11,12^. We found that earlier in the cellular response to proteasome inhibition, HSF1, the master regulator of the heat-shock response, drives POMP transcription. HSF1 may be more broadly implicated in the regulation of POMP levels, as it was also activated in response to cytokine treatment.

Building on these findings, we observed that when POMP becomes stoichiometrically uncoupled from its canonical interactors (the proteasome core particle subunits), which can occur in the early phases of proteasome inhibition, in response to cytokine signalling, in inflammation and disease, it takes on a new function and regulates the transcriptional output of cells. Additional evidence supporting a non-canonical moonlighting function of POMP comes from its localization and structural properties. The intrinsically unstructured nature of POMP aligns with its potential localization to the nucleolus and nuclear bodies^17^. While POMP chaperones proteasome assembly on intracellular vesicles in mammalian cells, it has also been detected by others in the nucleus and nuclear bodies^46^. In addition, it was identified in a screen for nuclear quality control factors^71^ and a recent study, mapping human subcellular architecture, classified it in the splicing speckle assembly^72^. AI-based structural predictions^73^ and empirical data^53^ suggest that POMP lacks a defined tertiary structure, and cryo-EM studies^74–76^ have shown that during proteasome assembly it undergoes significant conformational changes as it interacts with incoming β-subunits. This structural adaptability lends support to the data we show here that POMP forms new interactions to sense and resolve disturbances in cellular protein homeostasis.

Our data indicate that under conditions of proteasome inhibition, POMP accumulates in the nucleolus and contributes to a pro-survival gene expression program. POMP has been previously identified as a vulnerability factor in proteasome inhibitors–resistant tumour cells^77^. These effects of POMP were thought to occur exclusively through its well-documented effect on proteasome biogenesis^77^; however, our data suggest that its proteasome-independent protective transcriptional function may also play a role. The fact that ESCs have a strategy to prevent the accumulation of POMP and its nucleolar translocation, which reverses upon differentiation, attests to the importance of this mechanism for cell fate decisions. It is possible that in cells critical for healthy organismal development (like stem cells) it is judicious to swiftly eliminate cells with an impaired proteostatic network in order to preserve tissue integrity and function^78–80^. Conversely, terminally differentiated cells that cannot be readily replaced, such as neurons, may rely on the POMP pro-survival mechanism to manage proteostatic stress, retain functional integrity and withstand age- and disease-associated challenges^81–85^. This fits with our observation that, in a mouse model of Huntington’s disease^86^, POMP levels are elevated and it localizes to the nucleoli of cortical and hippocampal neurons. This process appears to be facilitated by elevated oxidative stress^69^—evidenced by increased protein tyrosine nitration—and may represent an effort to protect cells from proteostatic collapse.

Conceptually, our work sheds new light on mechanisms that have been observed before, but whose functional relevance remains poorly understood. For instance, the nucleolus is known to be a stress-sensing hub^13,15^ into which stress-responsive transcription factors (e.g. p53^87^, RelA^88^ and c-Jun^20^) translocate in response to stimuli such as DNA damage, hypoxia and proteotoxic stress. However, their roles within the nucleolus are not yet fully elucidated. Our data show that POMP, which under basal conditions is almost exclusively cytoplasmic, under stress can act as a coincidence detector that integrates stimuli (i.e. proteasome activity, oxidative stress, inflammation and cytokine signalling) and then translocates into nucleolus to modulate gene and transcript isoform expression. Mutations and alterations in POMP levels have been implicated in the etiology of different immunodeficiency^89^, autoinflammatory and autoimmune diseases^34,54^ through its role in proteasome assembly. Our data indicate that POMP coordinates a previously unidentified signalling axis that may be at play in a number of scenarios of altered proteostasis. More broadly, our work illustrates how the cell can program a direct response to pathway perturbations by endowing specific assembly factors with context-dependent moonlighting functions - encoding within the pathway a mechanism that operates in parallel with, and fine-tunes, other stress-response cascades.

## Resource availability

### Lead contact

Further information and requests for resources and reagents should be directed to and will be fulfilled by the Lead Contact, Erin M. Schuman.

### Materials availability

All unique materials generated in this study are available upon request from the lead contact.

## Methods

### Mammalian cell culture

HEK293 (CRL-1573) were obtained from ATCC and maintained in DMEM+Glutamax (Invitrogen, 10566016) supplemented with 10% FBS (Invitrogen, 16141079). ES-E14Tg2A (CRL-1821) and R1/E (SCRC-1036) mouse embryonic stem cells (mESCs) were obtained from ATCC and adapted to feeder-free culture on 0.1% gelatin-coated plates (Sigma-Aldrich, G1890) in DMEM+Glutamax (Invitrogen, 10566016) supplemented with 0.001% β-mercaptoethanol (Sigma-Aldrich, M6250), 10% FBS (Invitrogen, 16141079), MEM amino acids (Invitrogen, 11130051) and 103 U/ml of ESGRO Recombinant Mouse LIF Protein (Sigma-Aldrich, ESG1106). HEK293 and mESCs were grown at 37°C in 5% CO2, passaged weekly with TrypLE Express (Invitrogen, 12605028) and regularly mycoplasma tested. For imaging purposes HEK293 and mESCs were seeded onto MatTek dishes (MatTek, P35GC-1.5-14-C) coated with 0.1 mg/ml poly-D-Lysine (PDL) (Invitrogen, A3890401) or 0.1% gelatin (Sigma-Aldrich, G1890), respectively. Dissociated rat primary cortical neuron cultures were prepared and maintained as previously reported^90^. Briefly, cortices and hippocampi from postnatal day 0-1 rat pups of either sex (Sprague Dawley, IGS, Crl:CD(SD), Charles River Laboratories, RRID:RGD_734476) were dissected, digested with Papain (Sigma, P3125), and plated at densities of ∼3-4×10^4^ cells per imaging dish (MatTek, P35G-1.5-14-C,) or 4×10^6^ cells per 10 cm dish for biochemistry. All culture dishes used for neuronal culture were coated with 0.1 mg/ml poly-D-lysine (Invitrogen, A3890401). After 4 hr, 500 μl glia-conditioned medium was added and the cells were maintained for up to 30 DIV at 37°C in 5% CO2 with weekly feedings with 700 μl of normal growth media [NGM; Neurobasal A (Invitrogen, 10888-022), B27 (Invitrogen, 17504-044), GlutaMAX (Invitrogen, 035050-038)].

### Yeast cell culture

Parental (BY4743) and Ump1-/- (Accession: YBR173C, Clone ID: 33313) S. cerevisiae strains were purchased from Horizon streaked out and cultured on YPD agar plates (Sigma-Aldrich, Y1500), supplemented with 200 µg/mL G418 (Sigma-Aldrich, A1720) in the case of the Ump1-/- mutant. Starter cultures were inoculated from single colonies and grown overnight in liquid medium. For experimental cultures, overnight precultures were diluted to an optical density at 600 nm (OD600) of 0.1–0.2 and grown to mid-log phase (OD600∼0.5–0.7).

### Yeast genotyping

Genomic DNA was extracted from parental (WT) and Ump1-/- strains using the YeaStar Genomic DNA Kit (Zymo Research, D2002) and the following primers were used for PCR genotyping: primer_A- CATGTGATGTGACTAGTGTTTGTGA, primer_B-TTTAAAGGTATCTTGTGGGACGATA, primer_C- CACCGTGACATACTACTGAACAAAG, primer_D-TGGGCTGAGAAGTTGAGTATATAGG, primer_E- TGCAGTGCCCTCCTTGCC, primer_F-AGGGTAGACGTCCTCCCAATCGAT, Kan-F- CTGCGATTCCGACTCGTCCAACA, Kan-R-ACTAAACTGGCTGACGGAATTTATGCCTC. Genotyping PCR was performed using GoTaq Green master mix (Promega, M7122) a Ta of 57°C and 1.5 min as extension time.

### siRNA transfection

Pre-designed siRNA oligo duplex sets against POMP (SR309742), TC11/NRF1 (SR303155), HSF1 (SR320556), PSMB1 (SR303824) and PSMB7 (SKU SR303830) were purchased from OriGene. The siRNAs were resuspended in OriGene’s duplex buffer at a concentration of 20 µM, heated to 94°C and allowed to cool to room temperature to ensure efficient oligo annealing. HEK293 cells were transfected with siRNAs at a concentration of 10 nM with Lipofectamine RNAiMax following the vendor’s instructions. Briefly, siRNAs and Lipofectamine RNAiMax were mixed (1:1) in OptiMEM reduced serum medium (Invitrogen 31985062) and incubated at room temperature for 5 min before adding to 50-60% confluent HEK293 cells. The next day the cells were fed with fresh medium (1:1 old and fresh medium) and grown for two more days. Gene knockdown and phenotypic effects were assayed 72h post-transfection.

### Plasmid cloning

Plasmids pCMV6-Entry (PS100001), pCMV6-POMP-Myc-Flag (SKU RC204812, POMP-Myc-Flag) and the pGFP-C-shLenti kit against R. norvegicus POMP (TL704218) were purchased from OriGene. The constructs HA-POMP (86765), pHBS1389 IBB-GFP-mCherry3E (118803) and RG6 (80167) were purchased from Addgene. The construct pCMV-XL5-POMP (POMP) was generated by restriction ligation using NotI (Invitrogen, FD0596) to open the pCMV-XL5 (OriGene, PCMV6XL5) plasmid and to generate sticky ends in the POMP ORF amplified from the HA-POMP construct. The following constructs were generated by linearizing pcDNA3.1(-) (Invitrogen, V79020) by restriction digestion and cloning in gBlocks (IDT) by Gibson assembly (NEB, E2611L): pcDNA3.1-HA-POMP-Myc-Flag (HA-POMP-Myc-Flag), pcDNA3.1-HA-Scarlet (HA-Scarlet), pcDNA3.1-NLS-HA-Scarlet (NLS-HA-Scarlet), pcDNA3.1-HA-POMP-Scarlet (POMP-Scarlet), pcDNA3.1-NLS-HA-POMP-Scarlet (NLS-POMP-Scarlet), pcDNA3.1-NoTS-HA-Neon (NoTS-Neon) and pcDNA3.1-NoTS-HA-POMP-Neon (NoTS-POMP-Neon). For the splicing reporter assays the mNeon CDS in the pcDNA3.1-NoTS-HA-Neon (NoTS-Neon) and pcDNA3.1-NoTS-HA-POMP-Neon (NoTS-POMP-Neon) constructs the was replaced with mTagBFP2 by linearizing the backbone by PCR and cloning an mTagBFP2 gBlock (IDT) by Gibson assembly. The viral construct used to generate AAVs for transduction of cortical neurons (pAAV-CMV-HA-POMP-Myc-Flag) was generated by restriction digestion using MluI (Invitrogen, FD0564) and EcoRV (Invitrogen, FD0304) to linearize the construct pAAV-hSyn-GFP (Addgene, 50465), and replace by ligation the hSyn-GFP cassette with the CMV-HA-POMP-Myc-Flag insert excised from pcDNA3.1-HA-POMP-Myc-Flag. The mutation C36A was introduced into the pcDNA3.1-NLS-HA-POMP-Scarlet by site directed mutagenesis (NEB, E0554S). Gel extraction and PCR purification was done using standard kits from Qiagen ad Zymo Research. DNA ligation was performed using T4 ligase (NEB, M0202L) and for all cloning steps involving amplification Q5 high-fidelity polymerase (NEB, M0492S) was used. The reactions were transformed into high-efficiency NEB 5-alpha (NEB, C2987H), NEB stable (NEB, C3040H) and TOP10 (NEB, C404003) chemically competent E. coli plated antibiotic selection plates. Colonies were screened either by restriction digestion or colony PCR and validated by Sanger sequencing. All constructs were stored as glycerol stocks at −80°C. Bacteria were grown under selection in terrific broth (Invitrogen, 22711022) and plasmids were prepped using Qiagen plasmid plus midi kits and endofree maxi kits.

### Plasmid transfection

Hippocampal neurons were transfected at 7 DIV using Lipofectamine 2000 (Invitrogen, 11668019) with a final plasmid concentration of 250 ng/ml. Briefly, for each transfection, 1.5 μL of Lipofectamine 2000 was combined with 75 μL of Neurobasal medium supplemented with GlutaMAX and incubated at room temperature for 5 minutes. In a separate tube, 500 ng of plasmid DNA was mixed with 75 μL of the same supplemented Neurobasal medium. The Lipofectamine-containing solution was then added to the DNA solution, gently mixed, and incubated at RT for 20 minutes to allow complex formation. Meanwhile, the culture medium of the neurons was carefully removed and preserved. A total of 150 μL of the transfection mixture was then applied to the neurons cultured on MatTek dishes, followed by incubation at 37°C in a humidified 5% CO₂ atmosphere for 1 hour. After incubation, the transfection mixture was aspirated, and the original culture medium was added back. The cells were fixed and processed for downstream immunofluorescence analysis 8 days post-transfection. HEK293 cells were transfected using either TransIT-293 (Mirus bio, MIR 2704), as outlined in the product’s manual, or 1 mg/ml PEI ̴̴pH 7 (Sigma-Aldrich, 408727), as previously described^91^. Briefly, HEK293 cells were seeded 24-48 hr before transfection to allow them to attach and reach 60-70% confluency. Transfection was performed using a DNA:transfection reagent ratio of 1:3. First, DNA (0.5-4 µg/ml) and then transfection reagent were added to 100 µl of OptiMEM reduced serum medium, the mixture was briefly vortexed and incubated for 10 minutes at room temperature to allow complex formation. The transfection mixture was then added dropwise to the cells. The next day the cells were fed with fresh medium (1:1 old and fresh medium) and grown for two more days. Phenotypic effects were assayed 72h post-transfection.

### AAV packaging and transduction

Packaging of the pAAV-CMV-HA-POMP-Myc-Flag construct was outsourced to VectorBuilder and used the AAV-DJ pilot-scale packaging service, which included the EGFP AAV-DJ control virus. Transduction was carried out on DIV 7 cortical neurons plated on 10 cm PDL-coated dishes (4×106 cells/dish) with an MOI of 125, based on viral genome copies (GC/cell). The cells were cultured until DIV 28 with weekly feedings. The pENN.AAV.CamKII.GCaMP6f.WPRE.SV40 virus was obtained from Addgene (100834-AAV1) and was used to transduce mESC-derived cultures one week after organoid dissociation with an MOI of 100, based on viral genome copies (GC/cell).

### SDS-PAGE analysis

For reducing SDS-PAGE analysis, mammalian cells were washed three times in DPBS+Ca^2+^/Mg^2+^ (Invitrogen, 14040141) and lysed by scraping in RIPA buffer (Invitrogen, 89900) supplemented with Halt protease and phosphatase inhibitor cocktail (Invitrogen, 78440). Cell extracts were snap-frozen, thawed, centrifuged at 12×103 g for 5 min at 4°C and protein concentration was determined using the Precision Red advanced protein assay. Equal amounts of protein (10-70 μg) were mixed with NuPAGE LDS sample buffer (Invitrogen, NP0007), NuPAGE sample reducing agent (Invitrogen, NP0009) and water to the same final concentration, and incubated at 85°C for 5 min in a thermomixer before electrophoresis. For non-reducing SDS-PAGE analysis, mammalian cells were washed three times in DPBS+Ca^2+^/Mg^2+^ (Invitrogen, 14040141) and lysed by scraping in urea buffer pH 8 (8M urea, 75 mM NaCL, 4% CHAPS, 200 mM Tris-HCl pH 8.4) supplemented with Halt protease and phosphatase inhibitor cocktail (Invitrogen, 78440). Cell extracts were clarified by centrifugation at 12×103 g for 5 min at room temperature and protein concentration was determined using the Precision Red advanced protein assay. Equal amounts of protein (10-70 μg) were mixed with ROTI Load 3 (Carl Roth, 3359.1) and distilled water to the same final concentration and incubated at room temperature for 1h before electrophoresis. For protein gel electrophoresis Novex Tris-Glycine (Invitrogen), Novex Bis-Tris (Invitrogen) and Criterion TGX (Bio-Rad) pre-cast gels varying concentrations were used (i.e. 10, 12, 4-12, 4-20%) depending on the protein target. The gels were run in either NuPAGE MOPS SDS running buffer (Invitrogen, NP0001) or Tris-Glycine SDS running buffer (25 mM Tris, 192 mM glycine, and 0.1% SDS, pH 8.3) for 1-2 hr at 100-150 V. Proteins were transferred onto nitrocellulose by semi-dry transfer using the Trans-Blot Turbo system (Bio-Rad), mini (Bio-Rad, 1704158) and midi (Bio-Rad, 1704159) transfer packs. The membranes were stained as described in the Western blot section of the Methods.

### Native PAGE analysis of mammalian cells and in gel proteasome activity profiling

Mammalian cells were washed three times in DPBS+Ca^2+^/Mg^2+^ (Invitrogen, 14040141) and lysed by scraping in HR buffer^92^ (50 mM Tris, 5 mM MgCl2, 250 mM sucrose, 1 mM DTT and 2 mM ATP, pH=7.4) and then passed through a 27G syringe needle for ten times. The extract was clarified by centrifugation at 10^4^ g for 10 min at 4°C and protein concentration was determined using the Precision Red advanced protein assay. Equal amounts of protein (80-100 μg) were supplemented with 10X native sample buffer (0.05% (w/v) bromophenol blue, 50%glycerol, Tris 500 mM pH 7.5) and gently mixed by pipetting. Native gel electrophoresis was performed in 3–8% Criterion™ XT Tris-Acetate Protein Gels (Bio-Rad), pre-run for 1h at 30V at room temperature in TBE running buffer (Invitrogen, J62788.K2) supplemented with 5 mM ATP pH7, 0.5 mM DTT and 12.5 mM MgCl2. For protein electrophoresis, the gel tank was moved to the cold room and run at 30 Amp for ∼3h. After the run, the gel was incubated in 20 ml of peptidase assay solution (50 mM Tris-HCl pH 7.5, 5 mM MgCl2, 1 mM ATP pH 7 and 0.02% SDS) supplemented with 50 µM Suc-LLVY-AMC fluorogenic substrate (Abcam, ab142120) for 20-40 min at 37°C and room temperature for mammalian and yeast samples, respectively. After visualization of the in-gel fluorogenic substrate signal on an Azure 280 system, proteins were transferred onto nitrocellulose membranes (Bio-Rad, 1620112) by wet-transfer in Tris-Glycine transfer buffer (25 mM Tris, 192 mM Glycine, 20% (v/v) methanol, pH ∼8.3) in the cold room at 20V overnight. The membranes were stained as described in the Western blot section of the Methods.

### Western blot analysis

The membranes were Ponceau stained (Cell Signalling, 59803), destained and blocked for 30 min at room temperature in Intercept PBS blocking buffer (Li-Cor Bio, 927-70001) or TBS blocking buffer (Li-Cor Bio, 927-60001), in the case of phosphorylated targets. The membranes were probed overnight with primary antibodies, washed three times with TBST, stained with fluorescently-labelled or HRP-linked secondaries diluted 1:3000 in blocking buffer for 30 min at room temperature. After three washes in TBST and two additional washes in TBS, the fluorescent or signal was acquired using either the Azure Sapphire FL RGB or the Li-Cor Odyssey 9120 gel imaging system. For chemiluminescence detection the HRP-stained membranes were developed using the SuperSignal West Femto (Invitrogen, 34094) or Dura (Invitrogen, 34075) substrates and the chemiluminescence signal was visualized on the Azure 280 system.

### Immunofluorescence staining

Cells were incubated in glyoxal fixative^93,94^ (7.8% (v/v) glyoxal, 19.7% (v/v) ethanol, 0.75% acetic acid, pH adjusted to 4-5 with 1M NaOH) for 30 min at room temperature, permeabilized for 15 min (0.5% Triton-X-100 (v/v) 4% goat/donkey serum (v/v) in PBS pH 7.4) and blocked for 1h (4% goat/donkey serum (v/v) in PBS) before incubation with primary antibodies diluted in blocking buffer overnight at 4°C. The next day the cells were washed three times (5min) in DPBS + Ca²⁺/Mg²⁺ (Invitrogen, 14040141) and stained with secondary antibodies (DF=1:500) and DAPI (0.1 µg/mL) in blocking buffer for 1h at room temperature. Cells were washed three times (5 min each) with DPBS + Ca²⁺/Mg²⁺ (Invitrogen, 14040141), then either mounted with Aqua-Poly/Mount (Polysciences, 18606-20) for long-term storage or kept in PBS for immediate imaging.

### Antibodies

Primary antibodies used for protein detection, with their corresponding dilutions for immunofluorescence (IF) and Western blotting (WB) were: mouse anti-PSMA1-7 (Enzo, BML-PW8195, WB 1:1000), mouse anti-PSMA1 (Enzo, BML-PW9390, WB 1:1000), mouse anti-PSMC3 (Enzo, BML-PW8770, WB 1:1000), PSMA3 (Enzo, BML-PW8110, IF 1:200, WB 1:1000), rabbit anti-PSMB5 (CST, 12919, WB 1:1000), rabbit anti-POMP (CST, 15141, IF 1:200, WB 1:1000), mouse anti-GAPDH (Sigma-Aldrich, G8795, WB 1:3000), rabbit anti-ACTB (Abcam, ab8227, IF 1:1000, WB 1:3000), rabbit anti-TCF11/NRF1 (CST, 8052, WB 1:1000), mouse anti-Lamin B1 (Abcam, ab8982, WB 1:2000), mouse anti-Hsp70 (Abcam, ab2787, WB 1:2000), rabbit anti-HSF1 (CST, 4356, WB 1:1000), rabbit anti-HSF1 (phospho S326) (Abcam, ab76076, WB 1:1000), mouse anti-Calnexin (Proteintech, 66903-1-Ig, IF 1:200, WB 1:1000), rabbit anti-Calreticulin (Abcam, ab92516, IF 1:200), rabbit anti-Histone H3 (Abcam, ab1791, WB 1:3000), mouse anti-GM130 (BD Biosciences, 610823, WB 1:1000), mouse anti-NCAD (BD Biosciences, 610921, WB 1:1000), rabbit anti-ERGIC53 (Sigma-Aldrich, E1031, WB 1:1000), rabbit anti-EEA1 (Abcam, EPR4245, WB 1:1000), rabbit anti-TOMM70 (Proteintech, 14528-1-AP, WB 1:1000), rabbit anti-VAMP2 (Proteintech, 10135-1-AP, WB 1:1000) mouse anti-ATG5 (Proteintech, 66744-1-Ig, IF 1:200, WB 1:1000), mouse anti-ATP6V1H (Proteintech, 68425-1-Ig, IF 1:200, WB 1:1000), rabbit anti-ATP6V1G1 (Proteintech, 16143-1-AP, WB 1:1000), rabbit anti-Sec13 (Proteintech, 15397-1-AP, WB 1:1000), chicken anti-MAP2 (Abcam, ab5392, IF 1:2000, WB 1:1000), mouse anti-Fibrillarin (Abcam, ab4566, IF 1:500), rabbit anti-Rps3 (Invitrogen, A303-840A, WB 1:1000), rabbit anti-Rpl24 (Sigma-Aldrich, SAB2700500, WB 1:1000), rabbit anti-Upf1 (Sigma-Aldrich, HPA019587, IF 1:200), rabbit anti-PSMB1 (Proteintech, 11749-1-AP, WB 1:1000), rabbit anti-PSMB7 (Proteintech, 30283-1-AP, WB 1:1000), anti-mouse α-Tubulin (Sigma-aldrich, T6199, WB 1:3000), anti-rabbit p62/ SQSTM1 (CST, 5114, IF 1:500, WB 1:1000), rabbit anti-PSMG2 (Proteintech, 10959-1-AP, WB 1:1000), rabbit anti-HA tag (CST, 3724S, WB 1:1000, IF 1:500), rabbit anti-Flag tag (CST, 2368P, WB 1:1000), mouse anti-Myc tag (CST, 2276S, IF 1:200, WB 1:1000), mouse anti-β-Tbulin III (Sigma-Aldrich, T5076, WB 1:1000), mouse anti-Bassoon (Enzo, ADI-VAM-PS003-D, IF 1:500), mouse anti-vGlut1 (SYSY, 135011, IF 1:500), rabbit anti-GFAP (Abcam, ab7260, IF 1:200, WB 1:1000), mouse anti-GAD65/67 (Enzo, ADI-MSA-225-E, IF 1:500), rabbit anti-MBP (Abcam, ab40390, IF 1:1000), rabbit anti-c-Fos (CST, 2250, WB 1:1000), rabbit anti-cleaved Caspase7 (CST, 9491, WB 1:1000), mouse anti-cleaved PARP (CST, 9546, WB 1:1000), Phospho-Histone H2A.X (CST, 9718, WB 1:1000), rabbit anti-PSMB10 (Proteintech, 15976-1-AP, WB 1:1000), mouse anti-PSMB3 (Abnova, H00005691-B01P, WB 1:500) rabbit anti-PSMB4 (Proteintech, 11029-1-AP, WB 1:1000) rabbit anti-PSMB2 (Proteintech, 15154-1-AP, WB 1:1000), rabbit anti-PSMB8 (Proteintech, 14859-1-AP, WB 1:1000), rabbit anti-p21 Waf1/Cip1 (CST, 2947, WB 1:1000), rabbit anti-JunB (CST, 3753, WB 1:1000), goat anti-Rpl22 (LSBio, 58905, IF 1:200), rabbit anti-RpsA (Proteintech, 14533-1-AP, IF 1:200), rabbit anti-Rpl30 (Proteintech, 17403-1-AP, IF 1:200), rabbit anti-Rps23 (Proteintech, 29834-1-AP, IF 1:200), rabbit anti-Rps3A (Proteintech, 14123-1-AP, IF 1:200), rabbit anti-Rpl34 (Proteintech, 15179-1-AP, IF 1:200), mouse anti-HA tag (Roche, 10952100 IF 1:200), rabbit anti-ABCE1 (Sigma-Aldrich, HPA036846, IF 1:), rabbit anti-Eif4e (CST, 2067, IF 1:100), rabbit anti-Eif4g1 (Invitrogen, 16197544, IF 1:100), rabbit anti-Tdp43 (Proteintech, 10782-2-AP, IF 1:100), mouse anti-SQSTM1/p62 (Proteintech, 66184-1-Ig, IF 1:100), rabbit anti-SQSTM1/p62 (CST,5114, WB 1:1000), rabbit anti-LC3 (CST, 2775, WB 1:1000), rabbit anti-SOX2 (Proteintech, 11064-1-AP, WB 1:1000), rabbit anti-MCM2 (CST, 4007, WB 1:1000), mouse anti-α-tubulin (Sigma-Aldrich, T6199, WB 1:3000), mouse anti-β-tubulin III (Sigma-Aldrich, T8578, WB 1:3000), rabbit anti-nitrotyrosine (3-NT) (Invitrogen, A-21285, WB 1:1000). The fluorescently-linked secondary antibodies used in the study were were: Alexa Fluor 405 goat anti-chicken IgY (Abcam, ab175674), Alexa Fluor 405 goat anti-mouse IgG (Invitrogen, A31553), Alexa Fluor anti-mouse IgG2a (Sigma-Aldrich, SAB600476), Alexa Fluor 405 anti-mouse IgG1 (Sigma-Aldrich, SAB4600474, IF 1:500), Alexa Fluor 488 goat anti-rabbit IgG (Invitrogen, A11008), 488 goat anti-mouse IgG (Invitrogen, A11001), Alexa Fluor 488 goat anti-mouse IgG2b (Invitrogen, A21141) Alexa Fluor 488 goat anti-chicken IgY (Invitrogen, A11039), Alexa Fluor 568 goat anti-rabbit IgG (Invitrogen, A11011), Alexa Fluor 568 goat anti-mouse IgG (Invitrogen, A11004, IF 1:500), Alexa Fluor 568 goat anti-mouse IgG1 (Invitrogen, A21124), Alexa Fluor 568 goat anti-mouse IgG2a (Sigma Aldrich, SAB4600315), Alexa Fluor 568 anti-chicken IgY (Invitrogen, A11041), Alexa Fluor 568 donkey anti-goat IgG (Invitrogen, A11057), Alexa Fluor 647 goat anti-mouse IgG2a (Invitrogen, A21241), Alexa Fluor 647 goat anti-rabbit IgG (Invitrogen, A21245), Alexa Fluor 647 goat anti-mouse IgG (Invitrogen, A21236), Alexa Fluor 647 goat anti-chicken IgY (Invitrogen, A32933TR), IRDye 680LT goat anti-mouse IgG (Li-Cor, 926-68020),), IRDye 680RD goat anti-rabbit IgG (Li-Cor, 926-68071), IRDye 680LT goat anti-mouse IgG1 (Li-Cor, 926-68050), IRDye 800CW goat anti-rabbit IgG (Li-Cor, 926-32211), IRDye 800CW goat anti-mouse IgG (Li-Cor, 926-32210), IRDye 800CW goat anti-mouse IgG2a (Li-Cor, 926-32351), IRDye 800CW goat anti-mouse IgG2b (Li-Cor, 926-32352), IRDye 800CW donkey anti-chicken IgG (Li-Cor, 926-32218), HRP-linked anti-rabbit IgG (CST, 7074), HRP-linked anti-mouse IgG (CST, 7076). For immunofluorescence imaging the secondary antibodies were used at a 1:500 dilution, for western blot they were used at a 1:3000 dilution. HRP-linked secondaries were diluted in 5% milk as blocking buffer.

### Fluorescence-Based Proteasome Activity Profiling

Proteasome activity profiling with the activity-based probe (ABP) Me4BodipyFL-Ahx3Leu3VS (R&D Systems, I-190-050) was performed as described in^92^. Briefly, after native protein extraction in HR buffer and protein concentration measurement, as detailed in the native PAGE analysis sections of the methods, 80-100 μg were gently mixed with the ABP (1 μM) and incubated for 1 hr at 37°C. After labelling the samples were analyzed wither by SDS-PAGE or native PAGE and the in-gel fluorescence was acquired on the Azure Sapphire FL RGB system using the Cy3 filter set.

### Co-immunoprecipitation (co-IP)

Co-immunoprecipitation of POMP to identify its interactors under control and proteasome inhibition conditions was performed using a co-IP kit (Abcam, ab206996) and following the protocol detailed in the product’s technical manual. Briefly, DIV 28 cortical neurons transduced with GFP and HA-POMP-Myc-Flag AVV particles (150 MOI) on DIV7 were washed three times with DPBS+Ca^2+^/Mg^2+^ before lysis in ice using cold RIPA buffer supplemented with Halt protease and phosphatase inhibitor cocktail. The lysate was collected by scraping and transferred to a chilled microcentrifuge tube. After mixing on a rotary mixer for 30 minutes at 4°C, the samples were centrifuged at 10,000 g for 10 minutes at 4°C, and the supernatant was transferred to fresh chilled tubes. Protein concentration was determined using the Precision Red advanced protein assay and for every co-IP reaction 1 mg of clarified extract was mixed with 0.5 µg of either anti-POMP antibody (Cell signalling, 15141) or rabbit isotype control (Cell signalling, 3900) in a total volume of 500 µl with RIPA buffer supplemented with protease and phosphatase inhibitors. The reactions were incubated overnight at 4°C on a rotary mixer. The next day protein A/G Sepharose beads (25 μL per reaction) were washed twice with 1 mL wash buffer by centrifugation at 2000 × g for 2 minutes, with supernatant removal between washes. The beads were resuspended as a 50% slurry in wash buffer. Following antibody binding, 25 μl of bead slurry was added to each tube and incubated for 2 hr at 4°C with gentle mixing. Beads were collected by centrifugation at 2000 × g for 2 minutes at 4°C and washed three times with 1 ml of the wash buffer supplied with the kits, and once with DPBS+Ca^2+^/Mg^2+^. After the final wash, excess buffer was removed without allowing the beads to dry completely and elution was performed by incubating the slurries with 60 μL of 2x SDS sample buffer (0.125 M Tris, 4% (w/v) SDS, 20% (v/v) glycerol) at 95°C for 5 min. After elution the reactions were centrifuged at 2000 g and eluates were recovered and supplemented with TCEP (10 mM, Sigma-Aldrich, C4706-2G), Chloroacetamide (10 mM, Sigma-Aldrich, 22790-250G-F) and molecular biology grade water up to 100 μL. Co-immunoprecipitation to verify the ability of POMP to oligomerize in response to proteasome inhibition and probe the role of Cys36 was done on HEK293 transiently expressing the constructs POMP-Myc-Flag together with NLS-HA-POMP-Scarlet WT and the C36A mutant. Three days after transfection the cells were treated for five hours with 10 µM MG132, washed three times in DPBS+Ca^2+^/Mg^2+^ and lysed in RIPA buffer (Invitrogen, 89900). The extracts were clarified by centrifugation (104 g for 5 min) and the protein concentration was measured using the Precision Red advanced protein assay. The anti-Flag magnetic agarose slurry (Invitrogen, A36797) was equilibrated by three washing steps (500 µl) in Co-IP dilution buffer (10 mM Tris/Cl pH 7.5, 150 mM NaCl, 0.5 mM EDTA). Each reaction contained 100 µg of clarified protein extract, 25 µL of bead slurry, and Co-IP dilution buffer to a final volume of 500 µL. The reactions were incubated overnight at 4°C on a rotary mixer. The next day the beads were recovered on a magnetic tube rack, the supernatant was discarded and the beads were washed three times with 500 µl of Co-IP wash buffer (10 mM Tris/Cl pH 7.5, 150 mM NaCl, 0.05 % NP-40, 0.5 mM EDTA). Elution was performed as described for the POMP co-IP and the samples were then analysed by Western blot as described in the corresponding Methods’ section.

### MS sample preparation for proteomics

Lysates of HEK293 or yeast cells as well as of the subcellular fractions were prepared in SDS-PAGE loading buffer for bottom-up proteomic analyses as previously reported^95^. In brief, all samples were mixed with 2x lysis buffer (10% SDS, 100 mM TRIS, pH 7.55 adjusted with H_3_PO_4_) in a 1:1 ratio and reduced using dithiothreitol in a final concentration of 20 mM for 10 min at RT. For cysteine-residue alkylation, the samples were incubated with iodoacetamide in a final concentration of 50 mM for 30 min at RT in the dark. Then, the lysates were acidified using H_3_PO_4_ in a final concentration of ∼1.2%. Binding/ wash buffer (90% methanol, 50 mM TRIS, pH 7.1 with H_3_PO_4_) was added in a 1:7 lysate-to-buffer ratio. The protein suspension was loaded onto the filter of an S-trap (size = micro; ProtiFi) by centrifugation for 20 s at 4,000 x g in 150 µl-steps. Trapped proteins were washed with 150 µl binding/ wash buffer four times. MS-grade trypsin (1 µg; Promega) was added in 60 µl digestion buffer (40 mM ammonium bicarbonate) and digestion was carried out overnight at RT in a humidified chamber. For peptide elution, the filter was washed in three consecutive steps by centrifugation at 4,000 x g for 40 s with 40 µl digestion buffer and two 40 µl washes 0.2% formic acid in MS grade water. Peptides were desalted using C18 stage-tips^96^, dried by vacuum centrifugation and stored at −20°C until LC-MS analyses. Immunoprecipitated proteins in SDS-PAGE loading buffer were prepared using an SP3 approach^97,98^. In brief, 0.5 µl magnetic, carboxylate-modified beads (1:1 mix of Sera-Mag SpeedBeads (GE Healthcare, 45152105050250 & 65152105050250; pre-washed in MS-grade water three times) were added to the samples. Protein binding was facilitated by addition of 100% MS-grade ethanol to a final concentration of 50% ethanol and shaking for 5 min at RT at 1000 rpm. Additional incubation was carried out for 15 min at RT without agitation. Beads were then washed three times with 100 µl 80% ethanol using a magnetic rack. Proteins were digested in 20 µl digest buffer (50 mM triethylammoniumbicarbonate) with ∼0.1 µg MS-grade trypsin (Promega) in combination with 0.1 µg LysC (Wako) at 37°C shaking at 1000 rpm overnight. Peptides were desalted using C18 zip tips (cat. no. ZTC18S960, Millipore), dried by vacuum centrifugation and stored at −20°C until LC-MS analysis.

### LC-MS analysis

Dried peptides were reconstituted in 5% acetonitrile (ACN), 95% water and 0.1% formic acid (FA). For proteomic profiling in yeast and human, peptides were analyzed using a nano-UHPLC (nanoElute, Bruker Daltonics) coupled to a trapped ion mobility quadrupole time of flight mass spectrometer (timsTOF Pro 2, Bruker Daltonics). Peptides were loaded directly onto the analytical column (25 cm × 75 µm column with 1.9 μm C18-beads (PepSep); at max. 950 bar) maintained at 50°C and connected to a 20 µm ZDV sprayer/ emitter (Bruker Daltonics). The mobile phases used were 0.1% FA in MS-grade water (buffer A) and 0.1% FA in MS-grade ACN (buffer B). In 90 min gradients, peptides were separated in a non-linear gradient ramping from 2% to 3% B in 1 min, to 17% B in 55 min, to 25% B in 21 min and 38% B in 13 min with a constant flow rate of 400 nl/min. For data acquisition in DIA-mode, the ‘long-gradient’ DIA-PASEF method with a 1.1s cycle time was employed, as previously published^99^. For the subcellular fractionation and immunoprecipitation experiments, peptides were analyzed using a nano-UHPLC (U3000, Dionex) coupled to an orbitrap Fusion Lumos tribrid mass spectrometer (ThermoFisher) via a NanoFlex Source. Peptides were loaded onto a PepMap 100 C18 trap column (75 μm id × 2 cm length, 3 μm particle size; Thermo Fisher) at 6 μl/min for 6 min with 2% ACN and 0.05% trifluoroacetic acid followed by separation on an C18 analytical column (75 μm id × 50 cm length, 1.7 μm particle size; CoAnn Technologies) maintained at 55°C. Peptides were separated by a 120 min non-linear gradient matching the published MS method^100^ using mobile phase A (0.1% FA in MS-grade water) and B (80% MS-grade ACN, 20% MS-grade water, 0.1% FA). The samples were analyzed in positive polarity and data independent acquisition (DIA) mode. The DIA method defined MS1 scans followed by 40 DIA scans with optimized segment widths^100^. In brief, the method had the following settings. Full scan: orbitrap resolution = 120k, AGC target = 125%, mass range = 350-1650 m/z and maximum injection time = 100 ms. DIA scan: activation type: HCD, HCD collision energy = 27%, orbitrap resolution = 30k, AGC target = 2000%, maximum injection time = dynamic.

### MS-data post-processing and statistical analyses

#### Proteomic profiling in yeast and human

To process DDA-data of the proteomic profiling experiments in yeast and human, the FragPipe computational platform (version 18; https://fragpipe.nesvilab.org/) with MSFragger (version 3.5) and Philosopher (version 4.4.0) was used to identify peptide sequences from tandem mass spectra for library generation via easypqp. Protein sequence databases of *Saccharomyces cerevisiae* (UP000002311) and *Homo sapiens* (UP000005640) were downloaded from the UniProtKB database (RRID:SCR_002380). Decoy and common contaminant protein sequences were appended to the original databases in the software. Both precursor and fragment mass tolerances were set to 20 ppm. Enzyme specificity was set to ‘stricttrypsin’ and up to 2 missed cleavage sites were allowed. Peptide lengths from 7 to 50 and peptide masses from 500 to 5000 Da were allowed. For variable modifications, oxidation of methionine residues and acetylation of protein N-termini were included. Carbamidomethylation of cysteine residues was selected as a fixed modification. The maximum number of variable modifications per peptide was limited to 3.To process DIA-data of the respective samples, the computational platform dia-nn (RRID:SCR_022865, version 1.8) together with the spectral libraries (human or yeast, respectively) generated in the FragPipe was used. Mass accuracy tolerance was set to 10 ppm for all spectra. The default protein inference setting was disabled to base protein grouping on the generated spectral library with the FragPipe-settings. Quantification mode was selected as “robust LC (high precision)”. The match-between-run option was disabled. If not stated otherwise, dia-nn settings were set to default values. For the analysis of the proteomic profiles, the dia-nn R package (github.com/vdemichev/DiaNN) was employed. Protein quantification was performed using MaxLFQ-based LFQ-algorithm after filtering for precursor and global protein q-values < 1%. Differential protein expression, comparing knock-out and wildtype or siRNA and scrambled siRNA control, respectively, was assessed in an unpaired, two-sided t-test for each protein. Proteins had to be quantified in all biological replicates of both conditions. To adjust for multiple testing, Benjamini-Hochberg correction was applied with an FDR cut-off < 1%. Gene ontology (GO) overrepresentation analysis of the significantly regulated proteins was performed using the Gene List Analysis tool of the Panther Classification System (RRID:SCR_015893; pantherdb.org). GO term overrepresentation was tested by Fisher’s exact test with Benjamini-Hochberg FDR correction (< 5%).

### Proteomic profiling of subcellular fractions from rat primary rat cortical neurons

DIA raw files of the subcellular fractions were processed with dia-nn (version 1.8.2 beta 27) using a library-free approach. The predicted library was generated using the in silico FASTA digest (Trypsin/P) option with the UniProtKB database for *Rattus norvegicus* (UP000002494). Peptide length was set to 7-35 amino acids, missed cleavages to 2, and charge range to 1-5. Methionine oxidation was set as variable modifications and Cysteine carbamidomethylation as fixed modification. The maximum number of variable modifications per peptide was two. Protein inference was performed using genes with the heuristic protein inference option enabled. No normalization was applied (“--no-norm”). If not stated otherwise, dia-nn settings were set to default values. For the analysis of the subcellular fractions, the dia-nn output was imported in R via the dia-nn package and protein quantification was performed using MaxLFQ-based LFQ-algorithm^101^ filtering for proteotypic peptides and a precursor and protein level q-values < 1%. Log2-transformed protein intensities were normalized by median-centering. To assess protein enrichment of each fraction, a one-sided, one-sample t-test was used comparing protein abundance in the respective fraction to the overall, average expression (group mean). To adjust for multiple testing, Benjamini-Hochberg correction was applied with an FDR cut-off < 5%. GO overrepresentation analysis of the enriched proteins in each fraction was performed using the ShinyGO tool (v0.82, https://bioinformatics.sdstate.edu/go/; FDR < 5%) with a custom background list of all proteins identified in a whole cell lysate sample, applying default settings. Nodes and edges of the overrepresented GO-term networks were exported from ShinyGO and imported in CytoScape (RRID:SCR_003032) for visualization.

### POMP interactome

LC-MS/MS samples immunoprecipitation experiments were analyzed with Spectronaut version 18 (Biognosys) followed by statistical analysis using the MSstats R package. Targeted data extraction of DIA-MS acquisitions was performed in Spectronaut 18^102^ in DirectDIA mode with default settings using the UniProt reference proteome database for *Rattus norvegicus* (retrieved in December 2023), a database consisting of common mass spectrometry contaminants and the virally introduced POMP construct including tags. Pooled samples from a small subset of the experimental samples were acquired in DDA mode and added as library extension runs to the Spectronaut analysis. The proteotypicity filter “only protein group specific” was applied. Extracted features were exported from Spectronaut for statistical analysis with MSstats 4^103,104^ using default settings. Briefly, for each protein, features were log-transformed and fitted to a mixed effect linear regression model for each sample. For significance testing, contaminants were filtered out, all values below 5 were set to NA and we required a minimum of 4 features (combination of precursor and fragment ion) and 8 measurements per protein per condition. The model estimated fold change and statistical significance for all conditions. The Benjamini–Hochberg method was used to account for multiple testing and p-value adjustment was performed on all proteins that met the fold-change cutoff. Significantly different proteins were determined using a threshold fold-change >1.5 and adjusted p-value < 0.05. To compare POMP-enriched proteins between MG132 and DMSO treatment conditions, we tested for interaction effects in the linear model used by MSstats. Specifically we computed the contrast: anti-POMP+MG132 vs. anti-POMP+DMSO = (“anti-POMP MG132” - “Isotype MG132”) - (“anti-POMP DMSO” - “Isotype DMSO”), which captures differential enrichment of POMP-associated proteins between the two conditions. All further analysis was conducted using quantitative values obtained from MSstats 4 in the R environment (R Studio 2024 running R 4.2). Gene Ontology analysis was performed using the enrichR package^105^.

### Cytoplasmic nuclear fractionation

Isolation of cytoplasmic and nuclear fractions was performed following a modified version of the REAP method^106^. Briefly, HEK293 cells grown in 10 cm dishes up to ∼70-80% confluence were recovered by treatment with 3 ml/dish of TrypLE Express at 37°C for 5 min. Trypsinization was stopped with 7 ml/dish of cold culture medium, the cell suspension was moved to a 15 ml conical tube and centrifuged for 5 min at 500 g. After two washes with ice-cold DPBS+Ca^2+^/Mg^2+^ and recovery by centrifugation for 5 min at 500 g at 4°C, the cell pellet was resuspended in 450 µl of cytoplasm fractionation buffer (1% (v/v) NP-40, 1x Halt protease and phosphatase inhibitor cocktail in DPBS+Ca^2+^/Mg^2+^) and triturated by repeated gentle pipetting (5 strokes) with a P1000 pipette. To collect a whole cell extract (WCE) sample, 50 µl of the cell extract were taken and frozen down for later analyses. The cell suspension in cytoplasm fractionation buffer was spun down for 10 min at 3000 g at 4°C. To avoid nuclear contamination, as a cytoplasmic fraction the top 230 µl of the supernatant were collected, while the remaining ∼150-200 µl of the supernatant were discarded. The nuclear pellet was washed twice by gentle resuspension with ice-cold DPBS+Ca^2+^/Mg^2+^ and centrifugation for 5 min at 3000 g at 4°C. To prevent nuclear disruption and leakage of nuclear components, the pellet was carefully resuspended by gentle flushing. After the final wash, excess buffer was removed and the nuclear pellets were lysed in 60 µl of nuclear fractionation buffer (0.1% (v/v) SDS, 0.5% (v/v) Triton-X-100, 1% (v/v) NP-40, 1x Halt protease and phosphatase inhibitor cocktail in DPBS+Ca^2+^/Mg^2+^). To match the buffer composition across the different fractions, the WCE and cytoplasmic fractions were supplemented with 0.1% (v/v) SDS and 0.5% Triton-X-100. The WCE and the nuclear fractions were briefly sonicated to ensure efficient protein extraction, the extracts were clarified by centrifugation (10 min, 104 g at 4°C) and the protein concentration of the different fractions was determined using the Precision Red advanced protein assay. Equal amounts of proteins from the different fractions were used for downstream analysis.

### Subcellular fractionation

Two 10 cm dishes seeded with 4×10^6^ primary rat cortical neurons (28 DIV) were pooled for each subcellular fractionation reaction. All centrifugation steps were performed at 4°C and samples were kept on ice throughout the procedure. Neurons were washed twice in DPBS+Ca^2+^/Mg^2+^, transferred from a 10 cm dish into 250 μL of fractionation buffer (20 mM HEPES pH 7.4, 10 mM KCl, 5 mM MgCl2, 5 mM MgAc, 1mM DTT and Halt protease and phosphatase inhibitor cocktail) by scraping, pooled and incubated on ice for 15 minutes. The cell suspension was passed through a 27G needle ten times using a 1 ml syringe and the lysate was incubated on ice for 20 minutes. The sample was centrifuged at 700 g for 6 minutes and the resulting pellet corresponded to fraction I. The supernatant (corresponding to fractions II-IV) was carefully transferred into a fresh tube and kept on ice for further processing. The pellet (corresponding to fraction I) was washed with 500 μL of fractionation buffer, dispersed with a pipette, and gently passed through a 23G needle five times. The sample was centrifuged again at 700 g for 12 minutes. The supernatant was discarded and the pellet (corresponding to fraction I) was retained. The supernatant from the first centrifugation step (corresponding to fractions II-IV) was centrifuged at 104 g for 5 minutes. The resulting pellet was washed three times with 500 μL of fractionation buffer and corresponded to fraction II. The supernatant (corresponding to fractions III-IV) was transferred to a fresh tube and ultracentrifuged at 105 g for one hour. The supernatant (corresponding to fraction IV) was collected and the resulting pellet was washed by adding 400 μL of fractionation buffer, resuspended by pipetting, and passed through a 23G needle. The sample was re-centrifuged for one hour at 105 g, and the membrane pellet (corresponding to fraction III) was recovered. All pellets collected throughout the procedure were resuspended in TBS with 0.1% SDS (v/v) and Halt protease and phosphatase inhibitor cocktail, and sonicated briefly on ice (3 seconds at power setting 2-continuous) to shear genomic DNA (fraction I) and homogenize the lysate (fractions I-III). To match the buffer composition across the different fractions, fraction IV was supplemented with 0.1% (v/v) SDS. The protein concentration of the different fractions was determined using the Precision Red advanced protein assay and equal amounts of proteins from the different fractions were used for downstream analysis.

### Sucrose density gradient centrifugation

Compressed gradients (10 mM HEPES-KOH, 2.5 mM MgCl2, 100 mM KCl, 10-55% Sucrose) were made by layering 55%-10% sucrose solution in Polypropylene ultracentrifuge tubes (14×89cm, Beckman Coulter, 331372). A 7 ml cushion of the 55% sucrose solution was pipetted into the tube and frozen. Five consecutive layers of decreasing sucrose concentrations (50%, 40%, 30% - 500 µl and 20%, 10% - 1ml) were added, allowing each layer to freeze before adding the next. Primary rat cortical neurons were plated at a density of 4×10^6 cells per 10 cm dish four weeks before treatment. Three to four 10 cm dishes were treated with DMSO or MG132 for 5h, washed on ice twice with 10 ml of ice-cold DPBS+Ca^2+^/Mg^2+^ and lysed by scraping in 200 µl of non-denaturing lysis buffer (Abcam, ab206996) supplemented with RNase Inhibitor (Promega, #N2611). The lysate was homogenized by passing it through a 27G needle ten times and centrifuged at 10,000 g for 10 min at 4°C to clear the lysate. The cleared lysates were transferred to a fresh tube and the RNA concentration was measured on a nanodrop. For both conditions amounts of sample corresponding to OD260 3-4 were brought up to the same volume with non-denaturing lysis buffer supplemented with RNase Inhibitor, ABP (Me4BodipyFL-Ahx3Leu3VS,1 µM), DTT (1 mM) and ATP (2 mM) were added, and samples were then incubated for 1 hr at 37°C before loading onto the gradient. For ultracentrifugation, gradients were thawed in the cold room a day before use. After layering the lysates and pre-cooling the centrifuge, gradients were spun at 36,000 rpm for 3 hours at 4°C in an SW41Ti rotor. After centrifugation, the gradients were displayed using a continuous flow system, passing through a UV spectrophotometer to measure absorbance at 254 nm, generating a polysome profile. Fractions were collected every four seconds (850 μl/min flow rate) using an automated fraction collector.

### Droplet digital RT-PCR

HEK293 cells treated with proteasome inhibitors (Carfilzomib, Epoxomicin, MG132) for five hours were washed twice in DPBS+Ca^2+^/Mg^2+^ and total RNA was extracted using the RNeasy mini kit (Qiagen, 74104) according to the manufacturer’s instructions. RT-ddPCR was performed using a one-step RT-ddPCR advanced kit for probes (Bio-Rad, 1864021), as described in the product’s technical bulletin. The primer-probe sets used were ordered from IDT and their sequences are hereby reported: ACTB_Probe-5HEX/TCATCCATG/ZEN/GTGAGCTGGCGG/3lABkFQ, ACTB_F- ACAGAGCCTCGCCTTTG, ACTB_R-CCTTGCACATGCCGGAG, POMP_Probe-56- FAM/AGCTGCGGA/ZEN/AGATGAATGCCAGA/3IABkFQ, POMP_F-TCGGAAACGGAAGTGAGC, POMP_R-AGGTCCACTTGCTGAAAGTTC. Briefly, reaction mixes (20 μl/reaction) were prepared as follows: 20 ng total RNA, 900 nM/250 nM of primers/probe and 1x ddPCR supermix for probes and 50- 300 ng of genomic DNA (gDNA). The reactions were loaded into DG8 cartridges (Bio-Rad, 1864008) with droplet generation oil for probes (Bio-Rad, 1863005), the cartridge was then fitted with the DG8 gaskets (Bio-Rad, 1863009) and run on the QX200 droplet generator (Bio-Rad, 10031907). The droplet-oil emulsions were transferred to a ddPCR-compatible 96-well plate that was sealed with the PX1 PCR Plate Sealer (Bio-Rad) and the PCR reaction was run on a C1000 touch thermal cycler following the PCR protocol detailed in the technical bulletin. After thermal cycling, data were acquired on the QX200 Droplet Reader (Bio-Rad) using the QuantaSoft Software (Bio-Rad). A water-blank reaction was included as negative control. For analysis, the concentration of POMP mRNA molecules across the 6 biological replicates of the same treatment were normalized to their corresponding ACTB mRNA concentration. For each treatment condition the POMP mRNA fold-change (FC) relative to the DMSO control was calculated.

### RNA isolation and sequencing

RNA was isolated from HEK293 cells 72h after transfection with siRNA oligos using the Quick-RNA 96 kit (Zymo Research, R1052) according to the kit’s specifications and quantified using a Qubit Fluorometer (Invitrogen, Q33216). Input RNA quality was assessed by running the RNA samples on an Agilent Bioanalyzer RNA 6000 Nano Chip. mRNA sequencing libraries were prepared starting from 300ng of total RNA using NEBNext poly(A) mRNA Magnetic isolation (NEB, E7490) together with Ultra II Directional RNA library Prep Kit for Ilumina (NEB, E7760). To achieve a longer RNA insert size, we modified the RNA fragmentation time and cDNA size selection based on the manufacturer’s recommendations. cDNA libraries were amplified and multiplexed using NEBNext® Multiplex Oligos for Illumina® (96 UD-Index Primer Pairs, E6440). The quantity of the DNA libraries was measured on Qubit fluorometer and quality assessed on an Agilent Bioanalyzer HSDNA 2100 Chip. Libraries were sequenced on an Illumina NextSeq2000 using P3 XLEAP-SBS reagents with 151 bp paired-end reads.

### RNA-Seq preprocessing

Raw Illumina paired-end RNA sequencing data were processed on a high-performance computing cluster using a custom bash pipeline. Demultiplexing of BCL files was performed using bcl2fastq2 (v2.20.0.422), guided by a user-provided sample sheet, to generate per-sample FASTQ files. Adapter trimming and quality filtering were applied using fastp (v0.24.0), which also removed polyX tails, trimmed low-quality bases from both ends, and discarded reads shorter than 51 bp or with low complexity or average quality below Q20. Cleaned reads were aligned to the human reference genome (hs1) using STAR (v2.7.11b), producing coordinate-sorted BAM files with splice junction information. Read quantification at the gene level was then performed using subread/featureCounts (v2.0.8) with exon-level counting and gene ID grouping, using the corresponding genome annotation. All steps were parallelized for efficient execution across multiple samples.

### Sample quality control and differential expression analysis

Post-alignment read counts were imported into R (v4.4.3) and analyzed using the DESeq2 package (v1.44.0). Raw exon-level counts were aggregated across technical replicates and siRNA conditions to create a merged count matrix. Principal component analysis (PCA) was performed on variance-stabilized transformed (VST) data to assess sample quality and replicate clustering. Differential expression (DE) analysis was conducted using DESeq2, applying a linear model design = ∼replica + condition that accounted for both biological replicate and sample group. Log2 fold changes were shrunk using the lfcShrink function to improve interpretability, and genes with s-values below 0.05 and absolute log2 fold change greater than 0.3 were considered significantly differentially expressed. The log2 fold change cut-off was chosen based on the fact that siRNA knockdown of PSMB1/7 and POMP led to a 50% reduction in proteasome activity, so changes with a magnitude of 30% were considered biologically meaningful. Results and normalized counts were exported for downstream analysis and visualization.

### Differential Isoform Usage Analysis

Transcript-level expression was quantified using Salmon (v1.10.3) in quasi-mapping mode with automatic library type detection (--libType A), using a RefSeq-derived transcriptome index. Cleaned paired-end FASTQ files were grouped by experimental condition and quantified independently. Transcript abundance estimates were imported into R using the tximport package (v1.32.0), and summarized to the gene level via a transcript-to-gene mapping table. Gene-level differential expression analysis was performed using DESeq2 (v1.44.0), with a design formula accounting for biological replicates ∼replica + condition. Pairwise comparisons between POMP, PSMB, and SCRM conditions were tested using Wald statistics and further refined using lfcShrink with an lfcThreshold of 0.3 and svalue < 0.05 as the significance cutoff. Log2 fold changes were computed using the ashr shrinkage estimator for combined contrasts. For isoform-level analysis, transcript counts were imported from Salmon with txOut = TRUE and combined with the same transcript-to-gene annotation. Prior to analysis with DRIMSeq^107^ (v1.32.0), a pseudocount of +1 was added to all counts to stabilize downstream modeling. The data were filtered using dmFilter() with the following criteria: a gene was retained if it had expression in at least 15 samples (min_samps_gene_expr = 15), each transcript (feature) was required to be expressed in at least 6 samples (min_samps_feature_expr = 6), and both gene- and transcript-level expression had to exceed 10 counts (min_gene_expr = 10, min_feature_expr = 10). Additionally, transcripts contributing less than 10% to total gene expression were excluded (min_feature_prop = 0.1). A full model matrix including both replicate and condition effects was fitted (∼replica + condition), and differential usage was assessed using dmTest() for each contrast. Gene-level adjusted p-values were reported, and transcript usage proportions were extracted using DRIMSeq::proportions() for visualization and interpretation.

### Gene ontology (GO), pathway and feature enrichment analysis

Gene ontology (GO), pathway and feature enrichment analyses were performed using the ShinyGO^108^ tool (v0.82, https://bioinformatics.sdstate.edu/go/). The analysis included enrichment for Biological Process (BP), Molecular Function (MF), Cellular Component (CC), pathway databases such as KEGG and feature enrichment analysis (i.e. transcript length, 5′ UTR and 3′ UTR length, CDS length, genome span, GC content, number of exons and transcripts per gene). A false discovery rate (FDR) of <0.05 was used as the threshold for significance. The background gene list used comprised all the genes sequenced in the experiment. REVIGO^109^ (http://revigo.irb.hr/) was used to remove redundancy from GO term lists by clustering semantically similar terms.

### Transcript Feature Extraction and Discriminative Modeling

To identify transcript features associated with condition-specific regulation, we computed a comprehensive set of mRNA characteristics from genomic annotations, nucleotide sequences, and RNA structural predictions. Features included gene length, transcript length, exon/intron structure (e.g., number of exons, maximum exon/intron length), GC/AT content and skew, and folding properties such as minimum free energy (MFE) per base. These were derived from BED12 gene models, CDS sequences, and secondary structure predictions using RNAFold and custom Perl scripts. For each gene, sequences were parsed from FASTA files, and per-region features (CDS, 5′UTR, 3′UTR) were calculated where available. CAI (Codon Adaptation Index) was computed from coding sequences using relative codon usage frequencies. Differential expression results from DESeq2 (log₂ fold change and condition label) were integrated to form the final feature matrix. Transcript features were log-transformed (e.g., log10(len + 1)) and standardized using z-scores. To explore overall structure, principal component analysis (PCA) was performed on the scaled data, and variance contributions were extracted from component loadings. To identify features that discriminate conditions, linear discriminant analysis (LDA) was applied with transcript condition as the target variable. LDA loadings (absolute value of coefficients) were used to rank feature importance. In addition, regularized regression models (Lasso via glmnet), gradient-boosted trees (xgboost), and multivariate adaptive regression splines (MARS) were trained for both regression (predicting log₂ fold change) and classification (predicting transcript condition). Feature importance was assessed using model-specific metrics (e.g., coefficient magnitude for GLM, gain for XGBoost), and aggregated across all models. The final feature ranking was obtained by averaging scaled importance values across PCA, LDA, GLM, XGBoost, and MARS models. Visualizations and ranking tables were generated to highlight features with the highest discriminatory power.

### Generation of mESC-derived cortical neuron cultures

mESC-derived cortical neuron cultures established from DIV 17 cortical organoids generated following previously described unguided cerebral organoid protocols^110^ with modifications in timing to match the intrinsic developmental tempo of the cells. Briefly, E14Tg2A mESCs were maintained as feeder-free cultures on serum/LIF-medium, as detailed in the corresponding section of the Methods. On day 0, mESCs were dissociated to single cells with TrypLE Express (Invitrogen, 12605028), the reaction was stopped with mESC culture medium and the cells were harvested (500 g, 5 min) before dissociation in commercial EB Formation medium (StemCell Technologies, STEMdiff Cerebral Organoid kit, 8570) supplemented with 50 μM ROCK inhibitor Y27632 (SantaCruz Biotechnology, sc-281642A). For embryoid body (EB) generation, 4×103 cells/well were seeded into ultra-low attachment U-bottom 96-well plates (Corning, 7007). On day 2, the culture media was replaced with Induction medium (StemCell Technologies, STEMdiff Cerebral Organoid kit, 8570). On day 5, EBs were moved to Expansion medium (StemCell Technologies, STEMdiff Cerebral Organoid kit, 8570) supplemented with dissolved Matrigel (Corning #354234, DF 1:6) and transferred to standard 6-well plates. On day 6, the culture media was replaced with Maturation medium (StemCell Technologies, STEMdiff Cerebral Organoid kit, 8570). On day 7, Matrigel was removed from the organoids by repeated gentle washes in Maturation medium, and the organoids were then moved to 5 cm dishes on an orbital shaker (57 RPM, 25 cm orbit; 0.45 g). On day 14 the organoids were harvested, incubated with Accumax (Invitrogen, 00-4666-56) supplemented with 400 µg/ml DNAse I (Invitrogen, EN0521) and were subject to four 5 min incubation steps at 37 ° C. After the first 5 min the organoids were resuspended by flicking the tube, after 10 min the organoids displayed a fluffy appearance and were pipetted up and down once, then broken into cell clumps. After 15 min the cell clumps triturated by pipetting up and down 3–5 times, and then 10 more times after an additional 5 min incubation. Dissociation was stopped by addition of an equal volume of Maturation medium. The cell suspension was passed through a 70 µm nylon cell strainer (Corning, 352350). A small aliquot was taken for a live cell count and the remaining cell suspension was spun down at 300 x g for 5 min. The cell pellet was resuspended in Maturation medium and the cells were plated on PDL-coated cell cultureware, 105 cells/MatTek and 5×105 cells/well on 24-well plates. On day 18, the dissociated cultures were switched to Maturation medium supplemented with BDNF (Abcam, ab9794, 20 ng/ml) and mNT3 (Prospec, CYT-688, 20 ng/ml) to enhance synaptic maturation. The cultures were maintained with weekly feedings and phenotypic changes were assayed at 28-30 days in culture.

### Calcium imaging

One week after dissociation, mESC-derived neuronal cultures were transduced with AAV1-pENN.AAV.CamKII.GCaMP6f.WPRE.SV40 (Addgene, 100834-AAV1) at a multiplicity of infection (MOI) of 100, calculated as viral genome copies per cell (GC/cell). Calcium imaging was performed on 28–30-day-old cultures using a Zeiss LSM 880 confocal laser scanning microscope equipped with a Plan-Apochromat 40×/1.3 Oil lens, at an acquisition rate of 3–4 fps. Cells were imaged in the same NGM used for culture, on an incubated stage maintained at 37°C with 5% CO₂.

### Pharmacological treatment

The compounds and cytokines used for cell treatment along with vendor and catalog number, the solvent used for reconstitution, the concentration used and treatment duration (in those cases where only one was used) are hereby listed: Carfilzomib (Abcam, ab216469, DMSO, 2 µM), Epoxomicin (Millipore, 324800, DMSO, 2 µM), MG132 (Invitrogen, J63250.MCR, DMSO, 10 µM), Bortezomib (Invitrogen, J60378, DMSO, 1 µM), HSF1B (Axon Medchem, 2101, DMSO, 70 µM, 7 hr), KNK-437 (Sigma-Aldrich, SML0964, DMSO, 100 µM, 5hr in combination with proteasome inhibition), DTT (Millipore, 111474, 1 mM, H2O, 5hr in combination with proteasome inhibition), H2O2 (AlfaAesar, L13235, H2O, 1 mM, 1hr) TNFα (Peprotech, 300-01A-50UG, PBS+0.1%BSA, 0.2 µg/ml, 4 days), IFNα (Abcam, ab48750, PBS+0.1%BSA, 0.2 µg/ml, 4 days), IL1β (CST, 54059, PBS+0.1%BSA, 0.2 µg/ml, 4 days), GM-CSF (CST, 87015, PBS+0.1%BSA, 0.2 µg/ml, 4 days), chloroquine (Fisher Scientific, C230125G, H2O, 100 µM), (Tocris, 0130, DMSO, 10 µM), CNQX (Tocris, 1045, H2O, 20 µM), APV (Tocris, 0106, H2O, 50 µM).

### Splicing assay

HEK293 cells (50-60% confluent) were transfected with pcDNA3.1-NoTS-HA-mTagBFP2 (1 µg/ml) and pcDNA3.1-NoTS-HA-POMP-mTagBFP2 (0.5 µg/ml) combined with either the RG6 or the pHBS1389 IBB-GFP-mCherry3E splicing reporter (0.5 µg/ml). Transfection was performed as detailed in the corresponding Methods’ section. Three days after transfection the cells were fixed with PFA fixative solution (4% PFA (v/v), 4% sucrose (v/v), 1 mM MgCl2, 0.1 mM CaCl2 in PBS pH 7.4) for 15 min at room temperature and then imaged on a Zeiss LSM 880 confocal laser scanning microscope with a Plan-Apochromat 40×/1.3 Oil lens. Image analysis was performed using a custom custom-made Fiji macro as described in the ‘Image analysis’ section of the Methods.

### Propidium Iodide (PI) uptake cell death assay

HEK293 cells (50-60% confluent) grown on MatTek dishes were transfected with siRNAs against PSMB1, PSMB7, POMP and SCRM and cultured for 72h cells before PI staining. The cells were washed twice with DPBS+Ca^2+^/Mg^2+^, stained with 1 µg/mL PI (Invitrogen, P1304MP) and 5 µM DRAQ5 (CST, 4084) for 10 minutes in the dark, and then fixed with PFA fixative solution (4% PFA (v/v), 4% sucrose (v/v), 1 mM MgCl2, 0.1 mM CaCl2 in PBS pH 7.4) for 10 minutes at room temperature. Coverslips were mounted with coverslips using Aqua-Poly/Mount (Polysciences, 18606-20) and imaged on a Zeiss LSM 880 confocal laser scanning microscope with a Plan-Apochromat 20x/0.8 lens. PI-positive cells were quantified using custom-made Fiji macro described in the ‘Image analysis’ section of the Methods.

### CellROX Green staining

Cells were treated with proteasome inhibitors and 1 mM DTT (positive control), as detailed in the ‘Pharmacological treatment’ section of the Methods. In the last 30 min of treatment, CellROX Green reagent (Invitrogen, C10444) was added to the culture medium at a final concentration of 5 μM. The cells were then incubated at 37°C for 30 minutes to allow for reagent uptake. After incubation, the medium was removed, the cells were washed three times with DPBS+Ca^2+^/Mg^2+^ and fixed with PFA fixative solution (4% PFA (v/v), 4% sucrose (v/v), 1 mM MgCl2, 0.1 mM CaCl2 in PBS pH 7.4) for 15 min at room temperature. After fixation the cells were processed for immunofluorescence analysis as described in the corresponding Methods’ section. Images were acquired on a Zeiss LSM 880 confocal laser scanning microscope using a Plan-Apochromat 40×/1.3 oil lens and fluorescence intensity was quantified as described in the ‘Image analysis’ section of the Methods.

### *In situ* hybridization

HEK293 72 hrs after transfection were fixed at room temperature for 20 minutes using a solution containing 4% paraformaldehyde, 5.4% glucose, and 0.01 M sodium metaperiodate in lysine-phosphate buffer. In situ hybridization was carried out using the ViewRNA ISH Cell Assay Kit (Invitrogen), following the manufacturer’s protocol with slight modifications. In brief, cells were permeabilized at room temperature with detergent solution for 5 minutes, then treated with pepsin (0.01 mg/mL in 10 mM HCl) for 45 seconds. After rinsing with PBS, target-specific probes (Invitrogen, *CXCL10* VA6-13729, *MYC* VA6-10461) were diluted 1:100 in pre-warmed hybridization buffer and applied to the cells. Hybridization was performed at 40 °C for 3 hours, followed by washes and storage overnight at 4 °C in storage buffer. The next day, cells were incubated at 40 °C with three amplification steps: first with PreAmplifier mix (1:25) for 30 minutes, then with Amplifier mix (1:25) for 30 minutes, and finally with Label probe mix (1:25) for another 30 minutes—each diluted in their respective pre-warmed working buffers. After several washes, immunostaining was performed as detailed in the corresponding section of the methods, ACTB was used to stain all cells and identify untransfected cells.

### Metabolic labelling of RNA with 5-ethyl uridine (5-EU)

Metabolic labelling of RNA was performed using the Click-iT RNA Alexa Fluor (Invitrogen, C10330) according to the manufacturer’s instructions. Briefly, HEK293 were transfected with either pcDNA3.1-NoTS-HA-Neon (1 µg/ml) or pcDNA3.1-NoTS-HA-POMP-Neon (500 ng/µl) as described in the ‘Plasmid transfection’ section of the Methods. Three days after transfection the cells were fixed PFA fixative solution (4% PFA (v/v), 4% sucrose (v/v), 1 mM MgCl2, 0.1 mM CaCl2 in PBS pH 7.4) for 15 minutes at room temperature. After fixation, the cells were washed once with DPBS+Ca²⁺/Mg²⁺ before incubation with permeabilization solution (0.5% Triton-X-100 (v/v) 4% goat serum (v/v) in PBS pH 7.4) for 15 minutes at room temperature. A 1X working solution of Click-iT reaction buffer was freshly prepared and used on the same day. The Click-iT reaction cocktail was prepared according to the manufacturer’s instructions, ensuring that components were added in the specified order. Following permeabilization, cells were washed once with DPBS+Ca²⁺/Mg²⁺, and 500 µl of Click-iT reaction cocktail was added to each sample. The reaction was allowed to proceed for 30 minutes at room temperature, protected from light. After incubation, the reaction cocktail was removed, and the cells were washed once with 1 ml of Click-iT reaction rinse buffer. The samples were subsequently processed for immunofluorescence analysis, as described in the corresponding Methods section. When azide-PEG3-biotin (Sigma-Aldrich, 762024, 100 µM) was used for labeling instead of Alexa Fluor 594 azide, the secondary antibody solution included 2 µg/mL streptavidin Alexa Fluor 647 conjugate (Invitrogen, S32357). The cells were imaged on a Zeiss LSM 880 confocal laser scanning microscope using a Plan-Apochromat 63x/1.40 Oil lens. The fluorescence images were quantified as described in the ‘Image analysis’ section.

### Experimental animals

All animal procedures were conducted in accordance with institutional and national ethical guidelines. Experimental cohorts included male and female mice from the R6/1 transgenic line expressing mutant Huntingtin (mHtt; Jackson Laboratory, 006471) and the Nex-Cre;3xMetRS transgenic line expressing mutant Huntingtin (mHtt; doi: 10.1101/2024.11.28.625838). These animals are referred to as HTT were analysed aged 7 months. These were compared to male and female age-matched and one year old controls lacking the mutant transgene. For wild-type controls, referred to as WT, C57BL/6J (Jackson Laboratory, 000664) and GAD-Cre;1xMetRS^111^ transgenic mice were used. All animals were housed under a 12-hour light/dark cycle with *ad libitum* access to food and water. The animals were maintained with permission from the local government offices in Spain (Committee of Animal Experiments at UCM and Environmental Counseling of the Comunidad de Madrid, protocol number: PROEX 005.0/21).

### Mouse brain tissue dissection and lysis

Mice were euthanized using CO₂ and immediately decapitated. Whole brains were then extracted, hippocampi were dissected out and snap-frozen on dry ice. The dissected hippocampi were lysed in 200 µl of HR buffer (50 mM Tris, 5 mM MgCl2, 250 mM sucrose, 1 mM DTT and 2 mM ATP, pH=7.4) by first using a motorized homogenizer (VWR VOS14, 10x strokes, 1000 rpm) and by passing the homogenate through a 27G needle using a 1 ml syringe (10x strokes). After centrifugation (10^4^ g for 10 min at 4°C), the clarified extracts were incubated with the ABP and processed for downstream analyses as outlined in the ‘Fluorescence-Based Proteasome Activity Profiling’ section of the methods.

### Perfusion and tissues processing

Mice were anesthetized with isoflurane and transcardially perfused with 40 mL of 4% paraformaldehyde (PFA) in 1× PBS supplemented with 4% sucrose. Whole brains were carefully dissected and post-fixed in the same fixative for 1 hour at room temperature. Following fixation, brains were rinsed twice in 1× PBS and incubated overnight at 4 °C in 15% sucrose in PBS for cryoprotection. Brains were maintained in 15% sucrose at 4 °C until further processing. For cryostat sectioning into coronal sections, the olfactory bulb and hindbrain were removed with a scalpel, the remaining brain was plunge frozen in 2-methylbutane (Sigma-Aldrich, M32631) at ∼-40°C for 1-2 minutes, glued with O.C.T. compound (Science Services, SA62550-01) onto the cryostat stage and cryosectioned at a thickness of 20 μm. Sections were collected onto Superfrost Plus microscope slides (Invitrogen, 22037246), permeabilized for 4 hrs (0.5% Triton-X-100 (v/v) 4% goat/donkey serum (v/v) in PBS pH 7.4) at room temperature and stained overnight in permeabilization buffer with primary antibodies diluted as indicated in the ‘Antibodies’ section of the methods. The next day, the slides were washed three times in PBS, stained with secondary antibodies (DF=1:500) in permeabilization buffer for 2 hrs, washed four times in PBS and mounted with coverslips using Aqua-Poly/Mount (Polysciences, 18606-20).

### Microscopy

Images were acquired on either a Zeiss LSM 780 or 880 inverted confocal microscopes. Objectives used were: EC Plan-Neofluar 20x/0.50, Plan-Apochromat 40×/1.3 Oil lens and Plan-Apochromat 63x/1.40 Oil lens. Images were taken as z-stacks. Quantifications of data from all systems were compared to ensure reproducibility of the data.

### Image analysis

Image analysis was done using ImageJ2 v2.30/1.54. Depending on the type of analysis performed images were either projected or individual planes were used. All experiments analysed using a custom macro are listed below with a brief description of the analysis steps. For quantification of nucleolar POMP/p62/PSMA3/PSMG2 levels, Fibrillarin was used to generate a mask for the nucleolus with gaussian blur (σ = 1) and manual thresholding. The area and mean grey value of each mask was quantified in the POMP channel. For POMP and CellROX green quantifications in mESCs all channels were merged and thresholded after application of Gaussian blur (σ = 1). The thresholded image was converted to a mask, fill holes and watershed were applied before segmentation of ROIs. For each ROI the mean grey value was measured in the CellRox or POMP channel and reported as intensity fold change (FC) relative to control. For analysis of nascent RNA levels with 5-EU the Fibrillarin channel was processed by blurring (Gaussian blur, σ = 1) followed by automatic thresholding and used as a mask for the nucleolus. Area and raw integrated density were then quantified for the NoTS-HA-Neon and the NoTS-HA-POMP-Neon expression constructs in the nucleolus. For quantification of PI+ nuclei the PI and DRAQ5 channels were processed independently: images were blurred (Gaussian blur, σ = 1), converted to 8-bit, and automatically thresholded. Binary masks were created and watershed segmentation applied. The number of segmented particles per field of view was quantified for the PI channel (number of PI+) and this was reported as percent of total number of cells (number of DRAQ5+ cells). For analysis of the splicing reporter GFP (splicing reporter), BFP (construct tested) and RFP (splicing construct) channels were merged, Gaussian blurred (σ = 1), automatically thresholded and converted to binary masks. After watershed segmentation, ROIs were analyzed for mean gray value and the ratio of GFP over RFP was taken for each cell. Colocalization analyses were performed by measuring either the Mander’s overlap coefficient (MOC) or the Pearson’s correlation coefficient (PCC) using the ImageJ plug-in JACoP. For the experiments shown in Figure 5, to quantify the number of POMP nuclear puncta blind manual counting was performed using a custom plug-in, written by J. Schindelin, to measure the POMP intensity a mask was manually traced using a cell fill or DAPI and the POMP channel mean grey value was measured. Analysis of POMP nuclear relocalisation in WT and HTT cortical and hippocampal neurons was performed blind. Masks for the neuronal cytoplasm and nucleus were manually traced in the plane centered on the nucleus. Mean gray values were measured for both compartments, and their ratio (nucleus/cytoplasm) was calculated. To quantify ribosomal protein, POMP, and PSMA3 levels in transfected cells, analyses were performed blind. A mask was manually drawn around the cell fill channel marking transfected cells, and the mean grey value was measured in the relevant channel. Values from transfected cells were normalized to those of adjacent non-transfected cells to account for uneven immunostaining and potential laser instability during prolonged imaging. For analysis of the *in situ* hybridization puncta, masks were made for the transfected cell (using the cell fill channel) and the adjacent non-transfected cells, mRNA puncta (Find Maxima; threshold 340-500) were counted in both classes and the transfected count was normalised to adjacent non-transfected cells to account for uneven staining and potential laser instability during prolonged imaging. The POMP-NoTS data was expressed as a fold change relative to control (Neon-NoTS). Data were then normalized to the control sample and expressed as fold change relative to control. For immunoblot quantifications, the level of each protein was normalized to a loading control (ACTB, GAPDH, Lamin B1 or total protein) and expressed as fold change FC.

### Statistical analysis

Comparisons between two groups were conducted using unpaired two-tailed t-tests. For analysis of Western blots for which experimental replicates acquired on different membranes or on different days paired two-tailed t-tests on the loading control-normalized data were used. Ordinary one-way ANOVA tests were used when the experiment comprised more than two groups. For analysis of Western blots comparing more than two groups with experimental replicates acquired on different membranes or on different days RM one-way ANOVA tests were used. Post-hoc comparisons were conducted using Tukey’s test to assess pairwise differences among all experimental groups (in some cases the figures show only a subset of the comparisons), Šidák’s test to assess the pairwise differences among a subset of experimental groups, and Dunnett’s test to determine whether each treatment group differed significantly from the chosen control group. Statistical significance was set at p≤0.05. A minimum of three biological replicates was used for analysis in biochemical, proteomics and sequencing experiments. A minimum of fourteen cells and two biological replicates were used for confocal microscopy analyses. Statistical analyses and data visualization were performed using GraphPad Prism. Figures were assembled using Adobe Illustrator and schematics were in made in Illustrator with the help of Biorender.com.

## Acknowledgements

We thank the MPIBR Animal Facility, Imaging Facility and Proteomics Facility for their support and assistance with data acquisition. We are grateful to Ina Bartnik, Christina Thum, Nicole Fürst, Anja Staab for the preparation of primary cell cultures and members of the Schuman lab for helpful feedback and discussion. We thank Michaela Müller-McNicoll and Vladimir Despic for advice on splicing analyses, Walter Spevak for valuable input on apoptosis, and Noortje Kersten and Frederik Steiner for assistance in the early phases of the project. This work was funded by the Max Planck Society and DFG CRC 1080. S.L.G. was supported by a FEBS Long-Term Fellowship and a Hessen Horizon MSCA Fellowship.

## Author contributions

S.L.G. co-designed the project, performed all experiments, analysed data and co-wrote the manuscript. M.M. wrote Fiji analysis macros, analysed data and curated the figures. M.v.O. prepped and analysed IP-MS samples. K.D. prepped and analysed species comparison and subcellular fractionation proteomics samples. B.K. acquired microscopy images and performed image analysis. G.T. performed bioinformatic analysis. E.C. sequencing library prep. J.D.L. supervised proteomics. B.A.C. bred mice and harvested brain tissue for analysis. E.M.S. co-designed and supervised the project and co-wrote the manuscript. All authors contributed to manuscript editing.

---

**Figure S1.**
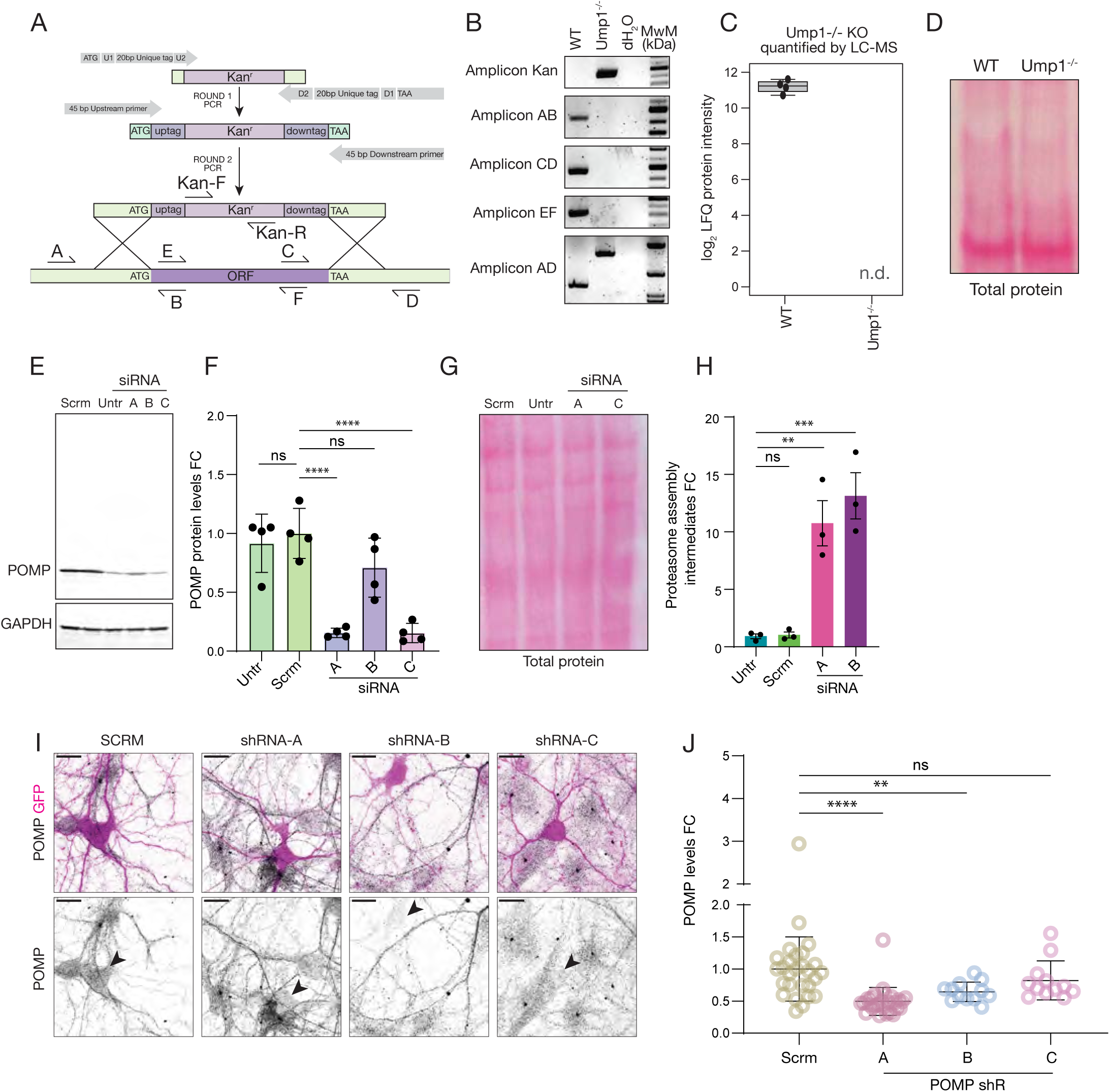
Validation of successful POMP depletion in yeast and mammalian cells. (A) Schematic representation of the yeast Ump1 locus, the genetic manipulation strategy used to knock-out Ump1 and primers used to genotype by PCR the WT and Ump1^-/-^ strains. (B) Genotyping PCR gels of WT and Ump1^-/-^ strains along with a water negative control sample and a molecular weight marker. Primer pairs AB, CD and EF, where at least one primer falls within the Ump1^-/-^ region ablated give a PCR product only the case of the WT. Primers to amplify the kanamycin selection cassette, used to disrupt the gene and select transformants, give a product only in the case of the Ump1^-/-^. Primer pairs AD yield a product in both strains but the size of the amplicon is higher in the Ump1^-/-^ strain due to addition of the kanamycin cassette used to disrupt the gene. (C) Label-free quantification (LFQ) of POMP protein abundance in WT and Ump1 Ump1^-/-^ yeast cells measured by liquid chromatography-mass spectrometry (LC-MS). Log₂-transformed LFQ intensity values are shown for individual biological replicates (dots). POMP protein was not detected (n.d.) in the KO samples, confirming efficient knockout at the protein level. n=4, the boxplot shows median, interquartile range, and 1.5×IQR whiskers. (D) Total protein stain corresponding to the blots and the Suc-LLVY-AMC fluorescence shown in Figure 1B. Loading was slightly higher in the WT sample. (E) Biochemical validation of the POMP antibody’s suitability for Western blot analysis and assessment of siRNA knock-down efficiency. HEK293 cells were transfected with siRNA constructs against POMP (A, B and C) or Scrm, and cultured for three days prior to Western blot analysis. This included the different siRNA samples and an untransfected control sample. siRNA A and C gave the strongest knock-down. The POMP antibody was highly specific and produced a single band at the expected molecular weight (∼16 kDa). (F) Analysis of experiments like the one shown in E. While siRNA B did not lead to a significant decrease in POMP protein levels, siRNA A and C yielded a strong and significant reduction in POMP. ns=p>0.05, ****p<0.0001, one-way ANOVA with post-hoc Dunnett’s test, n=4, mean±SD, FC=fold change. (G) Total protein stain corresponding to the blots and the Suc-LLVY-AMC fluorescence shown in Figure 1E. Loading was slightly higher in the POMP siRNA C sample. (H) Analysis of proteasome assembly intermediates abundance in native PAGE Western blots like the one shown in Figure 1E. Loss of POMP leads to a significant accumulation of proteasome assembly intermediates. **p<0.01, ***p<0.001, one-way ANOVA with post-hoc Dunnett’s test, n=3, mean±SD, FC=fold change. (I) Representative immunofluorescence images of primary rat hippocampal neurons transfected with constructs encoding GFP along with either one of three different shRNAs against POMP or a scramble control. GFP^+^ cells (upper panel, magenta) expressing shRNAs against POMP display reduced levels of POMP (grey) compared to neighbouring cells. This is not observed in the case of cells expressing the scramble control. In the POMP single channel images in the lower panel transfected cells are indicated by the arrowheads.Scale bars = 20 µm. (J) Analysis of images like the ones shown in I. The constructs expressing POMP shRNA A and B, but not C, leads to a significant reduction in POMP protein levels assayed by immunofluorescence. **p<0.01, ****p<0.0001, one-way ANOVA and post-hoc Dunnett’s test, n= 27 (Scr and shR A), 12 (shR B and C), mean±SD, FC=fold change.

**Figure S2.**
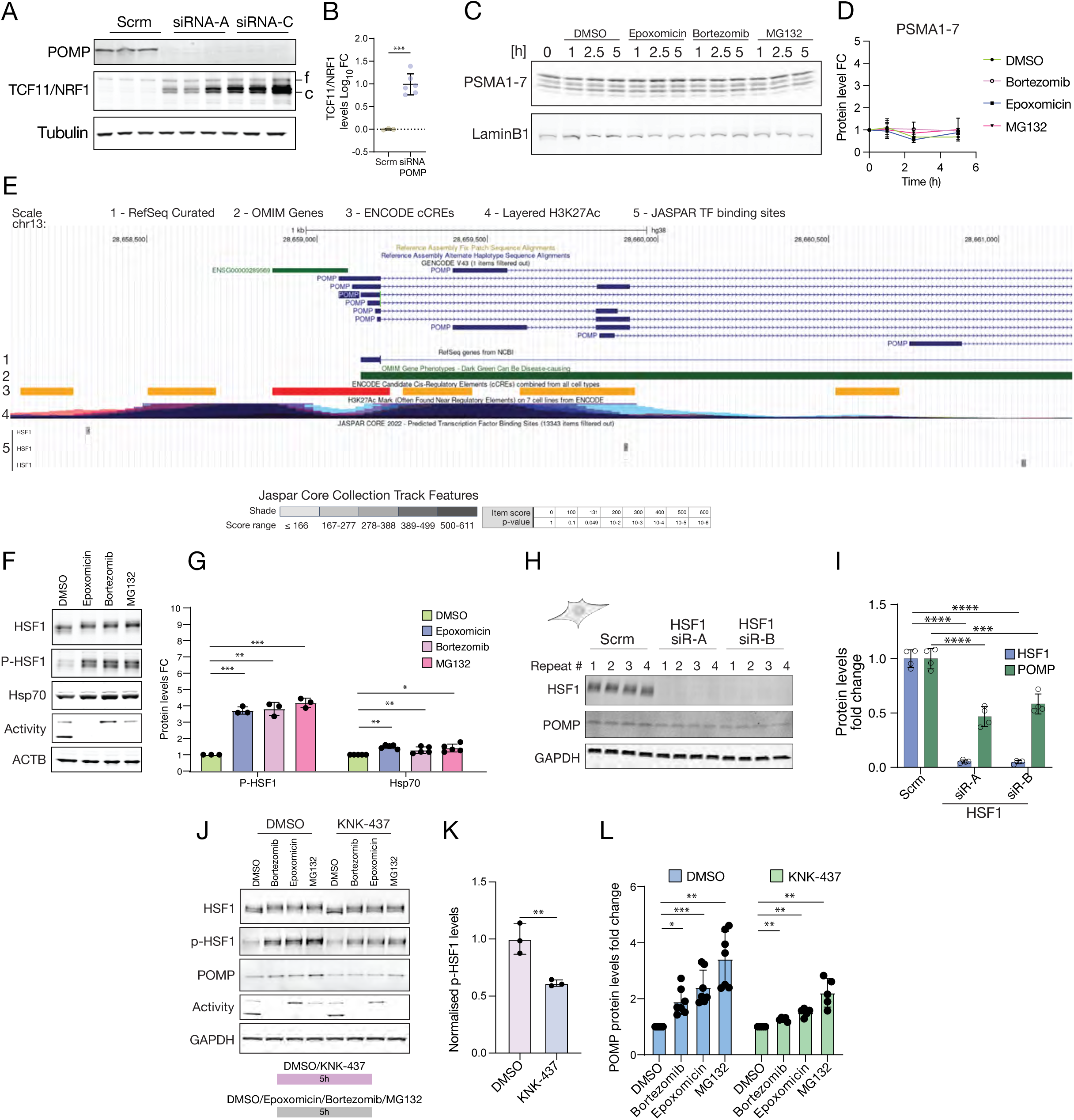
Regulation of POMP expression at the early stages of proteasome inhibition. (A) Western blot analysis of POMP and TCF11/NRF1 protein levels following siRNA-mediated knockdown of POMP using two independent siRNAs (A and C) in HEK293 cells. Scrambled siRNA (Scr) served as control. Tubulin was used as a loading control.The full-length (f) and cleaved (c) forms of TCF11/NRF1 are indicated next to the blot. (B) Analysis of the experiment shown in A. POMP knockdown leads to a significant increase in the levels of cleaved-TCF11/NRF1. ***p<0.001, unpaired two-tailed t-test, n= 3 (Scrm) and 6 (siRNA POMP), mean±SD, FC = fold change. (C) Representative Western blot time course from HEK293 cells treated with one each one of three different proteasome inhibitors (Epoxomicin, Bortezomib, or MG132) or DMSO for 1, 2.5 and 5 hr assessing the levels of PSMA1-7 and Lamin B1, as a loading control. (D) Analysis of the Western blot time course as shown in C. Reported are Lamin B1-normalised PSMA1-7 protein level fold changes relative to T = 0 hr in response to treatment with DMSO, Epoxomycin, Bortexomib, or MG132. ns=p >0.05, one-way ANOVA and post-hoc Dunnett’s multiple comparisons test on the 5 hr time point, n=4, mean±SEM, FC=fold change. (E) UCSC genome browser view of the POMP locus (chr13: 28,658,045-28,661,232), displaying RefSeq genes (1), OMIM annotations (2), ENCODE candidate regulatory elements (cCREs) (3), H3K27Ac chromatin marks (4), and JASPAR transcription factor binding sites (5) tracks. In the JASPAR transcription factor binding sites (5) track HSF1/2 transcription factor binding sites are shown. (F) Western blot analysis of HEK293 cells treated with proteasome inhibitors (Epoxomicin, Bortezomib, MG132) or DMSO for 5 hr, probed for the heat shock response transcription factor HSF1, its phosphorylated active form (P-HSF1), their transcriptional target Hsp70 and ACTB, as loading control. Proteasome activity changes in response to proteasome inhibition was assayed by ABP in-gel fluorescence. (G) Analysis of experiments like the one shown in F. Treatment with proteasome inhibitors leads to a significant activation of HSF1, measured as an increase in P-HSF1, and its transcriptional downstream target Hsp70. *p≤0.05, **p<0.01, ***p<0.001, paired two-tailed t-tests, n=3 (P-HSF1) and 5 (Hsp70) biological replicates, mean±SD, FC=fold change. (H) Western blot analysis of HEK293 transfected with either one of two different siRNAs targeting HSF1 or scrambled control. The cells were analysed 72 hrs after transfection and blots were probed for HSF1, POMP and GAPDH, as loading control. Transfection of siRNAs leads to strong depletion of HSF1 and a reduction in POMP levels. (I) Analysis of the experiment shown in H. Transfection of HSF1 siRNAs leads to a significant reduction in HSF1 and POMP levels. ***p<0.001, ****p<0.0001, one-way ANOVA and post-hoc Dunnett’s test, n=4 biological replicates, mean±SD, FC=fold change. (J) Western blot analysis HEK293 cells treated simultaneously with KNK-437 and either one of three different proteasome inhibitors (Bortezomib, Epoxomicin, MG132) or DMSO for 5h. The samples were assayed for HSF1, P-HSF1, POMP and GAPDH, as loading control. Changes in proteasome activity were measured by in-gel ABP fluorescence. (K) Analysis of P-HSF1 levels from (H), normalized to total HSF1 and expressed as fold change (FC) relative to DMSO control. KNK-437 significantly reduces phosphorylation and activation of HSF1. **p<0.01, unpaired two-tailed t-test, n=3, mean±SD. (L) Analysis of experiments like the one shown in H. Simultaneous treatment with KNK-437 and proteasome inhibitors significantly reduces the POMP increase induced by proteasome inhibition. RM one-way ANOVA with post-hoc Dunnett’s test. n=7 (DMSO-PI) and 5 (KNK-437-PI), mean±SD, FC=fold change.

**Figure S3A.**
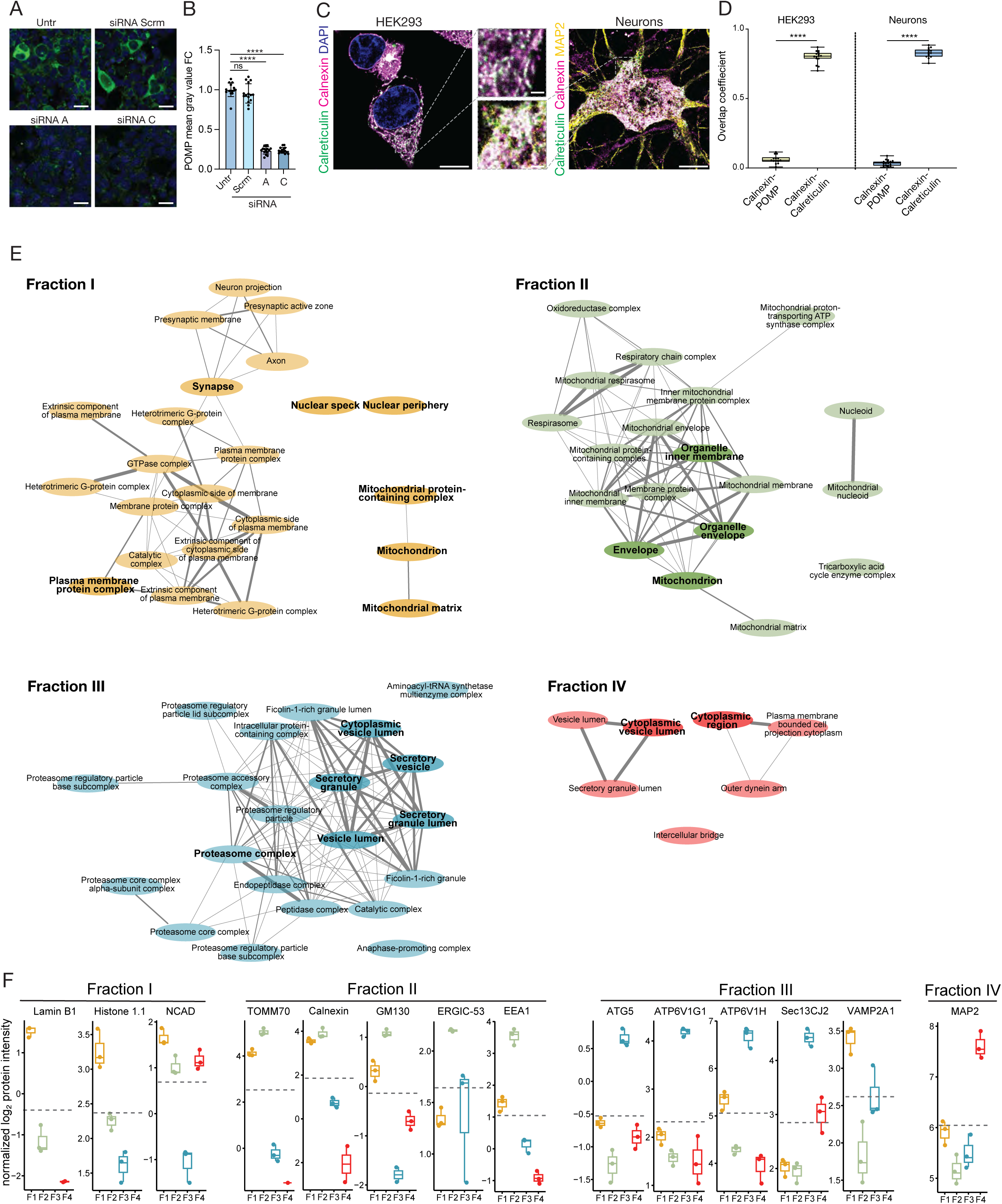
Methods validation and quality control experiments to localise POMP in cells. (A) Representative immunofluorescence images of cells either untransfected (Untr) or transfected with scrambled (Scrm) or two distinct siRNAs (A, C) against POMP. Cells were immunofluorescence stained for POMP (green) and nuclear marker DAPI (blue). Scale bars = 25 µm. (B) Analysis of images like the ones shown in A. POMP knock-down with validated siRNAs could be detected as a significant reduction in the POMP immunofluorescence signal, indicating that the antibody is suitable for this application. ****p < 0.0001, one-way ANOVA with post-hoc Dunnett’s test. n=15 fields of view, mean±SD, FC=fold change. (C) Representative confocal immunofluorescence image of HEK293 cells and primary neurons stained for the ER marker calnexin (magenta) and calreticulin (green), with the nuclear marker DAPI (blue) in HEK293 and the microtubule-associated protein MAP2 (yellow) in neurons. Scale bars = 10 and 1 µm. (D) Quantification of spatial colocalization (Mander’s overlap coefficient) of the ER marker calnexin with either calreticulin or POMP in HEK293 cells and neurons. Data show that while the two ER-resident proteins calnexin and calreticulin colocalise almost perfectly, POMP shows significantly lower Mander’s overlap coefficients in both cell types, suggesting limited localisation at the ER; ****p < 0.0001, unpaired two-tailed t-test, n=15 (Calnexin-POMP, HEK293), 16 (Calnexin-Calreticulin, HEK293), 20 (Calnexin-POMP, neurons), 15 (Calnexin-Calreticulin, neurons), boxplots show the median (line), interquartile range (box), and Min-Max whiskers. (E) Gene Ontology (GO) enrichment analysis of the top 100 proteins enriched in each subcellular fraction (I–IV) based on cellular components, as quantified by MS-based proteomic profiling. Networks show related GO terms clustered by functional similarity. In bold are the cellular compartments that best describe the different fractions. (F) Boxplots showing normalized log2 protein intensities of compartment-specific marker proteins across the four fractions (FI–IV). Marker proteins represent nuclear (Lamin B1, Histone 1.1), plasma membrane (NCAD), ER (Calnexin), ERGIC (ERGIC-53) Golgi (GM130), endosomes (EEA1) mitochondria (TOMM70), autophagy (ATG5, ATP6V1H, ATP6V1G1), secretory pathway (Sec13CJ2, VAMP2A1, ATP6V1H, ATP6V1G1), and cytosol (MAP2) compartments.

**Figure S3B.**
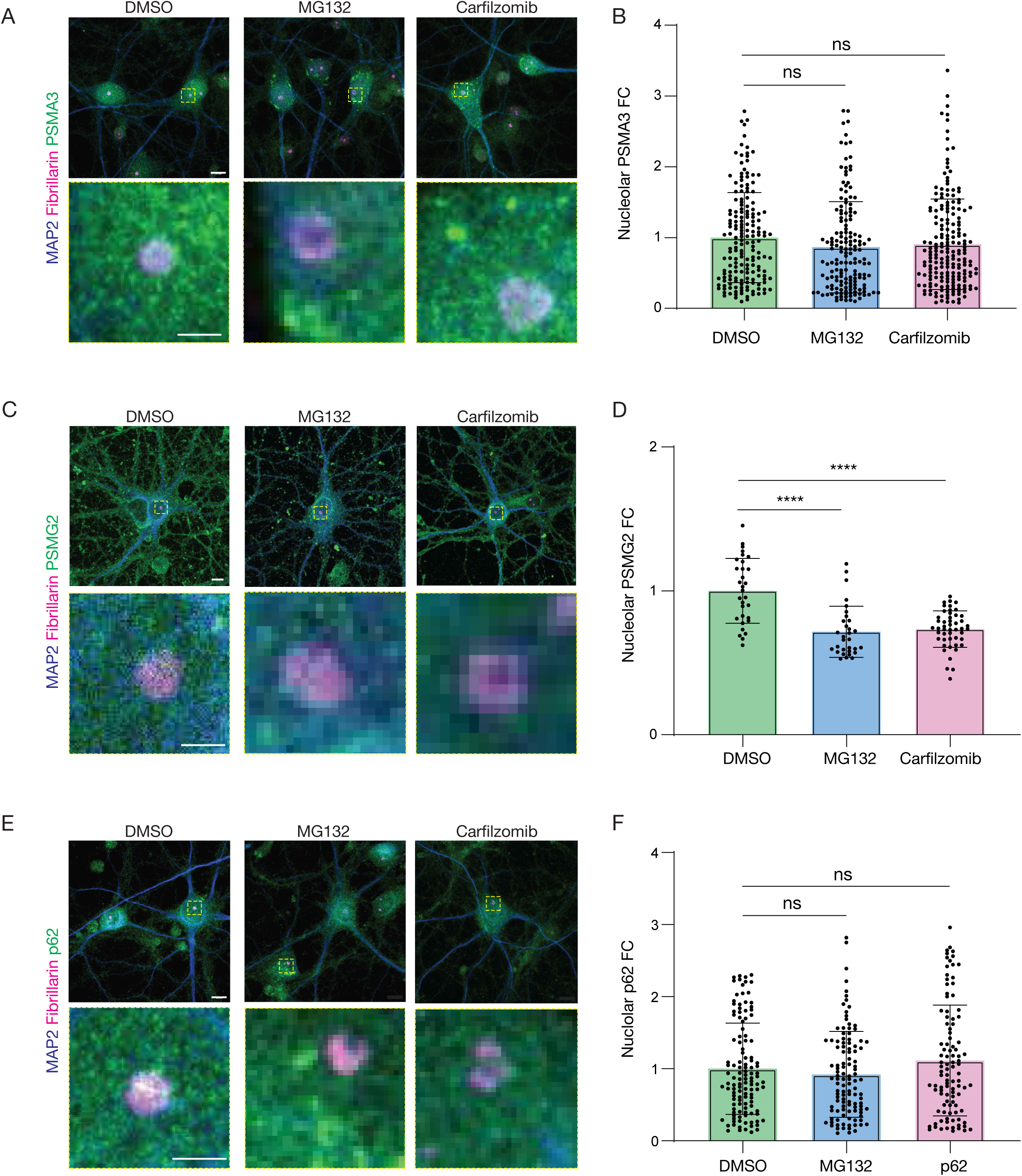
Nucleolar relocalisation of POMP is independent of the proteasome, its assembly intermediates and aggresome formation. (A) Representative immunofluorescence images of primary rat hippocampal neurons treated with either DMSO, MG132 or Carfilzomib for 5 hr and immunostained for the proteasome subunit PSMA3 (green), MAP2 (blue) and the nucleolar marker Fibrillarin (magenta). Insets below show higher magnification of the nucleolar region (dashed boxes). Scale bars = 10 (top panels) and 2 (bottom panels) µm. (B) Analysis of images like the ones shown in A. Proteasome inhibition does not lead to a significant change in the nucleolar fluorescence intensity for PSMA3 relative to the DMSO control. ns=p>0.05, one-way ANOVA and post-hoc Dunnett’s test, n= 172 (DMSO), 161 (MG132), 195 (Carfilzomib), mean±SD, FC= fold change. (C) Representative immunofluorescence images of primary rat hippocampal neurons treated with either DMSO, MG132 or Carfilzomib for 5 hr and immunostained for the proteasome CP assembly factor PSMG2 (green), MAP2 (blue) and the nucleolar marker Fibrillarin (magenta). Insets below show higher magnification of the nucleolar region (dashed boxes). Scale bars = 10 (top panels) and 2 (bottom panels) µm. (D) Analysis of images like the ones shown in C. Proteasome inhibition leads to a modest but significant reduction in the nucleolar fluorescence intensity for PSMG2 relative to the DMSO control. ****p<0.0001, one-way ANOVA and post-hoc Dunnett’s test, n= 31 (DMSO, MG132), 48 (Carfilzomib), mean±SD, FC= fold change. (E) Representative immunofluorescence images of primary rat hippocampal neurons treated with either DMSO, MG132 or Carfilzomib for 5 hr and immunostained for the autophagy receptor p62 (green), MAP2 (blue) and the nucleolar marker Fibrillarin (magenta). Insets below show higher magnification of the nucleolar region (dashed boxes). Scale bars = 10 (top panels) and 2 (bottom panels) µm. (F) Analysis of images like the ones shown in E. Proteasome inhibition does not lead to changes in the nucleolar levels of p62 compared to the DMSO control. ns=p>0.05, one-way ANOVA and post-hoc Dunnett’s test, n= 121 (DMSO), 113 (MG132), 102 (Carfilzomib), mean±SD, FC= fold change.

**Figure S4.**
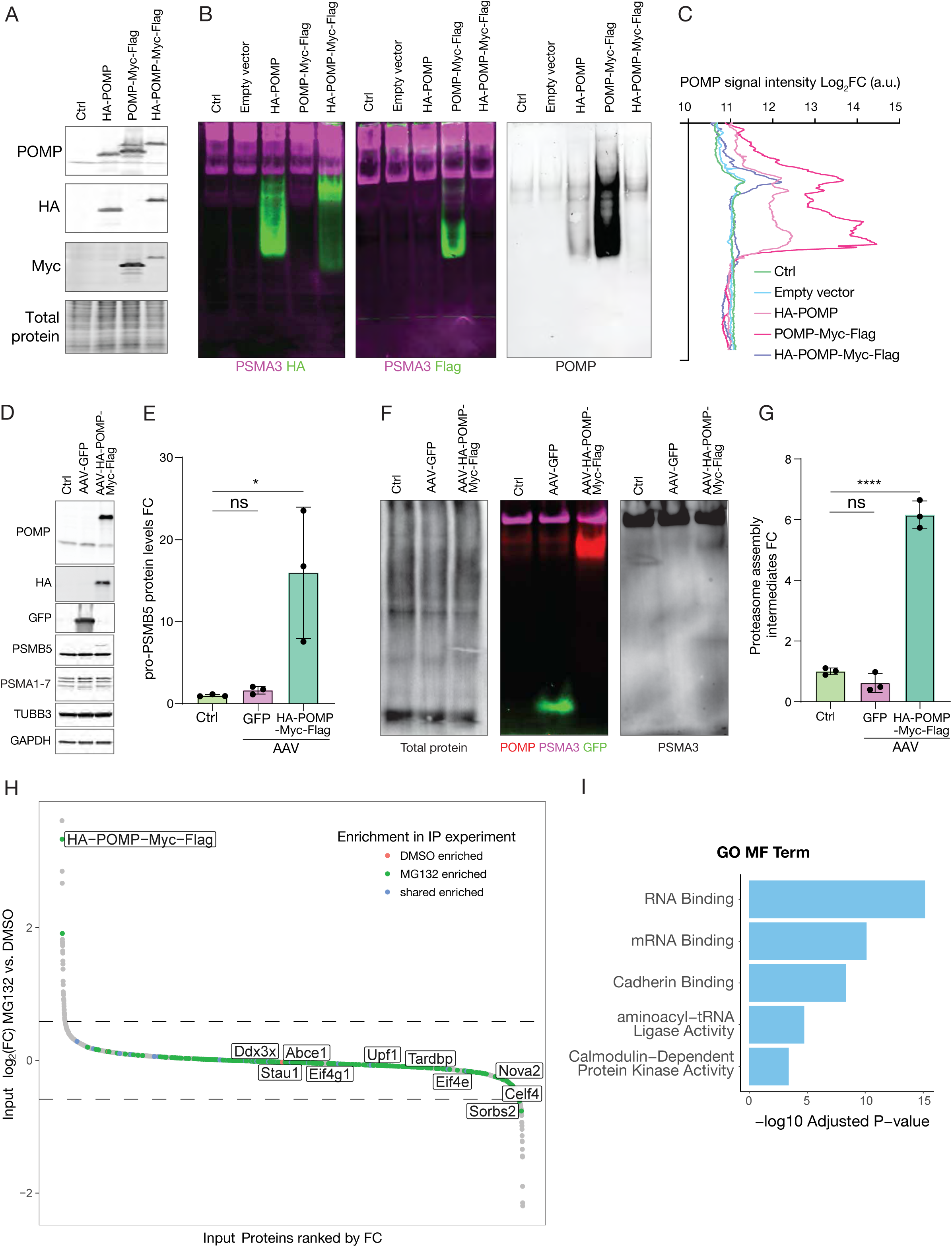
Establishment of a POMP bait construct for IP-MS analysis. (A) SDS-PAGE Western blot analysis of HEK293 cells expressing the test constructs: HA-POMP, POMP-Myc-Flag and HA-POMP-Myc-Flag. Blots were probed for POMP, HA, and Myc tags; total protein stain shown as a loading control. (B) Native PAGE Western blot analysis of HEK293 cells expressing the test constructs shown in A. Blots were probed for PSMA3 (magenta), HA (green), Flag (green) and POMP (grey). The different bait constructs show a different distribution across proteasome CP assembly intermediates. (C) Line plots of log-transformed fluorescence signal intensity profiles for the different POMP bait constructs along the gel lanes shown in B. While the HA-POMP and POMP-Myc-Flag constructs show higher expression levels but spurious integration across CP assembly intermediates, the HA-POMP-Myc-Flag construct shows a distribution pattern very similar to endogenous POMP. This suggests that HA-POMP-Myc-Flag integrates correctly into CP assembly intermediates and does not interfere with the CP assembly process. (D) SDS-PAGE Western blot analysis of primary rat cortical neurons, un-infected (Cntrl) or transduced with either AAV-GFP or AAV-HA-POMP-Myc-Flag. Blots show expression of POMP, HA, GFP, PSMB5, PSMA1-7, the neuronal marker TUBB3, and GAPDH, as loading control. POMP expression leads to an increase in the levels of pro-PSMB5. (E) Analysis of experiments like the one shown in D quantifying the levels of immature proteasome subunit (pro-PSMB5) following POMP overexpression. POMP overexpression leads to a significant increase in the levels of pro-PSMB5. ns=p>0.05, *p≤0.05, one-way ANOVA with post-hoc Dunnett’s multiple comparisons test, n= 3, mean±SD, FC=fold change. (F) Native PAGE Western blot analysis of neurons transduced as in D. A total protein stain is shown as loading control and the blots were probed for POMP, PSMA3 and GFP. While GFP does not interact with intracellular proteins and retains its expected size in native PAGE, POMP integrates into proteasome assembly intermediates, appearing as a high molecular weight band overlapping a PSMA3 band, corresponding to assembly intermediates. (G) Analysis of the levels of PSMA3^+^ proteasome assembly intermediates from experiments like the one shown in F. POMP overexpression in neurons leads to a significant increase in the levels of proteasome assembly intermediates. ns=p>0.05 ***p<0.001, one-way ANOVA and post-hoc Dunnett’s multiple comparisons test, n=3, mean±SD, FC=fold change. (H) Ranked abundance plot of quantified proteins in the input material used for IP-MS. Proteins are ranked by their log2 fold change (x-axis), with log2 fold change also shown on the y-axis. Selected proteins significantly enriched in the IP experiment are highlighted in color. Dashed lines indicate significance thresholds used to define differences in overall protein abundance between treatment conditions. (I) Gene ontology (GO) analysis of molecular function (MF) terms enriched among HA-POMP-Myc-Flag interactors. Top five significant terms shown, ranked by adjusted p-value (−log₁₀ scale).

**Figure S5.**
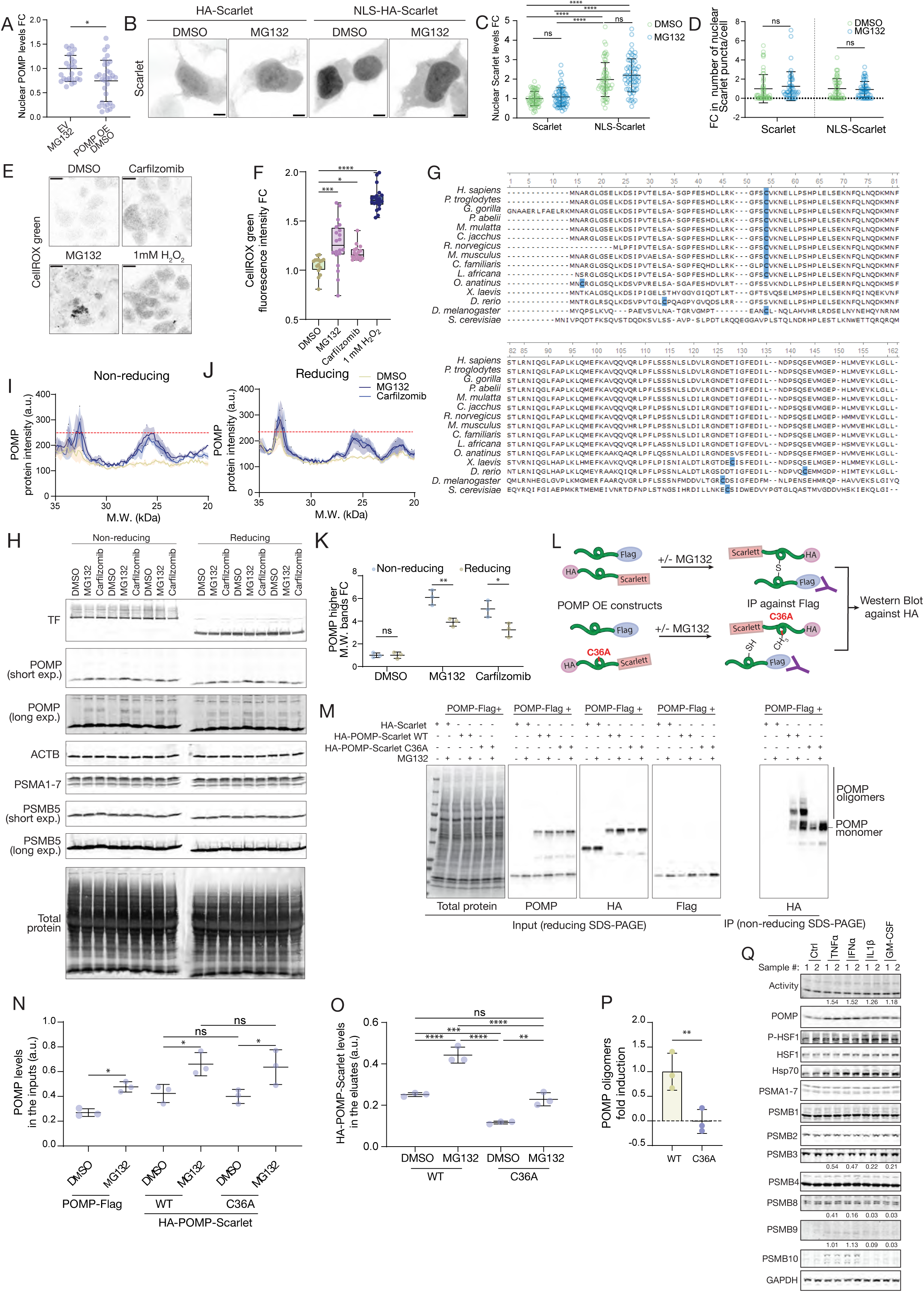
In response to cellular stress POMP forms oligomers and relocalizes to the nucleolus in a redox-sensitive manner. (A) Analysis of experiments like the one shown in Figure 5A. Quantification of nuclear POMP levels in HEK293 cells expressing either an empty vector control (EV) and treated with MG132 (5 hr) or expressing HA-POMP-Myc-Flag (POMP OE) and treated with DMSO (5 hr). POMP overexpression is not sufficient to elevate POMP in the nucleus to the same levels as endogenous POMP following MG132 treatment. *p≤0.05, unpaired two-tailed t-test, n=25 (EV MG132) and 31 (POMP OE DMSO), mean±SD, FC=fold change. (B) Representative images of HEK293 cells expressing HA-Scarlet (Scarlet) or NLS-HA-Scarlet (NLS-Scarlet) treated with DMSO or MG132 for 5 hr. Fusion of an NLS- to Scarlet leads to its strong nuclear accumulation but neither the expression levels or the staining pattern of the construct is affected by proteasome inhibition. Scale bars = 5 µm. (C) Analysis of the nuclear Scarlet levels in the HEK293 cells of experiments like the one shown in B. While NLS-fusion leads to a significant increase in the nuclear levels of Scarlet, MG132 treatment has no effect on the nuclear Scarlet levels for either construct. ns=p>0.05, ****p≤0.0001, one-way ANOVA and post-hoc Tukey’s multiple comparisons test, n=64 (Scarlet DMSO), 60 (NLS-Scarlet DMSO), 63 (Scarlet/NLS-Scarlet MG132) cells, mean±SD, FC=fold change. (D) Analysis of the number of nuclear Scarlet puncta in the HEK293 cells of the experiment shown in B. In general Scarlet does not form nuclear puncta and those few that form due to overexpression are not significantly affected by proteasome inhibition. ns=p>0.05, unpaired two-tailed t-test, n=53 (Scarlet DMSO), 54 (NLS-Scarlet DMSO), 46 (Scarlet MG132), 52 (NLS-Scarlet MG132) cells, mean±SD, FC=fold change. (E) Representative CellROX Green fluorescent images of HEK293 treated with either DMSO, the proteasome inhibitors MG132, Carfilzomib, or 1 mM H₂O₂, as a positive control. Proteasome inhibition and H₂O₂ treatments lead to an increase in CellROX Green fluorescence. Scale bars = 10 µm. (F) Analysis of experiments like the one shown in E. Proteasome inhibition and H₂O₂ treatment lead to a significant increase in CellROX Green fluorescence. *p≤0.05, ***p<0.001, ****p<0.0001, one-way ANOVA with post-hoc Dunnett’s multiple comparisons test, n=16 (DMSO), 17 (Carfilzomib), 20 (MG132, H₂O₂), boxplots show the median (line), interquartile range (box), and Min-Max whiskers. FC=fold change. (G) Multiple sequence alignment of POMP orthologs across species, highlighting conserved cysteine residues. (H) Reducing and non-reducing SDS-PAGE Western blot analysis of primary rat cortical neurons treated with either DMSO, MG132 or Carfilzomib for 5 hr. Blots were probed for proteasome components (PSMA1-7, PSMB5), POMP, ACTB and total protein (loading controls) and transferrin (TF) as a positive control for reduction. Proteasome inhibition leads to the appearance of oligomeric POMP bands, whose intensity decreases following reduction. (I) Line plot of the POMP oligomers intensity profiles under non-reducing conditions. The x-axis reports the molecular weight of the POMP^+^ bands and y-axis their intensity, the dashed line marks the maximal intensity of the peak at ∼25 kDa. Proteasome inhibition leads to the appearance of oligomeric bands. The lines and the areas represent mean and SEM, respectively. n=3 biological replicates. (J) Line plot of the POMP oligomers intensity profiles under reducing conditions. The x-axis reports the molecular weight of the POMP^+^ bands and y-axis their intensity, the dashed line marks the maximal intensity of the peak at ∼25 kDa under non-reducing conditions. The intensity of the peak at ∼25 kDa is decreased by incubation of the extracts with a reducing agent. The lines and the areas represent mean and SEM, respectively. n=3 biological replicates. (K) Analysis of the levels of POMP oligomeric species from experiments shown in H-J. Treatment of the extracts with a reducing agent leads to a significant reduction in the levels of the oligomeric POMP species induced by proteasome inhibition. *p≤0.05, **p<0.01, unpaired two-tailed t-tests, n=3 biological replicates, FC= fold change. (L) Schematic of the POMP oligomerization assay. HA-tagged WT POMP-Scarlet or C36A POMP-Scarlet constructs were co-expressed with a Flag-tagged POMP construct in HEK293 cells that were treated with either DMSO or MG132 for 5 hr. Lysates were subjected to Flag co-IP and the eluates were analysed by non-reducing SDS-PAGE and Western blotting for the HA tag to assay for oligomerization via covalent and non-covalent interactions. (M) Western blot analysis of input and eluate samples from the experiment in L. While WT HA-POMP-Scarlet can interact with POMP-Flag both via non-covalent interactions and disulfide bonds formed via the Cys-residue, the C36A mutant can only interact with the bait via non-covalent interactions and the ability to form higher molecular weight oligomers is entirely lost. Blots of the eluates show that the different constructs express to the same levels and the differences seen after IP cannot be explained by the inputs. (N) Analysis of the POMP levels in the inputs used for the co-IP experiments like the one shown in M. Treatment with MG132 leads to a similar increase in the levels of all three POMP constructs. The C36A mutation does not have any adverse effect on POMP expression. ns=p>0.05, *p≤0.05, one-way ANOVA with post-hoc Šidák’s multiple comparisons tests, n=3 biological replicates, mean±SD. (O) Analysis of HA-POMP-Scarlet levels in the eluates of experiments like the one shown in M. C36A mutation prevents formation of oligomers via disulfide bond formation and leads to a significant reduction in the levels of oligomers formed in response to MG132 treatment. However, MG132 treatment is able to induce a significant increase in the levels of C36A POMP in the eluates via increased non covalent interactions. ns=p>0.05, **p<0.01, ***p<0.001, ****p<0.0001, one-way ANOVA with post-hoc Tukey’s multiple comparisons test, n=3 biological replicates, mean±SD. (P) Analysis of the POMP oligomers formed in response to MG132 treatment in the eluates of experiments like the one shown in M. C36A mutation prevents formation of POMP-Scarlet oligomers in the Flag co-IP eluates. **p<0.01, unpaired two-tailed t-test, n=3 biological replicates, mean±SD. (Q) Western blot analysis of HEK293 cells treated with pro-inflammatory cytokines TNFα, INFα, IL1β, GM-CSF for four days. Proteasome activity was probed by in-ge ABP fluorescence. Blots were probed for POMP, HSF1 and its activated form P-HSF1, Hsp70, proteasome subunits (PSMA1-7, PSMB1, PSMB2, PSMB4), immunoproteasome subunits (PSMB8, PSMB9, PSMB10) and GAPDH, as loading control. The numbers reported above the POMP, PSMB4 (pro-form), PSMB9 and PSMB10 represent the log_2_FC_average_ relative to Cntrl for the different treatments. n=4 biological replicates. Quantifications are reported only where at least in one of the treatments log_2_FC_average_>0.4. For POMP, P-HSF1 and Hsp70 the quantifications are reported in Figure 5.

**Figure S6A.**
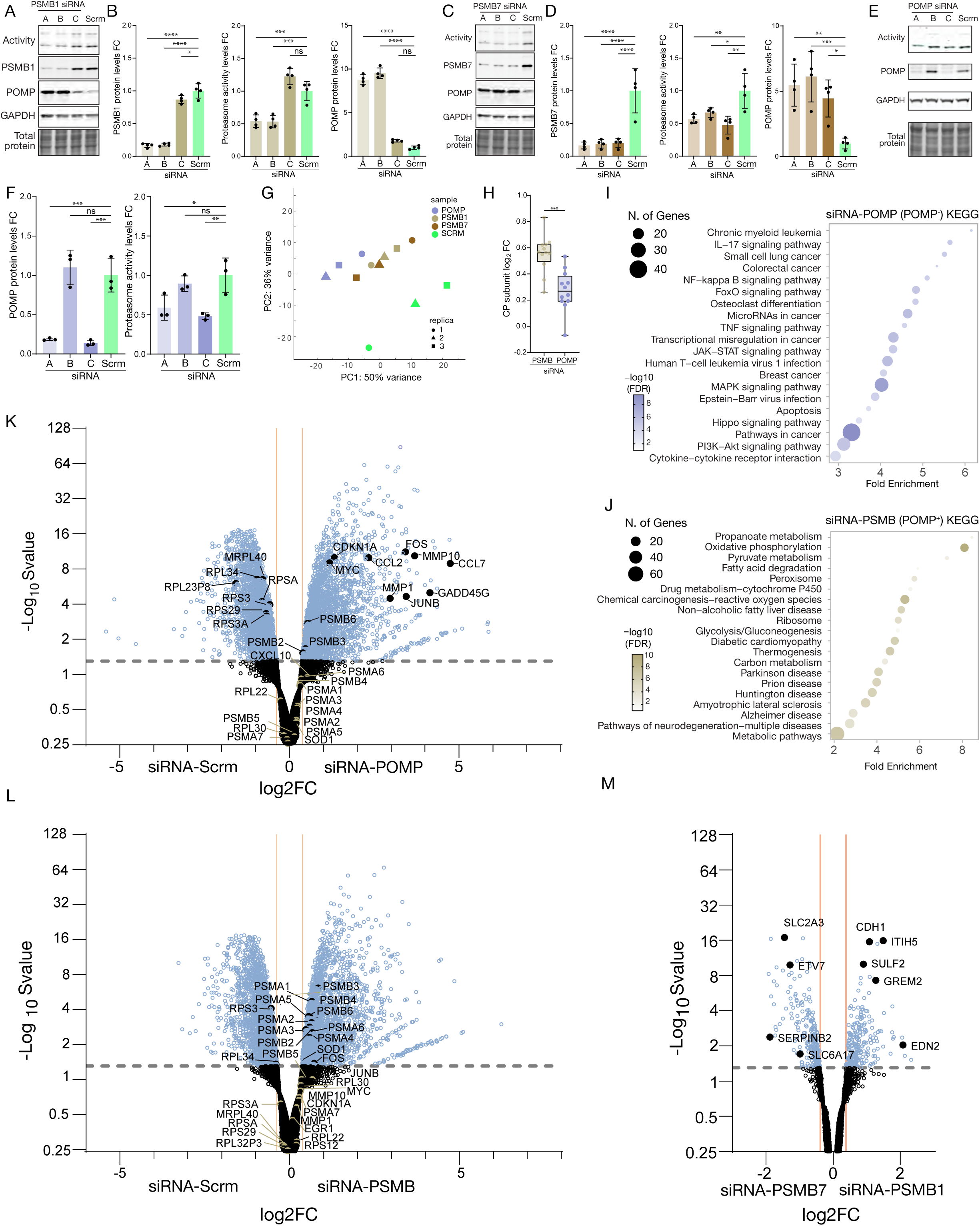
POMP and proteasome subunit depletion lead to distinct transcriptional responses. (A) Proteasome activity measured as in-gel ABP fluorescence and Western blot analysis of PSMB1, POMP and GAPDH levels in HEK293 cells transfected with each of three different siRNAs targeting PSMB1 or scrambled control (Scrm) and cultured for 72 hr before analysis. Total protein staining was used as a loading control. (B) Analysis of experiments like the one shown in A. Quantification of PSMB1 protein levels shows significant depletion with all three siRNAs, with siRNA-A and siRNA-B producing the strongest effects. Quantification of proteasome activity by in-gel ABP fluorescence indicates significantly reduced activity following PSMB1 knockdown. POMP protein levels are significantly increased in response to PSMB1 depletion. ns=p>0.05, *p≤0.05, ***p<0.001, ****p<0.0001, one-way ANOVA with post-hoc Dunnett’s test, n = 4 biological replicates, mean±SD, FC=fold change. (C) Proteasome activity measured as in-gel ABP fluorescence and Western blot analysis of PSMB7, POMP and GAPDH levels in HEK293 cells transfected with each of three different siRNAs targeting PSMB7 or scrambled control (Scrm) and cultured for 72 hr before analysis. Total protein staining was used as a loading control. (D) Analysis of experiments like the one shown in C. Quantification of PSMB7 protein levels shows significant depletion with all three siRNAs. Quantification of proteasome activity by in-gel ABP fluorescence indicates significantly reduced activity following PSMB7 knockdown. POMP protein levels are significantly increased in response to PSMB7 depletion. *p≤0.05, **p<0.01, ***p<0.001, ****p<0.0001, one-way ANOVA with post-hoc Dunnett’s test, n = 4 biological replicates, mean±SD, FC=fold change. (E) Proteasome activity measured as in-gel ABP fluorescence and Western blot analysis of POMP and GAPDH levels in HEK293 cells transfected with each of three different siRNAs targeting POMP or scrambled control (Scrm) and cultured for 72 hr before analysis. Total protein staining was used as a loading control. (F) Analysis of experiments like the one shown in E. Quantification of POMP protein levels shows a significant depletion with siRNA A and C. Quantification of proteasome activity by in-gel ABP fluorescence indicates significantly reduced activity following POMP knockdown. ns=p>0.05, *p≤0.05, **p<0.01, ***p<0.001, one-way ANOVA with post-hoc Dunnett’s test, n = 4 biological replicates, mean±SD, FC=fold change. (G) Principal component analysis (PCA) of transcriptomic data from cells transfected with siRNAs targeting POMP, PSMB1, PSMB7 or scrambled control. Samples of all three knock-down conditions cluster away from scrambled, consistent with the fact that all three result in proteasome inhibition. However, while PSMB1 and PSMB7 are interspersed with one another and cannot be clustered apart, POMP samples form a distinct cluster, suggesting that POMP knock-down gives rise to a distinct transcriptional signature. (H) Analysis of the induction of CP subunits’ mRNAs (Log_2_FC, vs Scrm) following PSMB (POMP^+^) and POMP (POMP^-^) knock-down. Data show a significant reduction in CP transcript levels upon POMP depletion, suggesting that POMP is required to facilitate expression of CP mRNAs. ***p<0.001, unpaired two-tailed t-test, n=14, each data point representing a different CP subunit. Boxplots show the median (line), interquartile range (box), and Min-Max whiskers. (I) KEGG pathway enrichment analysis of differentially expressed genes upon POMP knockdown (POMP^-^). Dot size corresponds to the number of genes per pathway; color indicates adjusted p-value (FDR). Pathways are ranked based on fold enrichment. The POMP-specific transcriptional signature is characterised by the activation of pro-inflammatory and cancer-related pathways. (J) KEGG pathway enrichment analysis of differentially expressed genes upon PSMB knockdown (POMP^+^). Dot size corresponds to the number of genes per pathway; color indicates adjusted p-value (FDR). Pathways are ranked based on fold enrichment. The PSMB-specific transcriptional signature (POMP is characterised by a rewiring of cellular metabolism, ribosome biogenesis and a neurodegeneration-like transcriptional signature. (K) Volcano plots of RNA-seq differential expression analysis comparing siRNA-POMP (POMP^-^) to scrambled control. Significantly differentially regulated genes are highlighted in blue. Following POMP knock-down cells downregulate the expression of ribosomal genes, are unable to mount robust compensatory expression of proteasome CP transcripts and strongly induce proinflammatory factors, proto-oncogenes and cell cycle inhibitors. The dashed horizontal and vertical lines indicate the thresholds used for differential gene expression. (L) Volcano plots of RNA-seq differential expression analysis comparing siRNA-PSMB (POMP^+^) to scrambled control. Significantly differentially regulated genes are highlighted in blue. Following PSMB knock-down cells are able to maintain expression of ribosomal genes at control levels, induce compensatory expression of proteasome CP transcripts and prevent induction of proinflammatory factors, proto-oncogenes and cell cycle inhibitors. The dashed horizontal and vertical lines indicate the thresholds used for differential gene expression. (M) Volcano plot comparing differential gene expression between PSMB7 and PSMB1 knock-downs. Significantly regulated genes are shown in blue. Only few genes are differentially regulated and no clear transcriptional signature emerges from this comparison. The dashed horizontal and vertical lines indicate the thresholds used for differential gene expression.

**Figure S6B.**
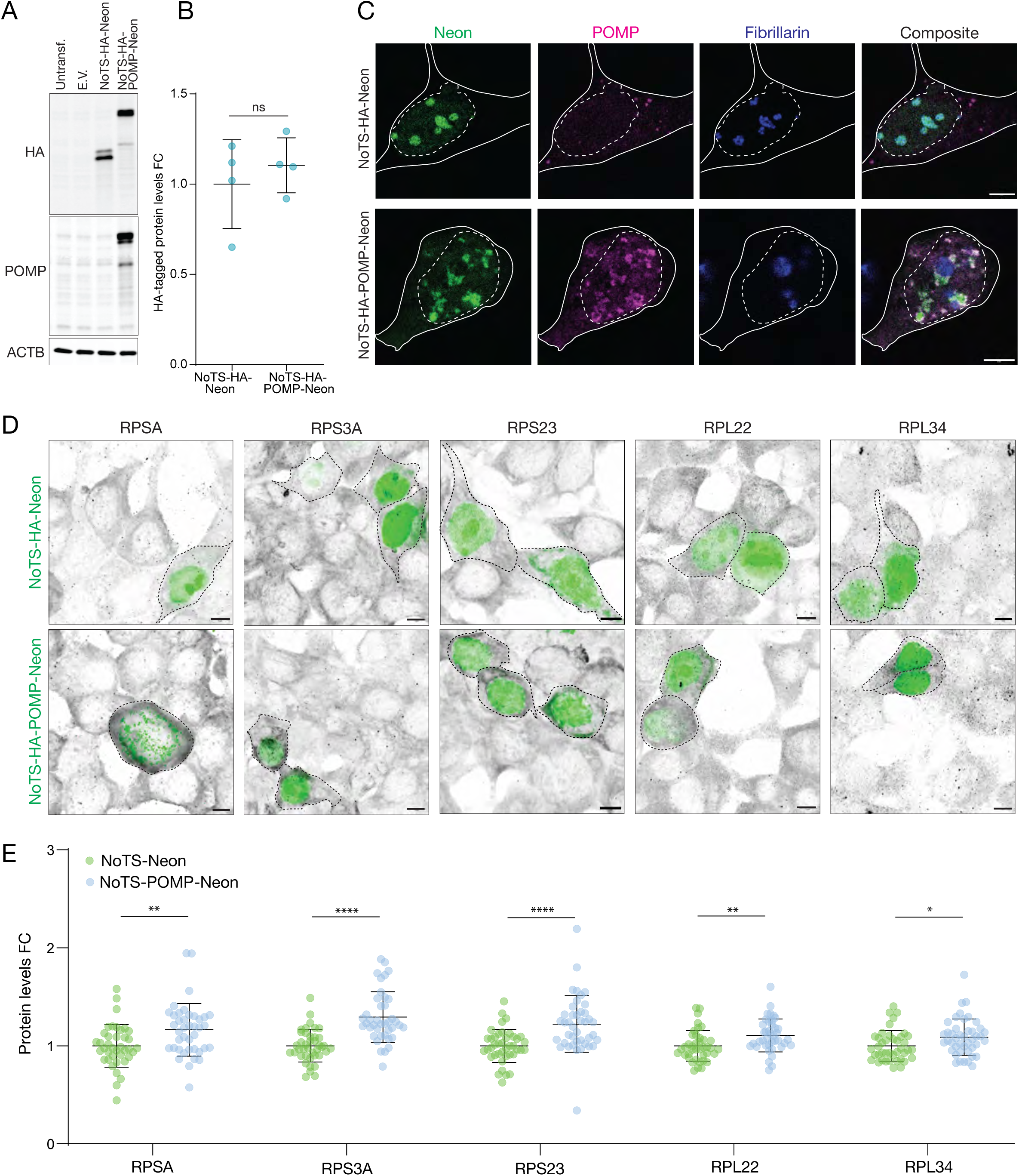
Enforcing nucleolar localisation of POMP drives ribosomal subunit synthesis and affects splicing. (A) Representative Western blot of untransfected HEK293 cells and HEK293 expressing either an empty vector control construct (E.V.), NoTS-HA-Neon or NoTS-HA-POMP-Neon. The blots were probed for HA-tag, POMP and ACTB as loading control. (B) Analysis of experiments like the one shown in A. Construct transfection conditions were optimised to obtain the same levels of transgene expression. ns=p>0.05, unpaired two-tailed t-test, n=4, mean±SD, FC=fold change. (C) Representative immunofluorescence images showing subcellular localization of the NoTS-HA-Neon and NoTS-HA-POMP-Neon constructs in HEK293 cells. The cells were stained for Neon (green), POMP (magenta) and the nucleolar marker Fibrillarin (blue). Dotted lines mark the nuclear boundaries and the solid lines mark the cell boundaries. Overexpression of NoTS-HA-POMP-Neon leads to accumulation of POMO in the nucleolus. Scale bars=5 μm. (D) Immunofluorescence images of HEK293 cells expressing either NoTS-HA-Neon or NoTS-HA-POMP-Neon for 72 hrs and stained for the ribosomal proteins (RPSA, RPS3A, RPS23, RPL22, RPL34). Neon is shown in green and ribosomal proteins in grey. Overexpression of NoTS-HA-POMP-Neon leads to increased labelling for the ribosomal subunits assayed. (E) Analysis of immunofluorescence images like those shown in D. Quantification of ribosomal protein levels cells expressing either NoTS-HA-Neon or NoTS-HA-POMP-Neon. POMP expression significantly increases the cellular levels of the ribosomal proteins assayed. *p≤0.05, **p<0.01, ****p<0.0001, unpaired two-tailed t-tests, mean ± SD, FC=fold change.

**Figure S6C.**
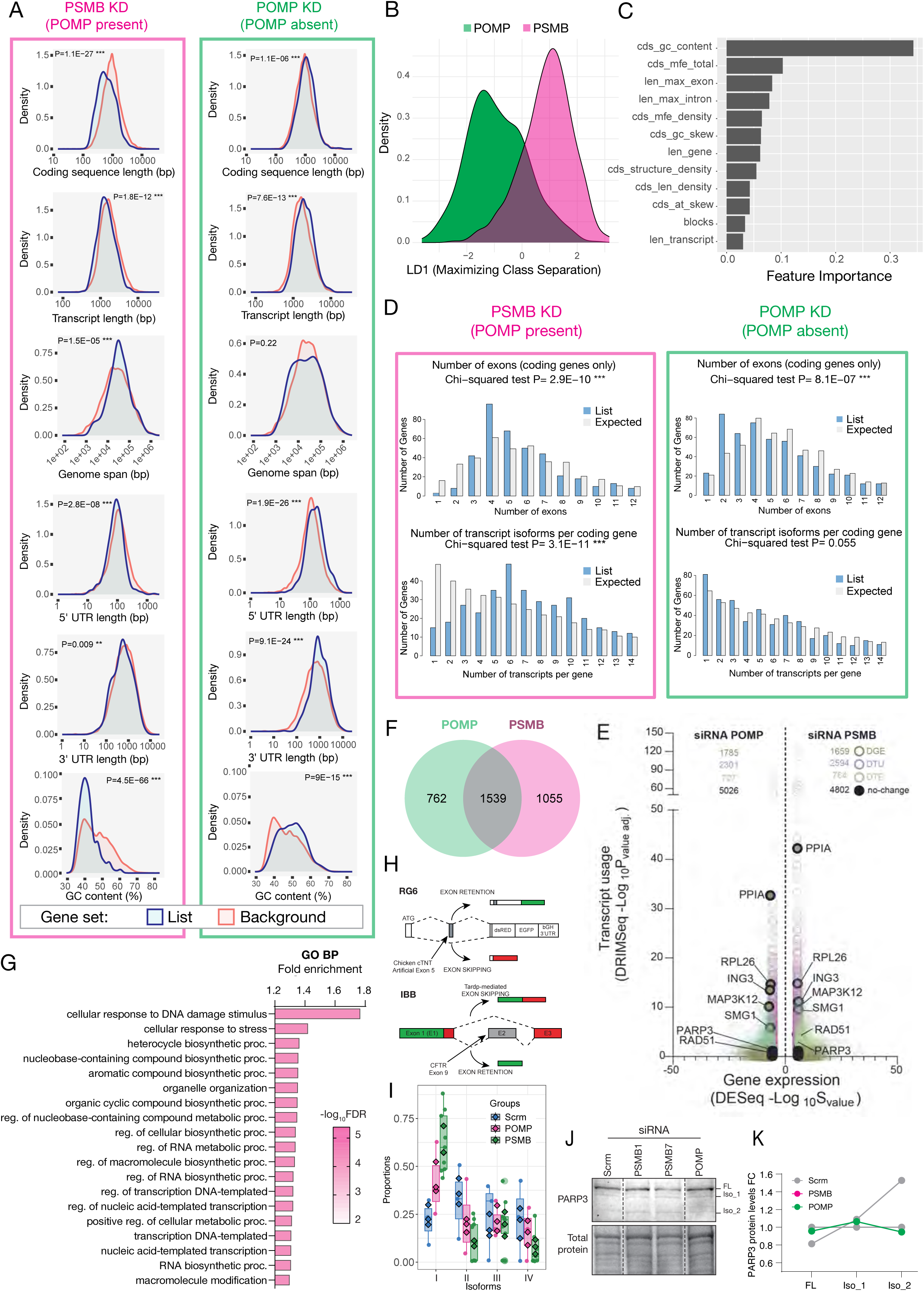
Transcriptomic features associated with POMP-dependent and POMP-independent regulation upon proteasome disruption. (A) Density plots comparing transcriptomic features of differentially regulated genes (blue, “List”) to background gene sets (red, “Background”) in PSMB KD (POMP present, left) and POMP KD (POMP absent, right) conditions. Features include coding sequence length, transcript length, genome span, UTR lengths, and GC content. (B) Linear discriminant analysis (LDA) based on sequence and structural features shows partial separation between gene sets regulated upon PSMB KD (magenta) and POMP KD (green). (C) Feature importance plot from a random forest classifier trained to distinguish between POMP-dependent and POMP-independent gene sets. Top contributing features include CDS GC content, minimum free energy (mfe), exon/intron lengths, and sequence skew metrics. (D) Feature enrichment analysis histograms showing distributions of exon counts (top panels) and transcript isoforms per coding gene (bottom panels) in PSMB KD (left, pink) and POMP KD (right, green) compared to genome-wide expectations (light blue). Chi-squared p-values indicate deviation from expected distributions. (E) Juxtaposed transcript usage over gene expression plots for POMP KD (left) and PSMB KD (right). The x-axis reports differential gene expression and the y-axis differential transcript usage. Genes showing differential gene expression (DGE), differential transcript usage (DTU), or both (DTE) are color-coded; selected genes that show differential transcript usage regulation and are involved in DNA-repair and stress response are labeled. (F) Venn diagram depicting the overlap between genes that, by DRIM-Seq analysis, show differential transcript usage in POMP KD and PSMB KD conditions. (G) Gene ontology (GO) enrichment analysis of biological process (BP) terms for genes that show differential transcript usage uniquely in the POMP-dependent condition (PSMB KD only,1055 genes). Top terms are related to DNA damage response and cellular stress response. (H) Schematic diagrams of splicing reporters used to monitor exon retention vs. exon skipping. Upper one is the RG6 general splicing reporter, which incorporates an artificial version of the chicken cardiac troponin (cTNT) exon 5 and is not selective for specific splicing factor. Lower one is the IBB reporter, which contains exon 9 from the CFTR gene and is selective for TDP-43-mediated splicing events. The two reporters work in the way that exon retention/skipping events will lead to different ratios of GFP:RFP signal. (I) DRIMseq analysis of differential isoform usage for PARP3 in HEK293 cells transfected with siRNAs against PSMB1, PSMB7, POMP and scrambled control (Scrm) for 72 hr. n=3 (Scrm), 6 (PSMB1, PSMB7 and POMP). Boxplots show the median (line), interquartile range (box), and Min-Max whiskers. (J) Western blot analysis of PARP3 isoform usage in HEK293 cells transfected with siRNAs against PSMB1, PSMB7, POMP and scrambled control for 72 hr. Knock-down of proteasome subunits, but not POMP, leads to an increase in the levels of one of the lower molecular weight isoforms of PARP3 (indicated as iso_1 and _2). Total protein stain is reported and was used as loading control. The dashed lines mark where the gel was spliced. (K) Quantifications of the relative abundance of PARP3 FL and its two isoforms in experiments like the one shown in I. PSMB1 and PSMB7 knock-down data were merged into a common PSMB term. The data show that in response to proteasome subunit knock-down expression of PARP3 FL decreases slightly in favour of its alternative isoforms, in particular iso_2. By contrast, POMP knock-down leads to a reduction in the levels of the two isoforms compared to scrambled control. n=2 (Scrm, POMP) and 4 (PSMB) biological replicates.

**Figure S7.**
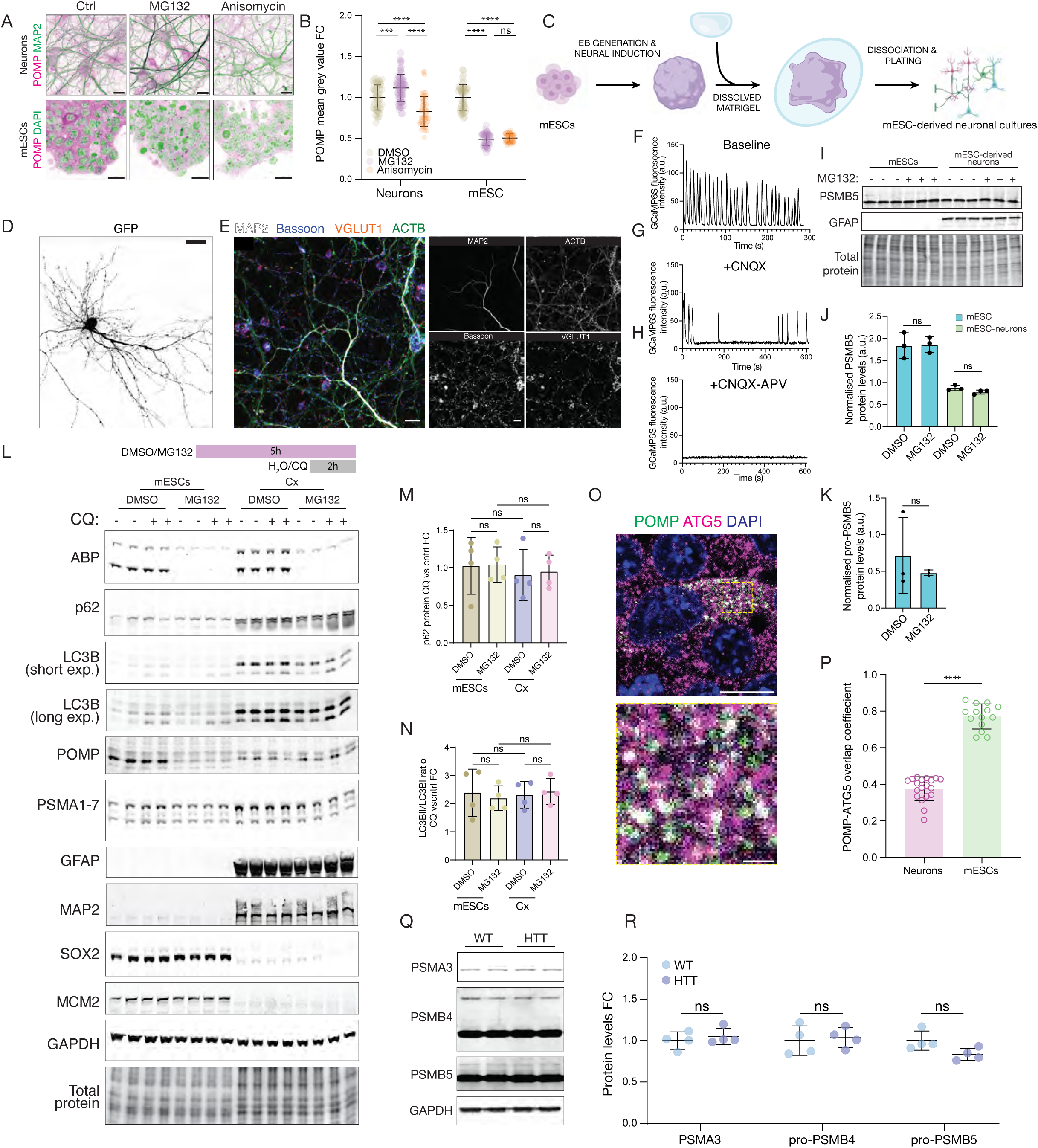
POMP levels dynamics and autophagic regulation in neurons and mESCs under proteostatic stress. (A) Representative confocal images of primary rat hippocampal neurons and mESCs treated with either DMSO (control), MG132 (proteasome inhibitor), or anisomycin (translation inhibitor) for 5 hr and stained for the proteasome maturation protein POMP (green) and DAPI/MAP2 (purple). Scale bars=10 µm. (B) Analysis of images like the ones shown in A. Quantification of POMP protein levels under the indicated treatments in neurons and mESCs. While in neurons proteasome inhibition leads to a significant accumulation of POMP and protein synthesis inhibition to a significant reduction, in mESCs both treatments lead to a significant reduction. ns=p>0.05, ***p<0.001, ****p<0.0001, one-way ANOVA with post-hoc Tukey’s multiple comparisons test, n= 49 (mESC DMSO and MG132), 50 (neurons DMSO and Anisomycin, mESC Anisomycin), 75 (neurons MG132) fields of view, mean±SD. (C) Schematic representation of the mESC-to-neuron differentiation using an unguided protocol and dissociation into cultures at early/mid-neurogenesis stage. (D) Representative image of a GFP-labeled mESC-derived neuron for morphological reference. Scale bar=20 µm. (E) Representative immunofluorescence images of mESC-derived neurons showing neuronal co-staining for the dendritic marker MAP2 (grey), the synaptic vesicle marker VGLUT1 (red), the presynaptic scaffold protein Bassoon (blue), and actin (ACTB, green). On the right in grey are the single channel images of the individual markers. Scale bars=10 µm. (F–H) Representative traces from calcium imaging of mESC-derived neurons under baseline (F), CNQX (AMPA/kainate receptor antagonist) (G), or CNQX + APV (NMDA receptor antagonist) (H) conditions. The x-axis reports time (s) and the y-axis the GCaMP6f fluorescence (a.u.). The traces show that mESC-derived neurons are active and synaptic activity is suppressed by glutamate receptor antagonists. (I) Western blot analysis of mESCs and mESC-derived neurons treated with DMSO or MG132 for 5 hr. Blots were probed for PSMB5 and the astrocytic marker GFAP. Total protein loading shown as control. (J) Analysis of the PSMB5 levels in the experiment shown in I. Proteasome inhibition (MG132, 5 hr) in mESCs and mESC-derived neurons does not lead to significant changes in the levels of mature PSMB5. ns=p>0.05, unpaired two-tailed tests, n=3, mean±SD. (K) Analysis of the pro-PSMB5 (immature PSMB5) levels in the mESC samples shown in I. Proteasome inhibition (MG132, 5 hr) in mESCs does not lead to significant changes in the levels of pro-PSMB5. pro-PSMB5 was not measured in mESC-derived neurons because it could not be detected. ns=p>0.05, unpaired two-tailed tests, n=3, mean±SD. (L) Representative Western blot analysis of mESCs and primary rat cortical neurons (Cx) treated with DMSO or MG132 for three hours prior to addition of either chloroquine (CQ, autophagy inhibitor) or water (ctrl), as control, and incubation for a further two hours to assess changes in bulk autophagic flux in response to proteasome inhibition. Proteasome activity reduction was monitored by in-gel ABP fluorescence. The blots were probed for the mESC markers SOX2 and MCM2, the neuronal marker MAP2, the astrocytic marker GFAP, the proteasome subunits PSMA1-7, POMP and the autophagy proteins LC3B and p62. (M) Analysis of the changes in autophagic flux in mESCs and primary rat cortical neurons (Cx) in response to proteasome inhibition. In the two cell types, the fold increase in the levels of autophagy receptor p62 in response to chloroquine (CQ) compared to control (water) was calculated under both baseline conditions and during proteasome inhibition. Proteasome inhibition did not lead to a significant change in autophagy flux in either of the cell types tested over the course of the experimental manipulations. ns=p>0.05, one-way ANOVA with Tukey’s post-hoc test, n=4 biological replicates, mean±SD. a.u.=arbitrary units. (N) Analysis of the changes in autophagic flux in mESCs and primary rat cortical neurons (Cx) in response to proteasome inhibition. In the two cell types, the fold change in the ratio of LC3B-II/LC3B-I was calculated between CQ and control (water) under DMSO and MG132 treatment conditions. CQ led to a noticeable accumulation of LC3B-II under both DMSO and MG132 conditions; however, the ratio between CQ and its corresponding control in response to proteasome inhibition or control treatment did not change significantly. This suggests that proteasome inhibition did not lead to a significant change in autophagy flux in either of the cell types tested over the course of the experimental manipulations. ns=p>0.05, one-way ANOVA with Tukey’s post-hoc test, n=4 biological replicates, mean±SDa.u.=arbitrary units. (O) Representative immunofluorescence images of mESCs stained for POMP (green), DAPI (blue) and the pre-autophagosome marker ATG5 (magenta). The dashed yellow box indicates the region shown in the magnified inset below. The images show extensive overlap between POMP and ATG5. Scale bars=10 and 1 µm. (P) Quantification of POMP-ATG5 colocalization coefficient (Mander’s overlap coefficient) in primary rat hippocampal neurons and mESCs. In mESCs the extent of POMP-ATG5 colocalization is significantly higher than in neurons. ****p<0.0001, unpaired two-tailed t-test. n=19 (neurons) and 14 (mESCs) cells, mean±SD. (Q) Western blot analysis of hippocampi from WT and mice expressing mutant huntingtin (HTT). The blots were probed for the proteasomal subunits (PSMA3, PSMB4 and PSMB5) and GAPDH, as loading control. (R) Analysis of experiments like the one shown in Q. Quantification of PSMA3, pro-PSMB4 and pro-PSMB5 protein levels. The data show that the levels of proteasome subunits and their precursors are unaffected by HTT pathology. ns=p>0.05, unpaired two-tailed t-tests, n=4 biological replicates.

